# Fluphenazine, an antipsychotic compound, ameliorates Alzheimer’s disease by clearing amyloid beta accumulation in *C. elegans*

**DOI:** 10.1101/2025.04.24.650484

**Authors:** Mayur Nimbadas Devare, Victoria Le, Vanessa Chung, TJ Vu, Matt Kaeberlein

## Abstract

The ageing population worldwide faces an increasing burden of age-related conditions, with Alzheimer’s disease being a prominent neurodegenerative concern. Drug repurposing, the practice of identifying new therapeutic applications for existing drugs, offers a promising avenue for accelerated intervention. In this study, we utilized the yeast *S. cerevisiae* to screen a library of 1,760 FDA-approved compounds, both with and without rapamycin, to assess potential synergistic effects on yeast growth. We identified 87 compounds that showed synergistic effect with rapamycin and caused growth defects in yeast. These compounds were further screened for their effects on paralysis in a *C. elegans* model of Alzheimer’s disease. We found that three compounds synergistically delayed paralysis in combination with rapamycin. Additionally, four other compounds delayed paralysis when tested at different concentrations. Moreover, we tested fluphenazine, an antipsychotic drug identified in our screen and found that it enhanced the overall health of treated worms. Western blot and X-34 staining confirmed that fluphenazine reduced amyloid-beta accumulation. These results indicate that repurposed drugs have significant potential to accelerate Alzheimer’s disease drug discovery.

## Introduction

In the realm of medical science, the pursuit of effective interventions to ameliorate aging and combat Alzheimer’s disease (AD) has fueled a dynamic landscape of research and innovation [1]. One promising avenue that has gained significant attention is “drug repurposing” – the exploration of existing pharmaceuticals for new therapeutic purposes [2]. This approach not only accelerates the drug development process but also holds the potential to revolutionize the treatment paradigms for aging-related conditions, offering new hope for individuals grappling with the challenges of advancing age and cognitive decline [3]. As the global population ages and the prevalence of Alzheimer’s disease continues to rise, the urgency to discover safe and effective interventions has never been more pressing [4, 5].

Yeast has been used extensively for high throughput screening of thousands of compounds for drug discovery primarily due to its high level of conservation in cellular processes when compared to human cells, as well as its ease of manipulation.[6]. Similarly in recent years *C. elegans* has emerged as a powerful and unique model for studying aging and AD because of its well-defined nervous system, short lifespan, and genetic manipulability [7]. Engineered transgenic *C. elegans* worms have been designed to express Aβ peptides, simulating the pathological characteristics of AD. *C. elegans* strain GMC101, expresses full length human amyloid beta (Aβ1–42) peptide product in muscle in a temperature dependent manner and results in worm paralysis and death [8] and offers valuable insights into key aspects of AD pathology [9-11].

Nematode and vertebrate neurons exhibit remarkable similarities at the molecular and cellular levels. For example, ion channels, receptors, classic neurotransmitters, vesicular transporters, and the neurotransmitter release machinery have comparable structures and functions in both vertebrates and *C. elegans* [12, 13].

Notably, many of these pathways have been shown to contribute to early stages of AD [14]. Despite this, there has been a noticeable gap in systematic drug development explicitly directed toward AD. Moreover, there exists the potential for undiscovered metabolic pathways, not conventionally linked with AD, that might respond positively to pharmacological intervention using existing approved drugs. This unexplored terrain presents an intriguing opportunity to uncover novel therapeutic approaches aimed at extending healthy aging and ameliorating associated diseases. It underscores the need for a more comprehensive exploration of existing drugs and their potential impacts on the intricate metabolic networks associated with AD.

The mammalian target of rapamycin (mTOR) is a serine/threonine multidomain protein that contains a kinase domain and an FKBP12-binding domain. Rapamycin binds to the FK506-binding protein 12 (FKBP12) and interacts with the FKBP12-rapamycin binding (FRB) domain of mTOR, leading to the inhibition of mTORC1 activity [15]. mTOR plays a crucial role in regulating various physiological processes by integrating signals from upstream regulators, including insulin, growth factors, AMPK, PI3K/Akt, and glycogen synthase kinase 3 (GSK-3) [16]. Dysregulation of mTOR has been increasingly implicated in aging [17, 18], as well as numerous disease including cancer [19, 20]), diabetes [21], obesity [22], cardiovascular disease [23], and neurodegenerative disorders [16, 24]. Notably, substantial evidence links mTOR activation to the progression of Alzheimer’s disease (AD), highlighting its intersection with AD pathology and clinical symptoms [16]. mTOR signaling is strongly associated with key hallmarks of AD, including amyloid-beta (Aβ) plaques, neurofibrillary tangles (NFTs), and cognitive decline. Consequently, targeting mTOR with inhibitors may hold promise for the prevention and treatment of AD.

In this study, we conducted high-throughput screening of FDA compounds with and without rapamycin to identify their synergistic effect on yeast growth. The compounds identified as positive in the screening were then tested in the *C. elegans* AD model to assess their impact on survival. It was found that fluphenazine and trimethiazide significantly delayed paralysis in combination with rapamycin. Additionally, at least six of the tested compounds were observed to delay paralysis, with fluphenazine being identified as robustly delaying it. Furthermore, we measured various health metrics including fecundity assay, body bends, and body length. The results showed improvement in these health parameters when treated with fluphenazine compared to the control. Lastly, it was observed that there was a significant reduction in amyloid Beta accumulation in worms treated with fluphenazine. Overall, this data shows that repurposed drugs, particularly fluphenazine, holds promise as a potential candidate for further investigation in the context of lifespan extension and ameliorating Alzheimer’s disease.

## Results

### Rapamycin shows synergistic effect with FDA compounds on *S. cerevisiae* growth

To investigate the effect of various compounds on cell growth under mTOR complex 1 (mTORC1) inhibition, we conducted a high-throughput screen of a library containing 1,760 FDA-approved compounds in *S. cerevisiae*, both in the presence and absence of rapamycin, a specific inhibitor of TORC1. Initially, we determined a concentration of rapamycin that did not affect yeast growth on its own (Fig. S1). We then used this concentration in combination with a 10 µM final concentration of each FDA compound and analyzed the effect on yeast growth over 24 hours, taking optical density (OD) readings every 30 minutes (Fig. 1A, Table S1). Compounds that did not show growth defects on their own but caused significant growth defects when combined with the sub-inhibitory concentration of rapamycin were identified as positive candidates (Fig. 1B). After two rounds of screening the entire library, 87 positive hit compounds were selected (Fig. 1C, Table S2).

**Figure 1.**
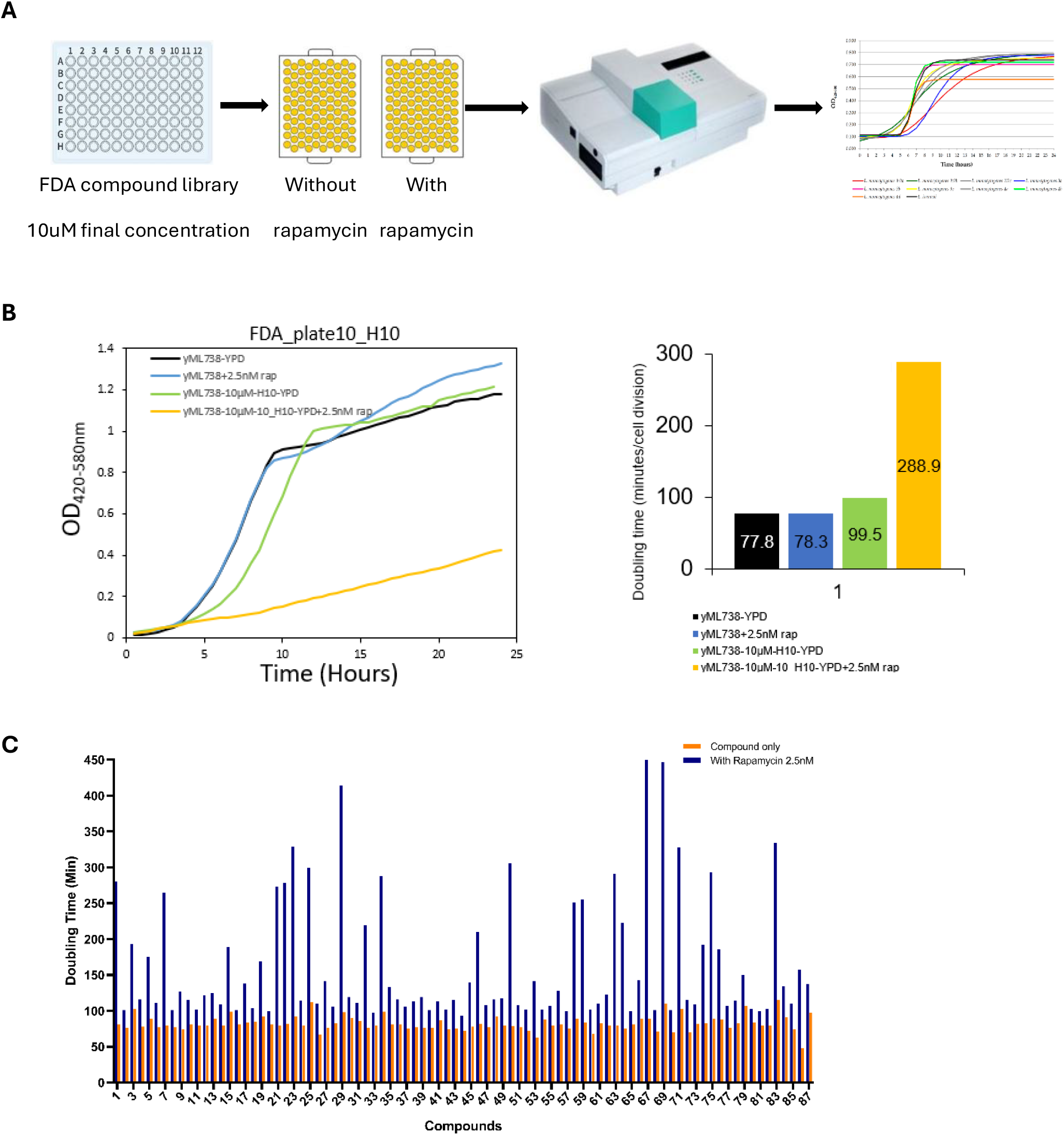
Rapamycin shows synergistic effects on *S. cerevisiae* growth in combination with FDA-approved compounds. (A) Experiment design. The BY4741 strain was seeded in YPD media with a single FDA compound or with rapamycin or both and OD at 420nM was analyzed every 30min for 24 hours. (B) Representative image of positive hit. If only FDA compound plus rapamycin combination showed a growth defect it was considered as positive hit. Graph shows the average doubling time in minutes for yeast in different conditions. (C) Graph showing number of identified positive hits in yeast screen.

Previous studies have shown that *fpr1Δ* strains are resistant to rapamycin [25, 26]. FPR1 encodes a protein that binds rapamycin, and the complex of rapamycin with FPR1 allosterically inhibits TORC1. To determine if the observed synergistic effect was indeed due to TORC1 inhibition, we tested the positive hit compounds in the *fpr1Δ* strain with and without rapamycin. Interestingly, none of the positive hit compounds affected cell growth in the presence of rapamycin in this strain (Fig. S2), confirming that the observed effects are indeed due to TORC1 inhibition.

### Rapamycin and FDA compounds synergistically delayed paralysis in *C. elegans* Alzheimer’s disease model

To understand the effect of these positive hit compounds on amyloid pathology, we utilized *C. elegans* Alzheimer’s disease model, GMC101 strain. The rapid paralysis induced by Aß_1-42_ expression upon temperature shift in this *C. elegans* strain makes it an ideal model for rapidly evaluating drug effects [8]. Among the tested compounds, at least three demonstrated a synergistic effect on the onset of paralysis in this background when combined with rapamycin. Initially, L4 stage worms were exposed to 80 µM trichlormethiazide, a diuretic compound, and 50 µM rapamycin and incubated at 25^0^C. When treated individually or in combination at these concentrations, there was no significant effect on the onset of paralysis in the GMC101 strain (Fig. 2A). However, increasing the trichlormethiazide concentration to 160 µM while maintaining 50 µM rapamycin resulted in delayed paralysis by 25% (Fig. 2B). Fluphenazine, compound in the phenothiazine class, exhibited a robust delay in paralysis when used alone. At 25 µM, fluphenazine delayed paralysis by 178%, whereas at 50 µM, by 915% compared to the control (Fig. 3A). When combined with 10 µM rapamycin, the 25 µM fluphenazine dose resulted in an even greater delay in paralysis by 387.5%. However, combining 50 µM fluphenazine with rapamycin did not result in further delay (Fig. 2C, D). Perphenazine, another phenothiazine class compound, also showed significant delay in paralysis. At 80 µM, perphenazine delayed paralysis by 460%, and at 160 µM, by 223% (Fig. 3B). Interestingly, when combined with 10 µM rapamycin, unparalyzed worm percentage was reduced at 80 µM to 322% and even further at 160 µM to 73.9% (Fig. 2E, F). These findings indicate that while some compounds like trichloromethiazide and fluphenazine can synergize with rapamycin to delay paralysis, others like perphenazine may have a complex interaction where higher concentrations in combination with rapamycin can be detrimental.

**Figure 2.**
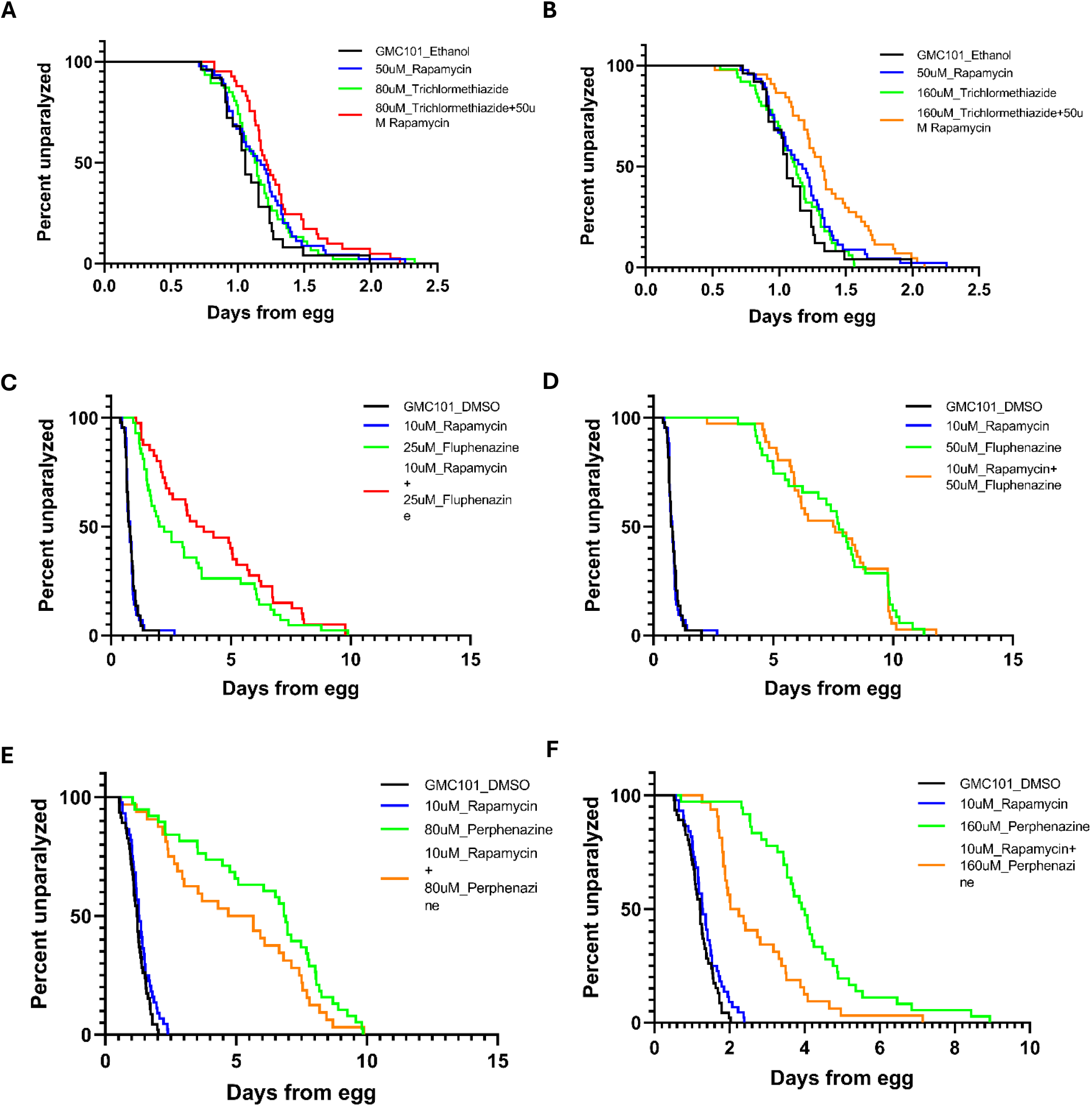
Rapamycin and FDA approved compounds synergistically delayed paralysis in C. elegans Alzheimer’s disease model. (A), (B) Trichlormethiazide delayed paralysis synergistically with 50uM rapamycin at 160uM conc. (C), (D) 25uM Fluphenazine delayed paralysis synergistically with 10uM rapamycin. No additional delay in paralysis was observed at 50uM conc. (E), (F) 80uM concentration. Perphenazine delayed paralysis in combination with 10uM rapamycin and further increased when concentration was increased to 160uM. Quantitative data and statistical analyses for the representative experiments are included in Table 1.

**Figure 3.**
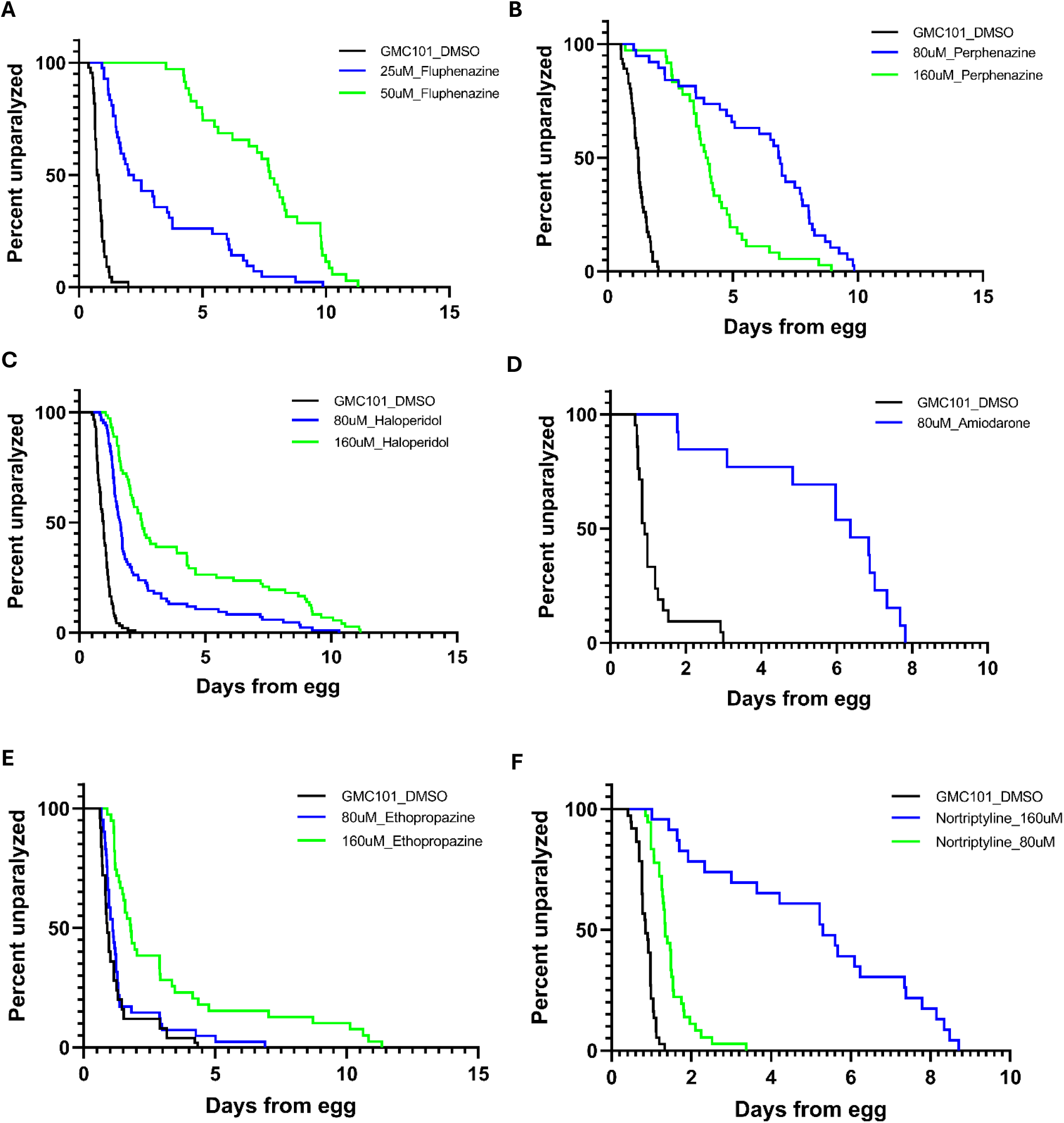
Repurposing FDA approved drugs delayed paralysis in C. elegans Alzheimer’s disease model. (A) to (F) FDA approved compounds were used to treat GMC 101 with indicated concentrations and paralysis was analyzed. These compounds showed robust suppression of paralysis. Quantitative data and statistical analyses for the representative experiments are included in Table 1.

### Repurposing FDA-approved drugs delayed paralysis in *C. elegans* Alzheimer’s disease model

In our screen we observed fluphenazine and perphenazine delayed paralysis on their own at higher concentrations. So, we decided to analyze the effect of a few other compounds from the positive hit list on worm paralysis. We observed at least four more compounds that delayed the onset of paralysis. Treatment with 80 μM and 160 μM Haloperidol, an antipsychotic agent, led to an average delay in the onset of paralysis by 125% and 563%, respectively, compared to the control group (Table 1, Fig 3C). Similarly, Amiodarone, an anti-arrhythmic drug, delayed paralysis by 598% at 80uM. However, at 160uM worms foraged off plate, likely as a response to the drug, so the effect could not be analyzed (Fig 3D). Ethopropazine, another compound in the phenothiazine class, did not have effect at 80uM. However, at 160uM concentration delayed the onset of paralysis by almost double as compared to control (Fig 3E). Nortriptyline, a tricyclic antidepressant, 80uM and 160uM treatment led to delay in paralysis by 59.1% and 525.7% respectively (Fig. 3F).

**Table 1.** Median survival of worms using different concentrations for FDA approved drugs with or without rapamycin and percent change increase in survival.

| Sr.No. | Compound | Median survival<br>(in Days) | % change | P value (Log-rank (Mantel-Cox) test) |
| --- | --- | --- | --- | --- |
| 1 | GMC101_DMSO | 0.92 |  |  |
| 2 | 10uM_Rapamycin | 1.08 | 16.6 | 0.3051 (GMC101_DMSO vs 10uM_Rapamycin) |
| 3 | 80uM_Haloperidol | 2.08 | 125.4 | 0.0001 (GMC101_DMSO vs 80uM_Haloperidol) |
| 4 | 10uM_Rapamycin + 80uM_Haloperidol | 1.88 | 104.3 | 0.6313 (80uM_Haloperidol vs 10uM_Rapamycin + 80uM_Haloperidol) |
| 5 | 160uM_Haloperidol | 6.12 | 563.5 | 0.0001 (GMC101_DMSO vs 160uM_Haloperidol) |
| 6 | 10uM_Rapamycin + 160uM_Haloperidol | 6.65 | 620.9 | 0.3922 (160uM_Haloperidol vs 10uM_Rapamycin + 160uM_Haloperidol) |
| 7 | GMC101_DMSO | 1.06 |  |  |
| 8 | 50uM_Rapamycin | 1.17 | 10.7 | 0.1354 (GMC101_DMSO vs 50uM_Rapamycin) |
| 9 | 80uM_Trichlormethiazide | 1.14 | 7.3 | 0.2177 (GMC101_DMSO vs 80uM_Trichlormethiazide) |
| 10 | 80uM_Trichlormethiazide+50uM_Rapamycin | 1.22 | 3.6 | 0.0854 (80uM_Trichlormethiazide vs 50uM_Rapamycin + 80uM_Trichlormethiazide) |
| 11 | 160uM_Trichlormethiazide | 1.12 | 6.1 | 0.4958 (GMC101_DMSO vs 160uM_Trichlormethiazide) |
| 12 | 50uM_Rapamycin + 160uM_Trichlormethiazide | 1.33 | 25.3 | 0.0001 (160uM_Trichlormethiazide vs 50uM_Rapamycin + 160uM_Trichlormethiazide) |
| 13 | GMC101_DMSO | 0.76 |  |  |
| 14 | 10uM_Rapamycin | 0.79 | 3.5 | 0.9692 (GMC101_DMSO vs 10uM_Rapamycin) |
| 15 | 25uM_Fluphenazine | 2.12 | 177.8 | 0.0001 (GMC101_DMSO vs 25uM_Fluphenazine) |
| 16 | 10uM_Rapamycin+ 25uM_Fluphenazine | 3.72 | 387.4 | 0.028 (25uM_Fluphenazine vs 10uM_Rapamycin+ 25uM_Fluphenazine) |
| 17 | 50uM_Fluphenazine | 7.74 | 915.2 | 0.0001 (GMC101_DMSO vs 50uM_Fluphenazine) |
| 18 | 10uM_Rapamycin+ 50uM_Fluphenazine | 7.54 | 888.9 | 0.8905 (50uM_Fluphenazine vs 10uM_Rapamycin+ 50uM_Fluphenazine) |
| 19 | GMC101_DMSO | 1.14 |  |  |
| 20 | 10uM_Rapamycin | 1.03 | -10.1 | 0.358 (GMC101_DMSO vs 10uM_Rapamycin) |
| 21 | 80uM_Fluocinolone | 1.15 | 0.6 | 0.2828 (GMC101_DMSO vs 80uM_Fluocinolone) |
| 22 | 10uM_Rapamycin+ 80uM_Fluocinolone | 0.95 | -16.3 | 0.1177 (80uM_Fluocinolone vs 10uM_Rapamycin + 80uM_Fluocinolone) |
| 23 | 160uM_Fluocinolone | 1.21 | 6.3 | 0.2011 (GMC101_DMSO vs 160uM_Fluocinolone) |
| 24 | 10uM_Rapamycin+ 160uM_Fluocinolone | 1.03 | -9.7 | 0.4136 (160uM_Fluocinolone vs 50uM_Rapamycin + 160uM_Fluocinolone) |
| 25 | GMC101_DMSO | 1.23 |  |  |
| 26 | 10uM_Rapamycin | 1.29 | 4.7 | 0.0985 (GMC101_DMSO vs 10uM_Rapamycin) |
| 27 | 80uM_Perphenazine | 6.86 | 459.1 | 0.0001 (GMC101_DMSO vs 80uM_Perphenazine) |
| 28 | 10uM_Rapamycin+ 80uM_Perphenazine | 5.18 | 322.0 | 0.0451 (80uM_Perphenazine vs 10uM_Rapamycin + 80uM_Perphenazine) |
| 29 | 160uM_Perphenazine | 3.97 | 223.0 | 0.0001 (GMC101_DMSO vs 160uM_Perphenazine) |
| 30 | 10uM_Rapamycin+ 160uM_Perphenazine | 2.13 | 73.9 | 0.0001 (160uM_Perphenazine vs 10uM_Rapamycin + 160uM_Perphenazine) |
| 31 | GMC101_DMSO | 0.91 |  |  |
| 32 | 10uM_Rapamycin | 0.94 | 3.1 | 0.3276 (GMC101_DMSO vs 10uM_Rapamycin) |
| 33 | 80uM_Amiodarone | 6.36 | 598.4 | 0.0001 (GMC101_DMSO vs 80uM_Amiodarone) |
| 34 | 10uM_Rapamycin+ 80uM_Amiodarone | 5.68 | 523.5 | 0.5612 (80uM_Amiodarone vs 10uM_Rapamycin + 80uM_Amiodarone) |
| 35 | GMC101_DMSO | 0.90 |  |  |
| 36 | 80uM_Ethopropazine | 1.11 | 23.4 | 0.1464 (GMC101_DMSO vs 80uM_Ethopropazine) |
| 37 | 160uM_Ethopropazine | 1.79 | 99.3 | 0.0001 (GMC101_DMSO vs 160uM_Ethopropazine) |
| 38 | GMC101_DMSO | 0.85 |  |  |
| 39 | 80uM_Nortriptyline | 1.35 | 59.2 | 0.0001 (GMC101_DMSO vs 80uM_Nortriptyline) |
| 40 | 160uM_Nortriptyline | 5.30 | 525.7 | 0.0001 (GMC101_DMSO vs 160uM_Nortriptyline) |
| 41 | GMC101_DMSO | 0.84 |  |  |
| 42 | 40uM_Chlorpromazine | 1.83 | 116.9 | 0.0001 (GMC101_DMSO vs 40uM_Chlorpromazine) |
| 43 | 80uM_Chlorpromazine | 10.94 | 1199.4 | 0.0001 (GMC101_DMSO vs 80uM_Chlorpromazine) |

### Fluphenazine improved additional health metrics and reduced amyloid beta accumulation

The phenothiazine class of compounds showed the largest delay in the onset of paralysis in GMC101 animals. To determine whether these compounds improved or at least maintained normal health span, evaluated the effect of fluphenazine treatment on total fecundity of GMC101 animals by monitoring brood size after every 24hr. The untreated animals at the first day of adulthood had about 73 progenies (3 progenies/worm/hr); however, the fluphenazine treated worms on an average had only 29 (1.21 progenies/worm/hr). The untreated worms did not survive after day 1; however, treated worms not only survived but were also reproductively active until day 4 (Fig. 4A). Basal slowing response (BSR) behavior depends on the functionality of the dopaminergic neurons, and it is measured by the change in body bends [27]. Therefore, we assessed the effects of fluphenazine on the motility of *C. elegans* by analyzing the body bend rate. The number of body bends every minute were counted on day 1 of adulthood. The fluphenazine-treated groups exhibited a higher swinging motion compared to the untreated group (Fig. 4B). Additionally, we tested body length and width in fluphenazine treated and untreated L4 and Day 1 worms. Fluphenazine treated worms were significantly shorter and narrower than control worms on day 1 (Fig. 4C, D). This data suggests that fluphenazine treatment slowed the growth of the worms, however worms were in better health overall as compared to control.

**Figure 4.**
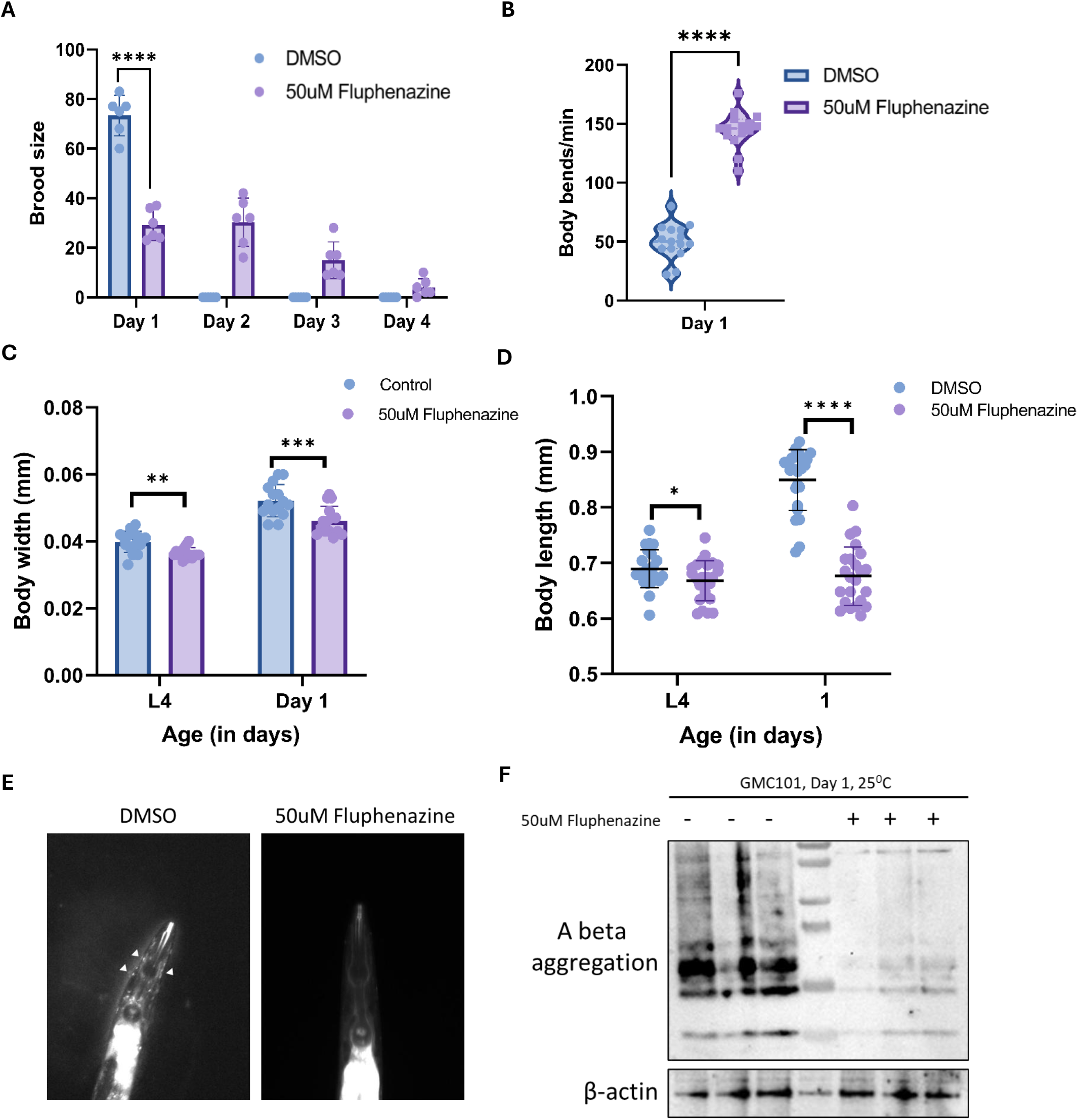
Fluphenazine improved health metrics and reduced amyloid beta accumulation. (A) Total number of progenies hatched each day counted from 50uM fluphenazine treated or untreated worms. (B) Comparison of thrashing between control and 50uM treated animals. Untreated worms showed significantly less trashing compared to the treated worms. (C), (D) Influence of 50uM Fluphenazine treatment on worm size. Body width and body length was measured in L4 and day 1 adult worms treated or not with 50uM Fluphenazine. (E) X-34 positive Aß1-42 aggregates are seen (arrowhead), untreated control (left side) and 50uM fluphenazine treated (right side). (F) Total protein extracts from untreated and 50uM fluphenazine treated GMC 101 were resolved via tricine-SDS-PAGE with immuno detection of Aß and ß-actin. p-Values were obtained by using unpaired two-tailed t-tests with Welch’s correction; **** = p-value < 0.0001, ** = p-value < 0.01 * = p-value < 0.05.

The primary pathological hallmark of Alzheimer’s disease is the accumulation of plaques in the brain, primarily composed of amyloid-beta (Aß) peptides [28]. This amyloid-beta accumulation in worms leads to rapid paralysis and subsequent death. We observed that worms treated with fluphenazine exhibited significantly higher motility compared to untreated worms, leading us to hypothesize that fluphenazine treatment may reduce amyloid-beta accumulation. To test the hypothesis, aggregated Aß1-42 was detected *in vivo*, via staining in live animals using the lipophilic congo red derivative, X-34 [29]. Untreated worms showed deposits of amyloid-beta aggregates in the head region, while fluphenazine treatment significantly reduced this accumulation (Fig. 4E). To further confirm these results, the amount of Aß1-42 protein expressed in fluphenazine treated and untreated worms was compared via immune-blot analysis. The results showed significant reduction in Aß1-42 protein expression in fluphenazine treated worms (Fig. 4F). These results suggest that fluphenazine may delay paralysis and improve health in GMC101 worms by reducing Aß1-42 accumulation.

## Discussion

Our study demonstrates the potential of drug repurposing for the treatment of Alzheimer’s disease (AD) by leveraging the synergistic effects of existing FDA-approved compounds with rapamycin. Using *Saccharomyces cerevisiae* as an initial screening model, we identified 87 compounds that exhibited growth defects when combined with rapamycin, suggesting possible interactions with the mTOR signaling pathway. Further validation in a *Caenorhabditis elegans* model of AD revealed seven compounds capable of delaying paralysis, with three demonstrating significant synergy with rapamycin. The identification of fluphenazine as a promising candidate further underscores the utility of repurposed drugs in mitigating AD pathology. Our Western blot and X-34 staining results confirmed that fluphenazine significantly reduced amyloid-beta accumulation, a hallmark of AD. This supports the notion that fluphenazine may influence protein aggregation and clearance mechanisms, which are critical for neurodegenerative disease intervention.

The ability of rapamycin to extend lifespan and delay age-related diseases, including neurodegeneration, has been well-documented [30]. Our findings suggest that certain FDA-approved compounds can enhance rapamycin’s beneficial effects, potentially through mechanisms related to proteostasis, mitochondrial function, or stress response pathways. Previous studies have used rapamycin at concentrations exceeding 100 µM to demonstrate lifespan extension in *C. elegans* [31-33]. In contrast, we employed significantly lower concentrations (10 µM and 50 µM) to minimize adverse effects associated with strong mTOR inhibition. While these concentrations synergistically interacted with FDA-approved compounds such as trichlormethiazide and fluphenazine to delay paralysis (Fig. 2), the addition of rapamycin in combination with higher concentrations of perphenazine resulted in accelerated paralysis, suggesting a potential toxicity or off-target effects.

Phenothiazines, the class of drugs are primarily used as antipsychotics due to their antagonistic effects on dopamine, histamine H1, and serotonergic 5-hydroxytryptamine (HT) 2 receptors [34]. These compounds share a tricyclic structure and are known to modulate multiple neurotransmitter systems, including dopaminergic, serotonergic, and adrenergic pathways. Phenothiazines have also been shown to interact with ion channels [35], oxidative stress pathways [36], and autophagy-related mechanisms [37], making them of interest in neurodegenerative disease research. Recently Dao et al., developed a Phenothiazine-based theranostic compounds which effectively prevents Aβ fibril formation and disaggregate preformed Aβ fibrils [38]. This suggests that phenothiazine may hold therapeutic potential for Alzheimer’s disease by not only inhibiting amyloid aggregation but also reversing existing fibril deposits, potentially mitigating disease progression.

Fluphenazine, a typical antipsychotic primarily used to treat schizophrenia, functions as a dopamine D2 receptor antagonist. Dysregulation of dopamine receptors has been implicated in AD, particularly in relation to synaptic plasticity and neuronal survival [39]. Additionally, fluphenazine has been reported to interact with sigma-1 receptors [40], which are involved in modulating neuroprotection, calcium homeostasis, and amyloid-beta metabolism. Sigma-1 receptor activation has been shown to reduce oxidative stress and prevent neurodegeneration [41], suggesting a possible mechanism by which fluphenazine may exert its protective effects. Furthermore, fluphenazine has been found to affect autophagic pathways [36], which are crucial for the degradation of misfolded proteins, including amyloid-beta. By enhancing autophagy, fluphenazine may contribute to the clearance of toxic aggregates, thereby mitigating AD pathology.

While our study provides compelling evidence for the efficacy of certain repurposed drugs, several limitations must be considered. First, the use of yeast and *C. elegans* as model organisms, while valuable for high-throughput screening and mechanistic insights, does not fully recapitulate the complexities of human AD pathology. Further validation in mammalian models is essential to confirm the neuroprotective effects of the identified compounds. Second, the precise molecular mechanisms underlying the observed lifespan extension and amyloid-beta reduction remain to be elucidated. Future studies employing transcriptomic and proteomic analyses could provide deeper insights into the pathways involved.

Additionally, the pharmacokinetics and safety profiles of these compounds in humans need to be reassessed in the context of AD treatment. Although these drugs are already FDA-approved for other indications, their dosage, bioavailability, and long-term effects in neurodegenerative conditions warrant further investigation.

In conclusion, our findings highlight the promise of drug repurposing in accelerating the discovery of novel AD therapeutics. The identification of compounds that synergize with rapamycin provides a strong foundation for future studies aimed at optimizing combination therapies. Moving forward, translating these findings into clinical settings will require rigorous preclinical and clinical evaluations to assess efficacy and safety in AD patients.

## Methods and Material

### Strains

In this study, we used *BY4741* (*MATa his3Δ1 leu2Δ0 met15Δ0 ura3Δ0*) wild type *S. cerevisiae* and *fpr1Δ* strain. Cells were grown in standard YPD liquid media containing Bacto peptone 20 g/L (Becton Dickinson), yeast extract 10 g/L (Becton Dickinson), and dextrose 20 g/L (Junsei) and incubated at 30°C. Standard C. elegans procedures were used as previously described [42, 43]. For experiments in *C. elegans*, we used the N2 wild type as well as GMC101 (dvIs100[unc-54p::human Aβ1-42::unc-54 3′-UTR + mtl-2p:GFP]) model for Alzheimer’s disease. Strains were cultured and maintained using standard methods [44]. All worm lines were maintained on NGM plates seeded with live OP50 bacteria and maintained at 20 °C.

### Growth rate analysis

Growth rate analysis was performed as described in [45]. Briefly, to investigate the growth defect in yeast, BY4741 or *fpr1Δ* strain was grown in 150uL YPD media in honeycomb well plates (Fischer Scientific) containing only FDA approved compound (10uM final conc.) or in combination with rapamycin (2.5ng/ml final conc.). Optical density at 420nM and 580nm was measured every 30min for 24hrs using Bioscreen C MBR instrument. Raw optical density data were smoothed using the R package “smooth.spline.” Doubling times were calculated using the inflection method—identifying the maximum semi-log slope along growth curves within the optical density range linearly correlated with a number of yeast cells—with the online web tool Yeast Outgrowth Data Analyzer (YODA)[46, 47]

### Worm paralysis assay

Paralysis was assessed by standard methods [8, 48]. Briefly, the worms were bleach synchronized and cultivated on NGM plates with OP50 bacterial lawns. These worms were maintained at 20 °C until they reached late-stage larval L4 stage and then moved to a plate with FUDR (12 well plate) with either DMSO (control), rapamycin only, compound only and rapamycin plus compound. Then these plates were shifted to 25°C and body movement assessed over time using wormbot [49]. Statistical significance between each experimental condition calculated via Log-rank (Mantel-Cox) test.

### Lifespan analysis

Manual lifespan analysis was carried out according to standard methods [43]. Briefly, animals were bleach synchronized and then cultivated on NGM plate at 20^0^C. Animals were then visually assessed to be late L4 stage and manually transferred to plates containing 1 μM FuDR and increasing concentrations of FDA compounds. Worms were then maintained on these plates transferring worms to fresh plates every few days and manually examined each day for death events. Only confirmed death events were quantified; therefore, they were not influenced; the metrics are not influenced by worms lost off the plates. All death events were counted with no censoring. Each biological replicate contained at least three plates each with 30 + worms each. Statistical significance between each experimental condition calculated via Log-rank (Mantel-Cox) test.

### Worm growth assay

To compare growth rates of control and 50uM Fluphenazine treated GMC101worms, we bleached synchronized lines and then propagated them from egg for 48 h at 20 °C and then L4 stage worms were transferred on plates treated with or not 50uM Fluphenazine and shifted to 25°C for 4hrs and 24hrs. Then worms were imaged for each condition. Then quantified it using ImageJ to quantify the length by trancing the midline of each animal [50]. The pixel lengths were converted to microns in comparison to a ruler imaged at the same magnification. Statistical significance was calculated using unpaired, parametric t-tests with Welch’s correction using GraphPad Prism software.

### Fecundity assay

For brood size measurement, we bleached synchronized lines and then propagated them from egg for 48 h at 20 °C and then single L4 stage worm was transferred on plates treated or not with 50uM Fluphenazine and shifted to 25°C. After 24hrs worms were transferred to a new plate and progenies were counted from each plate after 2 days. Statistical significance was calculated using unpaired, parametric t-tests with Welch’s correction using GraphPad Prism software.

### Swimming body bend assays

Worm cohorts were bleach synchronized and then maintained on NGM plates seeded with OP50 bacteria. Subsequently, larval stage 4 animals were visually assessed and transferred to a fresh plate with or without 50uM Fluphenazine and shifted to 25°C. Some of these transferred animals were then analyzed for body bend rate after 4hrs, and then after 24hrs. The thrashing rate was determined by standard methods [51]. For each recording, a single animal was transferred to a 10 μL drop of M9 buffer placed on a siliconized microscope slide. Each worm was allowed to acclimate for 30 s, and then, each thrashing cycle was manually recorded using a cell counter. A single thrash was defined strictly as a C-shaped, head-to-tail movement.

### X-34 staining

L4 stage worm was transferred on plates treated or not with 50uM Fluphenazine and shifted to 25°C, after 16hrs, worms were transferred to 18 microliters drop of the 1mM X-34 solution for a 2 hr incubated at room temp. Picked the worms in to a drop of S-basal to wash them, then transferred them on a spotted NGM plate and let them destain at 16 degrees overnight, then mounted them on an agar pad for imaging. Imaging was done using DAPI filter sets with 360 nm wavelength.

### Western Blot

Western blotting was performed as described in [52]. Briefly, worm cohorts were bleach synchronized and then maintained on NGM plates seeded with OP50 bacteria. Subsequently, larval stage 4 animals were visually assessed and transferred to a fresh plate with or without 50uM Fluphenazine and shifted to 25°C for 24hrs. Worms were collected and washed 3x with M9 and the pellet was flash frozen. 100 μL lysis buffer (100mM NaCl, 100mM Tris pH 7.5, 1% NP-40) supplemented with an EDTA-free Protease Inhibitor cocktail tablet (Roche) added to the pellet along with glass beads and vortexed and centrifuged 15 minutes at 4°C. The supernatant was transferred to a new tube and 20 μL was set aside for protein quantification (Pierce BCA protein assay kit). 2x protein sample buffer (80 mM Tris-HCl, 2% SDS, 10% glycerol, 0.0006% Bromophenol blue with 10% β-mercaptoethanol and 8M Urea) was added to the supernatant. Samples were heated at 55°C for 5 minutes and 30 μg of protein per sample was run on Tris-Tricine 16.5% precast polyacrylamide gels (BioRad) for Aβ or 10% Tris-Glycine (TGX) precast protein gels (BioRad) for β-actin. Samples were transferred onto a nitrocellulose membrane (BioRad) for 40 minutes at 70V. Membranes were blocked in 5% non-fat milk in TBS with 0.1% Tween-20 one hour at room temperature. Primary antibody incubation was overnight at 4°C; Aβ (6E10 Biolegend cat#803001, 1:1000) and β-actin (Sigma cat# A2228, 1:5000) antibodies were used. Incubation with a goat anti-Mouse IgG-HRP (ThermoFisher Scientific cat#31430, 1:5000) secondary antibody conjugated to horseradish peroxidase was 1 ½ hr. at room temperature. Chemiluminescence detection used SuperSignal West Pico PLUS (ThermoFisher Scientific) and an iBright Imaging System (ThermoFisher Scientific).

## Notes

### Competing Interest Statement

The authors have declared no competing interest.

### Summary of Updates

The author's affiliation has been modified.

## References

1. Maniam, S. and S. Maniam, Screening Techniques for Drug Discovery in Alzheimer’s Disease. ACS Omega, 2024. 9(6): p. 6059–6073.

2. Vaiserman, A., et al., Repurposing drugs to fight aging: The difficult path from bench to bedside. Med Res Rev, 2021. 41(3): p. 1676–1700.

3. Antoszczak, M., et al., Old wine in new bottles: Drug repurposing in oncology. Eur J Pharmacol, 2020. 866: p. 172784.

4. Kumar, A., A. Singh, and Ekavali, A review on Alzheimer’s disease pathophysiology and its management: an update. Pharmacol Rep, 2015. 67(2): p. 195–203.

5. De-Paula, V.J., et al., Alzheimer’s disease. Subcell Biochem, 2012. 65: p. 329–52.

6. Barberis, A., et al., Yeast as a screening tool. Drug Discov Today Technol, 2005. 2(2): p. 187–92.

7. Chen, X., et al., Using C. elegans to discover therapeutic compounds for ageing-associated neurodegenerative diseases. Chem Cent J, 2015. 9: p. 65.

8. McColl, G., et al., Utility of an improved model of amyloid-beta (Abeta(1)(−)(4)(2)) toxicity in Caenorhabditis elegans for drug screening for Alzheimer’s disease. Mol Neurodegener, 2012. 7: p. 57.

9. Fang, E.F., et al., Mitophagy inhibits amyloid-beta and tau pathology and reverses cognitive deficits in models of Alzheimer’s disease. Nat Neurosci, 2019. 22(3): p. 401–412.

10. Hassan, W.M., et al., Identifying Abeta-specific pathogenic mechanisms using a nematode model of Alzheimer’s disease. Neurobiol Aging, 2015. 36(2): p. 857–66.

11. Lublin, A.L. and C.D. Link, Alzheimer’s disease drug discovery: in vivo screening using Caenorhabditis elegans as a model for beta-amyloid peptide-induced toxicity. Drug Discov Today Technol, 2013. 10(1): p. e115–e119.

12. Hardaway, J.A., et al., Forward genetic analysis to identify determinants of dopamine signaling in Caenorhabditis elegans using swimming-induced paralysis. G3 (Bethesda), 2012. 2(8): p. 961–75.

13. Barclay, J.W., A. Morgan, and R.D. Burgoyne, Neurotransmitter release mechanisms studied in Caenorhabditis elegans. Cell Calcium, 2012. 52(3-4): p. 289–95.

14. Aaldijk, E. and Y. Vermeiren, The role of serotonin within the microbiota-gut-brain axis in the development of Alzheimer’s disease: A narrative review. Ageing Res Rev, 2022. 75: p. 101556.

15. Laplante, M. and D.M. Sabatini, mTOR signaling at a glance. Journal of Cell Science, 2009. 122(20): p. 3589–3594.

16. Cai, Z., et al., Activation of mTOR: a culprit of Alzheimer’s disease? Neuropsychiatr Dis Treat, 2015. 11: p. 1015–30.

17. Papadopoli, D., et al., mTOR as a central regulator of lifespan and aging. F1000Res, 2019. 8.

18. Weichhart, T., mTOR as Regulator of Lifespan, Aging, and Cellular Senescence: A Mini-Review. Gerontology, 2018. 64(2): p. 127–134.

19. Huang, S., mTOR Signaling in Metabolism and Cancer. Cells, 2020. 9(10).

20. Fu, W. and G. Wu, Targeting mTOR for Anti-Aging and Anti-Cancer Therapy. Molecules, 2023. 28(7).

21. Yang, L., et al., Targeting mTOR Signaling in Type 2 Diabetes Mellitus and Diabetes Complications. Curr Drug Targets, 2022. 23(7): p. 692–710.

22. Cao, Y., et al., Targeting mTOR Signaling by Dietary Polyphenols in Obesity Prevention. Nutrients, 2022. 14(23).

23. Chong, Z.Z., Y.C. Shang, and K. Maiese, Cardiovascular disease and mTOR signaling. Trends Cardiovasc Med, 2011. 21(5): p. 151–5.

24. Querfurth, H. and H.K. Lee, Mammalian/mechanistic target of rapamycin (mTOR) complexes in neurodegeneration. Mol Neurodegener, 2021. 16(1): p. 44.

25. Heitman, J., et al., FK 506-binding protein proline rotamase is a target for the immunosuppressive agent FK 506 in Saccharomyces cerevisiae. Proc Natl Acad Sci U S A, 1991. 88(5): p. 1948–52.

26. Koltin, Y., et al., Rapamycin sensitivity in Saccharomyces cerevisiae is mediated by a peptidyl-prolyl cis-trans isomerase related to human FK506-binding protein. Mol Cell Biol, 1991. 11(3): p. 1718–23.

27. Rivard, L., et al., A comparison of experience-dependent locomotory behaviors and biogenic amine neurons in nematode relatives of Caenorhabditis elegans. BMC Neurosci, 2010. 11: p. 22.

28. Masters, C.L., et al., Amyloid plaque core protein in Alzheimer disease and Down syndrome. Proc Natl Acad Sci U S A, 1985. 82(12): p. 4245–9.

29. Styren, S.D., et al., X-34, a fluorescent derivative of Congo red: a novel histochemical stain for Alzheimer’s disease pathology. J Histochem Cytochem, 2000. 48(9): p. 1223–32.

30. Selvarani, R., S. Mohammed, and A. Richardson, Effect of rapamycin on aging and age-related diseases-past and future. Geroscience, 2021. 43(3): p. 1135–1158.

31. Seo, K., et al., Heat shock factor 1 mediates the longevity conferred by inhibition of TOR and insulin/IGF-1 signaling pathways in C. elegans. Aging Cell, 2013. 12(6): p. 1073–81.

32. Xie, J., et al., Regulation of the Elongation Phase of Protein Synthesis Enhances Translation Accuracy and Modulates Lifespan. Curr Biol, 2019. 29(5): p. 737–749 e5.

33. Peng, H.H., et al., Ganoderma lucidum stimulates autophagy-dependent longevity pathways in Caenorhabditis elegans and human cells. Aging (Albany NY), 2021. 13(10): p. 13474–13495.

34. Sudeshna, G. and K. Parimal, Multiple non-psychiatric effects of phenothiazines: a review. Eur J Pharmacol, 2010. 648(1-3): p. 6–14.

35. Mehrabi, S.F., S. Elmi, and J. Nylandsted, Repurposing phenothiazines for cancer therapy: compromising membrane integrity in cancer cells. Front Oncol, 2023. 13: p. 1320621.

36. Duarte, D. and N. Vale, Antipsychotic Drug Fluphenazine against Human Cancer Cells. Biomolecules, 2022. 12(10).

37. Lopes, R.M., et al., Targeting autophagy by antipsychotic phenothiazines: potential drug repurposing for cancer therapy. Biochem Pharmacol, 2024. 222: p. 116075.

38. Dao, P., et al., Development of Phenothiazine-Based Theranostic Compounds That Act Both as Inhibitors of beta-Amyloid Aggregation and as Imaging Probes for Amyloid Plaques in Alzheimer’s Disease. ACS Chem Neurosci, 2017. 8(4): p. 798–806.

39. Pan, X., et al., Dopamine and Dopamine Receptors in Alzheimer’s Disease: A Systematic Review and Network Meta-Analysis. Front Aging Neurosci, 2019. 11: p. 175.

40. Milenina, L.S., et al., Sigma-1 Receptor Ligands Chlorpromazine and Trifluoperazine Attenuate Ca(2+) Responses in Rat Peritoneal Macrophages. Cell tissue biol, 2022. 16(3): p. 233–244.

41. Nguyen, L., et al., Role of sigma-1 receptors in neurodegenerative diseases. J Pharmacol Sci, 2015. 127(1): p. 17–29.

42. Kaeberlein, T.L., et al., Lifespan extension in Caenorhabditis elegans by complete removal of food. Aging Cell, 2006. 5(6): p. 487–94.

43. Sutphin, G.L. and M. Kaeberlein, Measuring Caenorhabditis elegans life span on solid media. J Vis Exp, 2009(27).

44. Brenner, S., The genetics of Caenorhabditis elegans. Genetics, 1974. 77(1): p. 71–94.

45. Murakami, C. and M. Kaeberlein, Quantifying yeast chronological life span by outgrowth of aged cells. J Vis Exp, 2009(27).

46. Olsen, B., C.J. Murakami, and M. Kaeberlein, YODA: software to facilitate high-throughput analysis of chronological life span, growth rate, and survival in budding yeast. BMC Bioinformatics, 2010. 11: p. 141.

47. Lee, M.B., et al., Pterocarpus marsupium extract extends replicative lifespan in budding yeast. Geroscience, 2021. 43(5): p. 2595–2609.

48. Steinkraus, K.A., et al., Dietary restriction suppresses proteotoxicity and enhances longevity by an hsf-1-dependent mechanism in Caenorhabditis elegans. Aging Cell, 2008. 7(3): p. 394–404.

49. Pitt, J.N., et al., WormBot, an open-source robotics platform for survival and behavior analysis in C. elegans. Geroscience, 2019. 41(6): p. 961–973.

50. Rueden, C.T., et al., ImageJ2: ImageJ for the next generation of scientific image data. BMC Bioinformatics, 2017. 18(1): p. 529.

51. Russell, J.C., et al., Generation and characterization of a tractable C. elegans model of tauopathy. Geroscience, 2021. 43(5): p. 2621–2631.

52. Lam, A.B., K. Kervin, and J.E. Tanis, Vitamin B(12) impacts amyloid beta-induced proteotoxicity by regulating the methionine/S-adenosylmethionine cycle. Cell Rep, 2021. 36(13): p. 109753.

