## Supplement figure for "Fluphenazine, an antipsychotic compound, ameliorates Alzheimer’s disease by clearing amyloid beta accumulation in *C. elegans*"

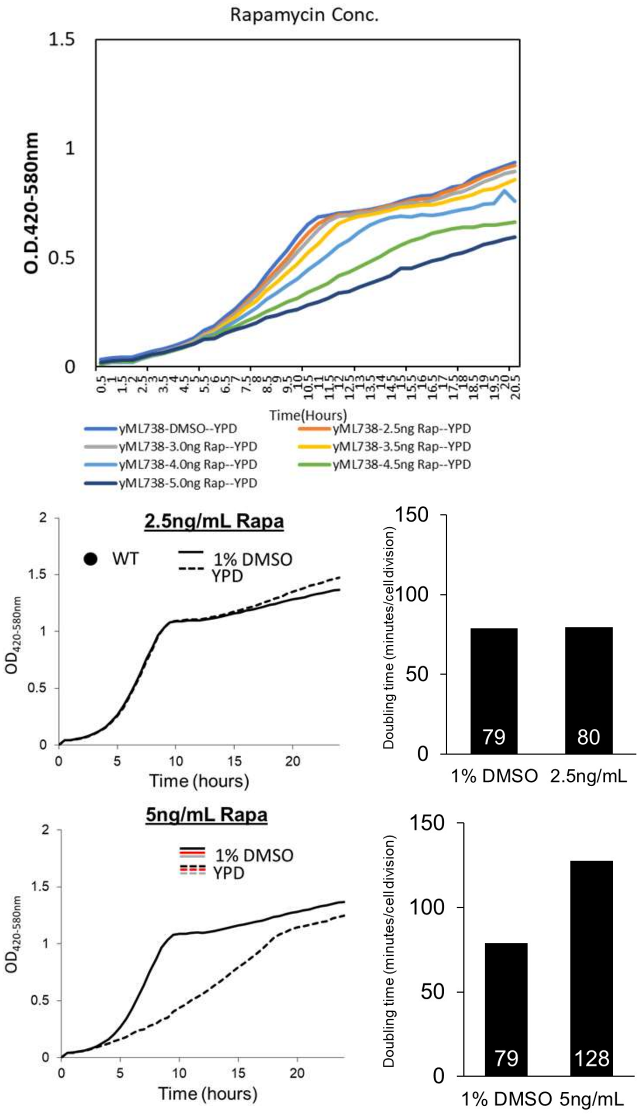


**Figure S1. Determination of rapamycin concentration that does not affect yeast growth.**

*BY4741* strain was treated or not with increasing concentration of rapamycin from 1ng/ml to 5 ng/ml. 2.5ng/ml Rapamycin did not affect the doubling time of the yeast growth; therefore this conc. was selected to use in combination with FDA approved compound.

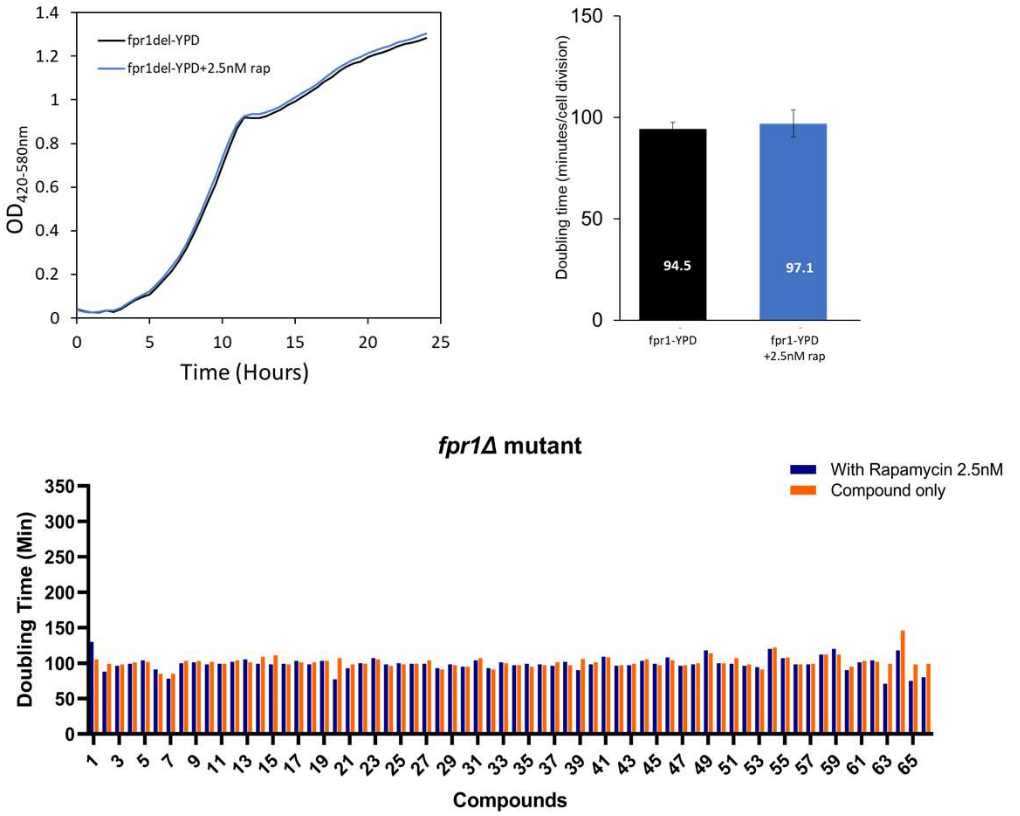


**Figure S2.** ***fpr1*** **deletion abolished synergistic effect with rapamycin.** *fpr1*Δ strain was treated or not with 2.5ng/ml conc. of rapamycin which did not affect the doubling time of the yeast growth. *fpr1*Δ strain was treated or not with 2.5ng/ml rapamycin with 10uM Positive hits compound panel.
