## Supplement table 1 for "Fluphenazine, an antipsychotic compound, ameliorates Alzheimer’s disease by clearing amyloid beta accumulation in *C. elegans*"

| | | FDA library compound | | | |
|---|---|---|---|---|---|
| **No.** | **Compound name** | **doubling time inflection (min) with 2.5ng/ml Rap RUN 1** | **doubling time inflection (min) with 2.5ng/ml Rap RUN 2** | **doubling time inflection (min) with DMSO RUN 1** | **doubling time inflection (min) with DMSO RUN 2** |
| 1 | GRISEOFULVIN | 91.6 | 80.5 | 80.4 | 86.0 |
| 2 | TESTOSTERONE | 86.2 | 77.9 | 80.1 | 80.2 |
| 3 | ALLOPURINOL | 90.1 | 76.1 | 79.3 | 84.1 |
| 4 | AMODIAQUINE DIHYDROCHLORIDE | 84.3 | 78.6 | 80.3 | 75.8 |
| 5 | BACITRACIN | 85.0 | 76.5 | 78.3 | 72.6 |
| 6 | BITHIONOL | 280.6 | 147.0 | 133.2 | 81.7 |
| 7 | CEFADROXIL | 84.2 | 77.2 | 79.4 | 82.5 |
| 8 | CHLORAMPHENICOL SODIUM SUCCINATI | 88.2 | 74.9 | 78.9 | 83.9 |
| 9 | SALSALATE | 88.0 | 78.3 | 79.5 | 82.7 |
| 10 | SANGUINARIUM CHLORIDE | 89.4 | 81.3 | 82.0 | 79.4 |
| 11 | ALVERINE CITRATE | 101.8 | 78.1 | 76.9 | 76.5 |
| 12 | AMOXICILLIN | 84.3 | 76.5 | 77.6 | 78.5 |
| 13 | BECLOMETHASONE DIPROPIONATE | 80.4 | 75.6 | 77.8 | 85.7 |
| 14 | BROMOCRIPTINE MESYLATE | 88.6 | 73.2 | 74.7 | 68.3 |
| 15 | CEFAZOLIN SODIUM | 86.9 | 75.5 | 73.2 | 76.7 |
| 16 | CHLORAMPHENICOL | 90.1 | 77.3 | 77.0 | 83.2 |
| 17 | DANTHRON | 100.9 | 84.0 | 96.4 | 94.2 |
| 18 | MITOMYCIN | 88.9 | 80.5 | 80.9 | 82.7 |
| 19 | AMANTADINE HYDROCHLORIDE | 86.4 | 78.7 | 78.6 | 77.5 |
| 20 | AMPHOTERICIN B | 193.6 | 111.0 | 94.4 | 103.2 |
| 21 | BENSERAZIDE HYDROCHLORIDE | 84.2 | 79.9 | 78.6 | 76.1 |
| 22 | BUSULFAN | 79.1 | 80.1 | 75.8 | 80.9 |
| 23 | CEFOTAXIME SODIUM | 79.3 | 72.3 | 78.4 | 79.2 |
| 24 | NORGESTIMATE | 85.6 | 80.7 | 82.0 | 78.2 |
| 25 | MEQUINOL | 86.6 | 78.3 | 77.6 | 78.6 |
| 26 | SODIUM NITROPRUSSIDE DIHYDRATE | 86.4 | 80.6 | 78.3 | 77.8 |
| 27 | LEVOMILNACIPRAN HYDROCHLORIDE | 86.8 | 80.2 | 79.4 | 82.3 |
| 28 | AMPICILLIN SODIUM | 88.4 | 76.4 | 73.6 | 76.4 |
| 29 | BENZETHONIUM CHLORIDE | 0.0 | 0.0 | 170.8 | 139.7 |
| 30 | CAFFEINE | 84.5 | 78.7 | 77.9 | 75.3 |
| 31 | CEPHALOTHIN SODIUM | 83.8 | 77.9 | 79.3 | 78.8 |
| 32 | CHLORHEXIDINE DIHYDROCHLORIDE | | | | |
| 33 | HYDROCORTISONE | 81.0 | 83.0 | 77.0 | 82.9 |
| 34 | SODIUM OXYBATE | 85.0 | 77.4 | 75.7 | 77.7 |
| 35 | AMILORIDE HYDROCHLORIDE | 80.3 | 75.6 | 74.2 | 80.4 |
| 36 | ANTHRALIN | 86.5 | 84.0 | 81.9 | 81.8 |
| 37 | BENZOCAINE | 79.8 | 75.1 | 80.5 | 78.4 |
| 38 | CAMPHOR | 79.8 | 72.3 | 78.8 | 76.9 |
| 39 | CEPHAPIRIN SODIUM | 80.1 | 70.9 | 70.6 | 77.4 |
| 40 | CHLOROCRESOL | 87.4 | 78.8 | 80.4 | 77.9 |
| 41 | DESOXYCORTICOSTERONE ACETATE | 85.5 | 78.8 | 78.3 | 72.9 |
| 42 | MANNITOL | 81.2 | 76.0 | 78.6 | 75.4 |
| 43 | POTASSIUM p-AMINOBENZOATE | 83.5 | 75.5 | 78.3 | 72.2 |
| 44 | ANTIPYRINE | 85.4 | 78.2 | 75.6 | 79.8 |
| 45 | BENZTHIAZIDE | 81.7 | 74.1 | 79.8 | 73.9 |
| 46 | CARBACHOL | 84.0 | 76.3 | 77.0 | 76.8 |
| 47 | ACETAZOLAMIDE | 79.8 | 75.1 | 80.5 | 76.8 |
| 48 | CHLOROTHIAZIDE | 83.5 | 76.4 | 73.7 | 76.6 |
| 49 | TESTOSTERONE PROPIONATE | 90.4 | 84.0 | 81.2 | 71.3 |
| 50 | ACETAMINOPHEN | 81.0 | 79.8 | 80.8 | 68.3 |
| 51 | AMINOCAPROIC ACID HYDROCHLORIDE | 81.2 | 74.5 | 76.7 | 64.3 |
| 52 | APOMORPHINE HYDROCHLORIDE | 84.0 | 79.7 | 78.3 | 76.6 |
| 53 | CARBOPLATIN | 86.8 | 73.3 | 76.7 | 79.5 |
| 54 | CARBAMAZEPINE | 81.5 | 79.6 | 72.6 | 76.9 |
| 55 | CEPHRADINE | 80.4 | 75.7 | 78.6 | 79.5 |
| 56 | CHLOROXYLENOL | 116.4 | 88.0 | 80.1 | 78.6 |
| 57 | SPARTEINE SULFATE | 83.2 | 80.3 | 80.3 | 79.7 |
| 58 | ACETYLCHOLINE CHLORIDE | 80.1 | 74.6 | 78.3 | 76.6 |
| 59 | AMINOGLUTETHIMIDE | 79.8 | 75.9 | 79.5 | 73.8 |
| 60 | ASPIRIN | 70.9 | 76.8 | 75.7 | 74.2 |
| 61 | BETAMETHASONE VALERATE | 80.1 | 78.7 | 74.6 | 78.9 |
| 62 | CARBENICILLIN DISODIUM | 77.5 | 77.4 | 75.2 | 76.2 |
| 63 | CETYLPYRIDINIUM CHLORIDE | | | | |
| 64 | DEXCHLORPHENIRAMINE MALEATE | 77.1 | 75.5 | 78.9 | 77.2 |
| 65 | SALICYLANILIDE | 91.3 | 87.7 | 82.2 | 84.6 |
| 66 | ACETYLCYSTEINE | 78.1 | 75.5 | 80.2 | 75.4 |

| | | | | | |
|---|---|---|---|---|---|
| 67 | AMINOSALICYLATE SODIUM | 77.1 | 74.5 | 77.3 | 78.9 |
| 68 | ATROPINE SULFATE | 78.8 | 79.9 | 76.6 | 75.1 |
| 69 | BETHANECHOL CHLORIDE | 78.6 | 77.3 | 74.7 | 76.5 |
| 70 | CARBINOXAMINE MALEATE | 77.0 | 78.6 | 74.6 | 76.0 |
| 71 | CHLORAMBUCIL | 75.7 | 78.5 | 78.1 | 75.2 |
| 72 | CHLORPROMAZINE HYDROCHLORIDE | 175.7 | 128.8 | 83.2 | 89.5 |
| 73 | PHENYL AMINOSALICYLATE | 80.9 | 76.7 | 80.1 | 79.1 |
| 74 | ADENOSINE | 79.4 | 78.8 | 73.2 | 75.7 |
| 75 | AMITRIPTYLINE HYDROCHLORIDE | 111.7 | 84.7 | 77.4 | 77.4 |
| 76 | AZATHIOPRINE | 77.6 | 77.7 | 74.7 | 76.7 |
| 77 | BISACODYL | 77.5 | 74.4 | 74.2 | 77.2 |
| 78 | CARISOPRODOL | 77.8 | 74.8 | 77.3 | 77.4 |
| 79 | CHLORAMPHENICOL PALMITATE | 74.1 | 77.6 | 80.9 | 82.1 |
| 80 | CHLORPROPAMIDE | 71.8 | 79.1 | 78.0 | 77.0 |
| 81 | CHLORTETRACYCLINE HYDROCHLORIDE | 86.2 | 96.5 | 87.5 | 85.7 |
| 82 | CLOTRIMAZOLE | 265.7 | | | 270.2 |
| 83 | CYCLOPENTOLATE HYDROCHLORIDE | 84.5 | 95.6 | 84.7 | 76.5 |
| 84 | DEHYDROCHOLATE SODIUM | 88.5 | 88.8 | 87.0 | 78.4 |
| 85 | DICLOXACILLIN SODIUM | 88.6 | 93.7 | 85.3 | 79.1 |
| 86 | DIMERCAPROL | 92.7 | 89.8 | 83.4 | 78.4 |
| 87 | DOXYCYCLINE HYDROCHLORIDE | 87.4 | 95.3 | 90.8 | 80.0 |
| 88 | ERGONOVINE MALEATE | 92.2 | 95.9 | 80.9 | 80.3 |
| 89 | CHLORTHALIDONE | 91.5 | 94.2 | 82.3 | 78.9 |
| 90 | CLOXACILLIN SODIUM | 81.5 | 92.7 | 83.0 | 79.3 |
| 91 | CYCLOPHOSPHAMIDE | 86.6 | 89.1 | 80.7 | 77.9 |
| 92 | DEMECLOCYCLINE HYDROCHLORIDE | 84.6 | 93.2 | 81.3 | 79.7 |
| 93 | DICYCLOMINE HYDROCHLORIDE | 101.2 | 112.0 | 84.6 | 77.9 |
| 94 | DIMETHADIONE | 91.7 | 93.2 | 81.6 | 76.6 |
| 95 | DOXYLAMINE SUCCINATE | 91.0 | 90.6 | 80.5 | 78.3 |
| 96 | ERYTHROMYCIN ETHYLSUCCINATE | 87.6 | 95.3 | 83.9 | 79.5 |
| 97 | CHLORZOXAZONE | 90.7 | 94.8 | 85.3 | 80.7 |
| 98 | CLOXYQUIN | 115.6 | 129.5 | 99.0 | 93.7 |
| 99 | CYCLOSERINE (D) | 89.1 | 81.1 | 80.9 | 80.1 |
| 100 | DESIPRAMINE HYDROCHLORIDE | 107.1 | 113.3 | 84.4 | 78.7 |
| 101 | DIENESTROL | 87.0 | 85.5 | 81.8 | 76.4 |
| 102 | DIOXYBENZONE | 86.0 | 91.7 | 83.9 | 80.9 |
| 103 | DYCLONINE HYDROCHLORIDE | 127.6 | 123.6 | 89.0 | 74.3 |
| 104 | ERYTHROMYCIN | 89.3 | 89.2 | 84.3 | 83.9 |
| 105 | CICLOPIROX OLAMINE | 115.5 | 127.6 | 87.0 | 81.4 |
| 106 | COLCHICINE | 87.1 | 89.9 | 81.8 | 77.0 |
| 107 | CYPROTERONE ACETATE | 86.5 | 88.5 | 82.0 | 76.2 |
| 108 | DEXAMETHASONE | 85.3 | 91.7 | 80.8 | 76.4 |
| 109 | DIETHYLCARBAMAZINE CITRATE | 92.7 | 89.6 | 83.7 | 79.0 |
| 110 | DIPHENHYDRAMINE HYDROCHLORIDE | 86.9 | 90.1 | 80.0 | 76.6 |
| 111 | DYPHYLLINE | 83.7 | 90.1 | 77.5 | 79.1 |
| 112 | ESTRADIOL | 91.5 | 99.9 | 79.1 | 80.0 |
| 113 | CINOXACIN | 85.5 | 91.7 | 85.8 | 67.2 |
| 114 | COLISTIMETHATE SODIUM | 98.4 | 117.0 | 80.1 | 80.1 |
| 115 | CYTARABINE | 75.9 | 90.5 | 82.1 | 77.9 |
| 116 | DEXAMETHASONE ACETATE | 90.3 | 91.1 | 80.4 | 80.4 |
| 117 | DIETHYLSTILBESTROL | 88.8 | 85.7 | 81.5 | 78.2 |
| 118 | DIPHENYLPYRALINE HYDROCHLORIDE | 71.3 | 88.5 | 79.8 | 73.7 |
| 119 | ETHYLENEDIAMINE TETRACETIC ACID | 87.2 | 88.1 | 83.6 | 79.1 |
| 120 | ESTRADIOL CYPIONATE | 78.0 | 87.6 | 81.8 | 75.3 |
| 121 | CLEMASTINE FUMARATE | 102.9 | 114.8 | 79.8 | 80.2 |
| 122 | CORTISONE ACETATE | 80.3 | 91.2 | 77.8 | 79.2 |
| 123 | DACARBAZINE | 93.2 | 83.3 | 80.4 | 77.6 |
| 124 | DEXAMETHASONE SODIUM PHOSPHATE | 91.2 | 85.4 | 83.8 | 76.6 |
| 125 | DIFLUNISAL | 95.7 | 92.1 | 79.4 | 77.3 |
| 126 | DIPYRIDAMOLE | 86.2 | 92.1 | 82.3 | 76.9 |
| 127 | EMETINE DIHYDROCHLORIDE | 88.7 | 91.6 | 80.8 | 78.2 |
| 128 | ESTRADIOL VALERATE | 81.1 | 82.7 | 79.9 | 77.4 |
| 129 | CLIDINIUM BROMIDE | 92.6 | 94.8 | 82.2 | 66.1 |
| 130 | COTININE | 82.8 | 93.2 | 79.9 | 77.8 |
| 131 | DANAZOL | 84.4 | 89.4 | 84.0 | 76.5 |
| 132 | DEXTROMETHORPHAN HYDROBROMIDE | 94.9 | 99.9 | 79.6 | 76.6 |
| 133 | DIGITOXIN | 191.8 | 221.6 | 162.2 | 123.9 |
| 134 | PYRITHIONE ZINC | 378.7 | | | 237.3 |
| 135 | EPHEDRINE (1R,2S) HYDROCHLORIDE | 87.6 | 93.2 | 82.7 | 77.1 |
| 136 | ESTRIOL | 90.8 | 90.5 | 82.1 | 77.8 |
| 137 | CLINDAMYCIN HYDROCHLORIDE | 88.8 | 93.2 | 78.1 | 69.1 |
| 138 | CRESOL | 91.2 | 91.1 | 80.4 | 73.9 |

| | | | | | |
|---|---|---|---|---|---|
| 139 | DAPSONE | 88.3 | 90.5 | 80.1 | 78.8 |
| 140 | DIBENZOTHIOPHENE | 122.4 | 111.7 | 82.7 | 80.6 |
| 141 | DIGOXIN | 90.2 | 87.7 | 78.7 | 76.9 |
| 142 | DISULFIRAM | 125.7 | 125.8 | 78.3 | 89.8 |
| 143 | EPINEPHRINE BITARTRATE | 91.0 | 87.2 | 82.0 | 79.8 |
| 144 | ESTRONE | 109.4 | 111.9 | 80.9 | 80.4 |
| 145 | CLOMIPHENE CITRATE | 189.3 | 202.6 | 112.7 | 99.4 |
| 146 | CROMOLYN SODIUM | 85.2 | 87.6 | 81.7 | 73.3 |
| 147 | DAUNORUBICIN HYDROCHLORIDE | 94.3 | 89.3 | 81.2 | 81.2 |
| 148 | DIBUCAINE HYDROCHLORIDE | 98.4 | 101.4 | 80.2 | 77.7 |
| 149 | DIHYDROERGOTAMINE MESYLATE | 97.7 | 90.7 | 78.5 | 78.5 |
| 150 | DOPAMINE HYDROCHLORIDE | 89.2 | 86.4 | 79.7 | 78.7 |
| 151 | DOMPERIDONE | 84.7 | 87.0 | 79.5 | 77.2 |
| 152 | ETHACRYNIC ACID | 92.7 | 90.2 | 81.1 | 76.6 |
| 153 | CLONIDINE HYDROCHLORIDE | 86.4 | 90.6 | 82.3 | 75.2 |
| 154 | CYCLIZINE | 91.2 | 90.1 | 79.7 | 77.7 |
| 155 | DEFEROXAMINE MESYLATE | 90.3 | 87.6 | 75.7 | 78.7 |
| 156 | DICLOFENAC SODIUM | 88.9 | 86.1 | 80.8 | 78.5 |
| 157 | DIMENHYDRINATE | 80.9 | 92.2 | 84.6 | 79.0 |
| 158 | DOXEPIN HYDROCHLORIDE | 81.4 | 85.6 | 80.2 | 78.5 |
| 159 | ERGOCALCIFEROL | 84.7 | 86.2 | 83.0 | 78.2 |
| 160 | ETHAMBUTOL HYDROCHLORIDE | 81.9 | 85.3 | 78.4 | 78.8 |
| 161 | ETHINYL ESTRADIOL | 89.1 | 68.0 | 87.5 | 85.8 |
| 162 | DORAMECTIN | 101.8 | 95.1 | 80.9 | 81.0 |
| 163 | GRAMICIDIN (gramicidin A shown) | 87.5 | 76.0 | 82.5 | 77.6 |
| 164 | HOMATROPINE METHYLBROMIDE | 87.0 | 62.6 | 80.8 | 81.5 |
| 165 | IBUPROFEN | 94.6 | 68.6 | 82.2 | 82.4 |
| 166 | ISOPROTERENOL HYDROCHLORIDE | 90.1 | 71.8 | 76.7 | 84.2 |
| 167 | MAPROTILINE HYDROCHLORIDE | 99.5 | 90.1 | 89.4 | 84.7 |
| 168 | MERCAPTOPURINE | 86.6 | 92.1 | 79.1 | 84.7 |
| 169 | ETHIONAMIDE | 83.3 | 80.4 | 86.9 | 81.3 |
| 170 | GLUTATHIONE | 85.5 | 83.8 | 75.6 | 83.4 |
| 171 | GUAIFENESIN | 89.5 | 84.5 | 63.5 | 83.4 |
| 172 | HYDROCHLOROTHIAZIDE | 91.6 | 89.8 | 84.3 | 83.0 |
| 173 | IMIPRAMINE HYDROCHLORIDE | 94.4 | 97.6 | 85.5 | 78.7 |
| 174 | ISOSORBIDE DINITRATE | 84.0 | 79.3 | 79.6 | 81.3 |
| 175 | MECAMYLAMINE HYDROCHLORIDE | 88.2 | 66.4 | 86.5 | 83.2 |
| 176 | MESTRANOL | 93.2 | 69.8 | 68.3 | 82.9 |
| 177 | ETHOPROPAZINE HYDROCHLORIDE | 138.7 | 99.5 | 90.4 | 84.8 |
| 178 | FLURBIPROFEN | 88.9 | 82.5 | 76.9 | 83.6 |
| 179 | GUANABENZ ACETATE | 89.6 | 83.5 | 80.1 | 78.5 |
| 180 | HYDROCORTISONE ACETATE | 87.1 | 52.6 | 79.1 | 77.2 |
| 181 | INDAPAMIDE | 87.7 | 80.7 | 87.7 | 83.0 |
| 182 | ISOXSUPRINE HYDROCHLORIDE | 92.0 | 76.6 | 74.2 | 84.0 |
| 183 | MECHLORETHAMINE | 91.6 | 87.7 | 90.4 | 53.1 |
| 184 | METAPROTERENOL | 92.1 | 85.2 | 50.9 | 79.5 |
| 185 | EUCALYPTOL | 88.7 | 82.7 | 85.2 | 78.2 |
| 186 | FURAZOLIDONE | 93.2 | 80.9 | 88.5 | 83.5 |
| 187 | HALAZONE | 93.2 | 77.1 | 85.5 | 84.0 |
| 188 | HYDROCORTISONE HEMISUCCINATE | 88.5 | 82.5 | 66.6 | 83.3 |
| 189 | INDOMETHACIN | 94.9 | 90.8 | 89.6 | 82.4 |
| 190 | KANAMYCIN A SULFATE | 87.4 | 79.6 | 86.5 | 80.0 |
| 191 | MECLIZINE HYDROCHLORIDE | 104.3 | 56.4 | 84.2 | 85.9 |
| 192 | METHACHOLINE CHLORIDE | 90.6 | 53.4 | 84.9 | 81.4 |
| 193 | SEMUSTINE | 87.8 | 54.1 | 80.3 | 79.0 |
| 194 | FUROSEMIDE | 90.6 | 82.0 | 50.6 | 79.5 |
| 195 | HALOPERIDOL | 169.6 | 114.8 | 87.5 | 92.6 |
| 196 | HYDROCORTISONE PHOSPHATE TRIETHYI | 89.6 | 78.8 | 84.2 | 78.5 |
| 197 | CANAGLIFLOZIN | 103.3 | 104.0 | 82.7 | 85.5 |
| 198 | KETOCONAZOLE | 242.8 | 140.4 | 141.8 | 162.2 |
| 199 | MECLOFENAMATE SODIUM | 86.2 | 52.0 | 87.2 | 82.9 |
| 200 | METHENAMINE | 79.7 | 69.5 | 81.9 | 83.7 |
| 201 | EUGENOL | 84.7 | 87.5 | 78.3 | 77.0 |
| 202 | FUSIDIC ACID | 90.7 | 75.7 | 83.6 | 83.0 |
| 203 | HETACILLIN POTASSIUM | 87.6 | 81.2 | 52.6 | 81.4 |
| 204 | HYDROFLUMETHIAZIDE | 86.1 | 79.4 | 84.0 | 80.7 |
| 205 | INOSITOL | 91.5 | 53.1 | 85.2 | 82.1 |
| 206 | LACTULOSE | 89.6 | 88.9 | 86.2 | 82.5 |
| 207 | MEDROXYPROGESTERONE ACETATE | 90.3 | 72.8 | 85.4 | 82.7 |
| 208 | METHICILLIN SODIUM | 92.7 | 79.1 | 86.7 | 79.0 |
| 209 | FLUDROCORTISONE ACETATE | 81.6 | 77.4 | 81.5 | 77.9 |
| 210 | GALLAMINE TRIETHIODIDE | 85.5 | 54.9 | 75.4 | 83.4 |

| | | | | | |
|---|---|---|---|---|---|
| 211 | HEXACHLOROPHENE | 102.9 | 115.4 | 108.6 | 137.5 |
| 212 | HYDROXYPROGESTERONE CAPROATE | 88.1 | 88.0 | 86.1 | 80.9 |
| 213 | IODOQUINOL | 82.0 | 96.5 | 87.1 | 85.7 |
| 214 | LEUCOVORIN CALCIUM | 84.4 | 88.2 | 88.7 | 79.7 |
| 215 | MEDRYSONE | 79.1 | 55.4 | 49.3 | 80.4 |
| 216 | METHIMAZOLE | 84.4 | 67.8 | 70.4 | 73.6 |
| 217 | FLUMETHAZONE PIVALATE | 82.7 | 81.9 | 76.7 | 84.3 |
| 218 | GEMFIBROZIL | 83.4 | 67.4 | 86.5 | 84.0 |
| 219 | HEXYLRESORCINOL | 95.4 | 95.0 | 93.9 | 88.7 |
| 220 | HYDROXYUREA | 79.9 | 81.7 | 89.0 | 82.1 |
| 221 | IPRATROPIUM BROMIDE | 76.7 | 76.1 | 83.8 | 82.2 |
| 222 | LEVONORDEFRIN | 78.5 | 51.3 | 84.0 | 68.8 |
| 223 | MEGESTROL ACETATE | 78.3 | 75.0 | 83.4 | 82.7 |
| 224 | METHOCARBAMOL | 78.2 | 78.0 | 71.8 | 82.6 |
| 225 | FLUOCINOLONE ACETONIDE | 92.5 | 83.3 | 78.0 | 85.2 |
| 226 | GENTIAN VIOLET | 417.3 | 474.7 | 447.0 | 551.0 |
| 227 | HISTAMINE DIHYDROCHLORIDE | 87.4 | 78.9 | 70.1 | 84.8 |
| 228 | HYDROXYZINE PAMOATE | 83.0 | 55.3 | 87.0 | 81.3 |
| 229 | ISONIAZID | 85.4 | 53.4 | 74.7 | 80.4 |
| 230 | LINCOMYCIN HYDROCHLORIDE | 79.3 | 51.6 | 75.8 | 84.2 |
| 231 | MELPHALAN | 74.7 | 78.8 | 83.2 | 82.2 |
| 232 | METHOTREXATE HYDRATE | 80.9 | 81.5 | 83.4 | 84.1 |
| 233 | FLUOCINONIDE | 81.7 | 79.8 | 82.5 | 78.2 |
| 234 | GLUCOSAMINE HYDROCHLORIDE | 81.9 | 63.7 | 49.6 | 83.5 |
| 235 | HOMATROPINE HYDROBROMIDE | 83.2 | 68.2 | 48.6 | 84.1 |
| 236 | HYOSCYAMINE | 79.4 | 74.1 | 89.8 | 83.6 |
| 237 | ISOPROPAMIDE IODIDE | 80.1 | 74.1 | 75.2 | 76.7 |
| 238 | MAFENIDE HYDROCHLORIDE | 81.0 | 71.0 | 62.4 | 81.2 |
| 239 | MEPENZOLATE BROMIDE | 80.7 | 66.6 | 60.4 | 81.3 |
| 240 | METHOXAMINE HYDROCHLORIDE | 80.7 | 76.3 | 84.9 | 79.0 |
| 241 | METHOXSALEN | 91.6 | 104.4 | | 84.9 |
| 242 | MINOCYCLINE HYDROCHLORIDE | 88.1 | 100.6 | | 78.2 |
| 243 | NIFEDIPINE | 84.8 | 96.0 | | 83.8 |
| 244 | NORGESTREL | 90.1 | 101.5 | | 80.8 |
| 245 | OXYPHENBUTAZONE | 92.2 | 103.1 | | 80.8 |
| 246 | PHENELZINE SULFATE | 94.4 | 97.4 | | 83.5 |
| 247 | PINDOLOL | 92.6 | 100.4 | | 82.7 |
| 248 | PRIMAQUINE PHOSPHATE | 100.4 | 115.5 | | 81.5 |
| 249 | METHSCOPOLAMINE BROMIDE | 83.0 | 102.0 | | 80.2 |
| 250 | MOXALACTAM DISODIUM | 89.2 | 103.4 | | 80.8 |
| 251 | NITROFURANTOIN | 74.9 | 102.0 | | 80.6 |
| 252 | NOSCAPINE HYDROCHLORIDE | 87.2 | 97.1 | | 79.5 |
| 253 | OXYQUINOLINE HEMISULFATE | 96.2 | 117.3 | | 81.7 |
| 254 | PHENINDIONE | 88.6 | 95.6 | | 79.2 |
| 255 | PIPERACILLIN SODIUM | 92.1 | 98.7 | | 81.3 |
| 256 | PRIMIDONE | 88.0 | 98.1 | | 81.5 |
| 257 | METHYLDOPA | 90.6 | 106.5 | | 82.7 |
| 258 | NADIDE | 84.5 | 105.5 | | 81.8 |
| 259 | NITROFURAZONE | 90.2 | 102.3 | | 81.9 |
| 260 | NOVOBIOCIN SODIUM | 92.0 | 98.2 | | 69.0 |
| 261 | OXYTETRACYCLINE | 89.6 | 94.0 | | 82.3 |
| 262 | PHENIRAMINE MALEATE | 91.6 | 101.0 | | 78.9 |
| 263 | PIPERAZINE | 90.0 | 99.1 | | 78.4 |
| 264 | PROBENECID | 91.1 | 101.6 | | 82.9 |
| 265 | DEFERASIROX | 86.0 | 104.1 | | 81.9 |
| 266 | NAFCILLIN SODIUM | 81.8 | 95.1 | | 80.6 |
| 267 | NITROMIDE | 81.8 | 97.1 | | 80.0 |
| 268 | NYLIDRIN HYDROCHLORIDE | 88.2 | 104.5 | | 80.0 |
| 269 | PAPAVERINE HYDROCHLORIDE | 87.8 | 99.2 | | 80.8 |
| 270 | PHENOLPHTHALEIN | 83.6 | 93.2 | | 80.8 |
| 271 | PIROXICAM | 79.3 | 98.3 | | 82.0 |
| 272 | PROCAINAMIDE HYDROCHLORIDE | 88.0 | 102.2 | | 79.4 |
| 273 | METHYLPREDNISOLONE | 87.5 | 99.5 | | 84.2 |
| 274 | PHENOXYBENZAMINE HYDROCHLORIDE | 84.9 | 104.6 | | 83.0 |
| 275 | NOREPINEPHRINE TARTRATE | 85.3 | 99.7 | | 82.7 |
| 276 | NYSTATIN | 89.2 | 112.6 | | 91.4 |
| 277 | PARACHLOROPHENOL | 86.1 | 97.2 | | 83.9 |
| 278 | PHENYLBUTAZONE | 84.0 | 93.8 | | 81.7 |
| 279 | POLYMYXIN B SULFATE | 273.7 | 315.3 | | 80.0 |
| 280 | PROCAINE HYDROCHLORIDE | 82.0 | 88.8 | | 78.4 |
| 281 | METHYLTHIOURACIL | 81.5 | 97.3 | | 82.5 |
| 282 | NAPHAZOLINE HYDROCHLORIDE | 81.2 | 98.6 | | 82.5 |

| | | | | | |
|---|---|---|---|---|---|
| 283 | NORETHINDRONE | 78.1 | 102.3 | | 80.6 |
| 284 | ORPHENADRINE CITRATE | 84.3 | 106.9 | | 80.9 |
| 285 | PARGYLINE HYDROCHLORIDE | 278.7 | 231.4 | | 82.0 |
| 286 | L-PHENYLEPHRINE HYDROCHLORIDE | 82.7 | 100.3 | | 79.1 |
| 287 | PRAZIQUANTEL | 82.9 | 96.3 | | 82.0 |
| 288 | PROCHLORPERAZINE EDISYLATE | 329.1 | | | 145.9 |
| 289 | METOCLOPRAMIDE HYDROCHLORIDE | 94.2 | 106.0 | | 82.2 |
| 290 | NAPROXEN | 84.9 | 102.0 | | 81.5 |
| 291 | NORETHINDRONE ACETATE | 83.7 | 95.0 | | 80.8 |
| 292 | OXACILLIN SODIUM | 83.0 | 99.8 | | 81.0 |
| 293 | PENICILLIN G POTASSIUM | 87.4 | 96.3 | | 83.5 |
| 294 | 1S,2R-PHENYLPROPANOLAMINE HYDROC | 88.6 | 98.5 | | 82.5 |
| 295 | KITASAMYCINS [A1 shown] | 82.5 | 99.2 | | 79.2 |
| 296 | PROCYCLIDINE HYDROCHLORIDE | 89.0 | 86.2 | | 78.3 |
| 297 | METOPROLOL TARTRATE | 83.5 | 99.8 | | 80.1 |
| 298 | NEOMYCIN TRISULFATE | 80.5 | 93.2 | | 81.8 |
| 299 | NORTRIPTYLINE HYDROCHLORIDE | 114.9 | 162.2 | | 80.5 |
| 300 | OXIDOPAMINE HYDROCHLORIDE | 89.2 | 105.2 | | 84.0 |
| 301 | PENICILLIN V POTASSIUM | 83.5 | 94.7 | | 80.2 |
| 302 | PHENYTOIN SODIUM | 84.1 | 98.2 | | 82.0 |
| 303 | PREDNISOLONE | 81.7 | 97.6 | | 81.1 |
| 304 | PROGESTERONE | 92.1 | 114.1 | | 82.2 |
| 305 | METRONIDAZOLE | 84.2 | 101.3 | | 79.4 |
| 306 | NEOSTIGMINE BROMIDE | 79.6 | 96.9 | | 82.0 |
| 307 | NORETHYNODREL | 80.4 | 102.5 | | 79.9 |
| 308 | OXYBENZONE | 81.4 | 96.2 | | 86.3 |
| 309 | PHENACEMIDE | 84.2 | 100.8 | | 82.4 |
| 310 | APIXABAN | 85.3 | 102.3 | | 84.7 |
| 311 | PREDNISOLONE ACETATE | 80.7 | 98.7 | | 79.8 |
| 312 | PROMAZINE HYDROCHLORIDE | 92.7 | 116.7 | | 86.0 |
| 313 | MICONAZOLE NITRATE | 551.0 | 1091.7 | | 130.2 |
| 314 | NIACIN | 82.0 | 94.5 | | 81.7 |
| 315 | NORFLOXACIN | 83.6 | 100.5 | | 78.9 |
| 316 | OXYMETAZOLINE HYDROCHLORIDE | 78.3 | 93.2 | | 78.0 |
| 317 | PHENAZOPYRIDINE HYDROCHLORIDE | 95.8 | 114.1 | | 93.0 |
| 318 | PILOCARPINE NITRATE | 82.7 | 99.5 | | 79.0 |
| 319 | PREDNISONE | 78.0 | 100.1 | | 79.2 |
| 320 | PROMETHAZINE HYDROCHLORIDE | 89.2 | 122.0 | | 83.0 |
| 321 | PROPANTHELINE BROMIDE | 82.7 | 99.3 | | 85.8 |
| 322 | QUINIDINE GLUCONATE | 85.2 | 94.4 | | 77.3 |
| 323 | SCOPOLAMINE HYDROBROMIDE | 83.7 | 95.7 | | 83.6 |
| 324 | SULFAMERAZINE | 79.3 | 94.6 | | 83.2 |
| 325 | TAMOXIFEN CITRATE | 92.6 | 107.5 | | 81.9 |
| 326 | THIOTHIXENE | 90.4 | 101.8 | | 82.9 |
| 327 | TRIAMCINOLONE ACETONIDE | 83.7 | 91.2 | | 79.9 |
| 328 | TRIPELENNAMINE CITRATE | 81.6 | 89.4 | | 79.0 |
| 329 | DEXPROPRANOLOL HYDROCHLORIDE [R(- | 84.2 | 101.0 | | 84.1 |
| 330 | QUININE | 84.6 | 96.1 | | 74.6 |
| 331 | SALICYLIC ACID | 86.4 | 95.1 | | 75.7 |
| 332 | SULFAMETHAZINE | 78.9 | 92.6 | | 82.6 |
| 333 | TERBUTALINE HEMISULFATE | 88.4 | 94.4 | | 76.7 |
| 334 | TIMOLOL MALEATE | 79.8 | 95.1 | | 79.6 |
| 335 | BETAMETHASONE | 83.5 | 96.1 | | 82.1 |
| 336 | TRIPROLIDINE HYDROCHLORIDE | 86.6 | 95.8 | | 78.9 |
| 337 | PROPYLTHIOURACIL | 84.4 | 95.1 | | 76.3 |
| 338 | RACEPHEDRINE HYDROCHLORIDE | 83.4 | 98.1 | | 83.5 |
| 339 | RIBAVIRIN | 82.4 | 93.2 | | 83.7 |
| 340 | SULFAMETHIZOLE | 84.0 | 94.7 | | 81.9 |
| 341 | TETRACAINE HYDROCHLORIDE | 83.8 | 99.9 | | 79.2 |
| 342 | TOBRAMYCIN | 83.5 | 94.5 | | 76.9 |
| 343 | TRIAMTERENE | 74.8 | 96.9 | | 75.7 |
| 344 | TROPICAMIDE | 87.8 | 97.4 | | 78.7 |
| 345 | beta-CAROTENE [2mM] | 85.7 | 95.2 | | 71.8 |
| 346 | RESERPINE | 86.2 | 93.7 | | 75.6 |
| 347 | SPECTINOMYCIN HYDROCHLORIDE | 82.0 | 90.2 | | 74.1 |
| 348 | SULFAMETHOXAZOLE | 83.7 | 93.2 | | 72.7 |
| 349 | TETRACYCLINE HYDROCHLORIDE | 83.0 | 91.9 | | 76.0 |
| 350 | TOLAZOLINE HYDROCHLORIDE | 81.2 | 94.7 | | 74.1 |
| 351 | TRICHLORMETHIAZIDE | 79.0 | 92.7 | | 77.1 |
| 352 | TRYPTOPHAN (L) | 83.4 | 95.8 | | 77.2 |
| 353 | PYRANTEL PAMOATE | 86.2 | 95.3 | | 76.6 |
| 354 | RESORCINOL | 86.7 | 94.4 | | 73.7 |

| | | | | | |
|---|---|---|---|---|---|
| 355 | SPIRONOLACTONE | 83.3 | 94.0 | | 81.3 |
| 356 | SULFAPYRIDINE | 84.2 | 92.6 | | 78.1 |
| 357 | TETRAHYDROZOLINE HYDROCHLORIDE | 85.2 | 96.9 | | 80.8 |
| 358 | TOLBUTAMIDE | 84.2 | 90.2 | | 81.0 |
| 359 | TRIFLUOPERAZINE HYDROCHLORIDE | 299.1 | 348.7 | | 98.3 |
| 360 | TUAMINOHEPTANE SULFATE | 84.2 | 94.8 | | 81.7 |
| 361 | PYRAZINAMIDE | 84.6 | 98.2 | | 85.6 |
| 362 | RIFAMPIN | 84.2 | 99.5 | | 82.7 |
| 363 | STREPTOMYCIN SULFATE [5mM; 10% aq | 81.9 | 94.0 | | 72.7 |
| 364 | SULFASALAZINE | 82.7 | 95.1 | | 75.8 |
| 365 | THEOPHYLLINE | 80.2 | 96.8 | | 75.6 |
| 366 | TOLMETIN SODIUM | 85.8 | 98.1 | | 83.9 |
| 367 | TRIHEXYPHENIDYL HYDROCHLORIDE | 87.7 | 96.9 | | 74.1 |
| 368 | TUBOCURARINE CHLORIDE PENTAHYDRA | 79.5 | 94.0 | | 72.2 |
| 369 | PYRILAMINE MALEATE | 86.6 | 98.1 | | 85.8 |
| 370 | ROXARSONE | 83.0 | 92.0 | | 83.5 |
| 371 | STREPTOZOSIN | 82.5 | 93.7 | | 83.1 |
| 372 | SULFATHIAZOLE | 84.7 | 98.1 | | 81.3 |
| 373 | THIABENDAZOLE | 83.3 | 96.4 | | 83.2 |
| 374 | TOLNAFTATE | 81.8 | 89.2 | | 81.4 |
| 375 | TRIMEPRAZINE TARTRATE | 88.5 | 108.8 | | 83.0 |
| 376 | UREA | 85.5 | 95.2 | | 79.2 |
| 377 | PYRIMETHAMINE | 85.7 | 103.1 | | 76.3 |
| 378 | SALICYL ALCOHOL | 83.5 | 96.5 | | 74.6 |
| 379 | SULFABENZAMIDE | 81.2 | 97.2 | | 75.2 |
| 380 | SULFINPYRAZONE | 83.8 | 93.7 | | 74.7 |
| 381 | THIMEROSAL | | | | |
| 382 | TRANYLCYPROMINE SULFATE | 76.4 | 95.2 | | 75.8 |
| 383 | TRIMETHOBENZAMIDE HYDROCHLORIDE | 80.6 | 89.7 | | 77.3 |
| 384 | URSODIOL | 83.5 | 88.6 | | 77.5 |
| 385 | PYRVINIUM PAMOATE | 79.4 | 105.5 | | 92.5 |
| 386 | SALICYLAMIDE | 83.4 | 96.9 | | 73.7 |
| 387 | SULFACETAMIDE | 79.1 | 93.7 | | 85.6 |
| 388 | SULFISOXAZOLE | 76.8 | 94.6 | | 82.9 |
| 389 | THIOGUANINE | 76.6 | 94.3 | | 84.3 |
| 390 | TRIACETIN | 82.5 | 92.2 | | 84.3 |
| 391 | TRIMETHOPRIM | 70.3 | 88.0 | | 83.2 |
| 392 | VALPROATE SODIUM | 83.2 | 85.4 | | 78.6 |
| 393 | QUINACRINE HYDROCHLORIDE | 84.7 | 97.8 | | 88.8 |
| 394 | SODIUM SALICYLATE | 80.2 | 94.1 | | 72.7 |
| 395 | SULFADIAZINE | 85.4 | 96.2 | | 82.1 |
| 396 | SULINDAC | 80.4 | 97.9 | | 78.0 |
| 397 | THIORIDAZINE HYDROCHLORIDE | 110.0 | 170.0 | | 89.4 |
| 398 | TRIAMCINOLONE | 81.3 | 91.7 | | 89.0 |
| 399 | TRIOXSALEN | 80.1 | 93.6 | | 88.7 |
| 400 | VANCOMYCIN HYDROCHLORIDE | 82.3 | 88.9 | | 76.2 |
| 401 | VIDARABINE | 75.3 | 87.0 | | 77.1 |
| 402 | STRYCHNINE SULFATE | 76.6 | 86.5 | | 80.8 |
| 403 | ALBUTEROL | 62.8 | 86.0 | | 74.1 |
| 404 | CEFACLOR | 77.8 | 79.6 | | 78.5 |
| 405 | CANRENOIC ACID, POTASSIUM SALT | 74.4 | 89.6 | | 77.1 |
| 406 | FLUNARIZINE HYDROCHLORIDE | 78.7 | 88.9 | | 78.2 |
| 407 | FLUFENAMIC ACID | 74.1 | 84.5 | | 77.8 |
| 408 | MEBEVERINE HYDROCHLORIDE | 76.6 | 93.7 | | 72.7 |
| 409 | VINBLASTINE SULFATE | 78.9 | 87.1 | | 81.3 |
| 410 | DICUMAROL | 72.3 | 83.2 | | 80.7 |
| 411 | ARECOLINE HYDROBROMIDE | 64.1 | 83.5 | | 77.2 |
| 412 | IODIPAMIDE | 68.2 | 85.9 | | 75.6 |
| 413 | CHENODIOL | 77.5 | 84.5 | | 75.0 |
| 414 | FLUPHENAZINE HYDROCHLORIDE | 141.8 | 184.9 | | 76.2 |
| 415 | FENBENDAZOLE | 78.5 | 86.0 | | 81.4 |
| 416 | MECLOCYCLINE SULFOSALICYLATE | 80.7 | 87.2 | | 78.2 |
| 417 | WARFARIN | 69.5 | 79.5 | | 81.8 |
| 418 | YOHIMBINE HYDROCHLORIDE | 60.6 | 85.6 | | 79.4 |
| 419 | CAPTOPRIL | 72.6 | 89.5 | | 77.3 |
| 420 | LEVOTHYROXINE SODIUM | 65.7 | 84.7 | | 78.9 |
| 421 | CHOLECALCIFEROL | 85.1 | 101.5 | | 76.4 |
| 422 | FLUTAMIDE | 73.7 | 85.7 | | 79.1 |
| 423 | FENSPIRIDE HYDROCHLORIDE | 68.7 | 85.0 | | 77.6 |
| 424 | PROGLUMIDE | 61.4 | 88.0 | | 76.2 |
| 425 | XYLOMETAZOLINE HYDROCHLORIDE | 79.4 | 89.9 | | 77.5 |
| 426 | ACEBUTOLOL HYDROCHLORIDE | 77.2 | 86.2 | | 76.9 |

| | | | | | |
|---|---|---|---|---|---|
| 427 | CIMETIDINE | 72.0 | 86.1 | | 77.1 |
| 428 | LIOTHYRONINE | 73.6 | 84.2 | | 78.5 |
| 429 | CYANOCOBALAMIN | 82.4 | 99.9 | | 90.1 |
| 430 | DROPERIDOL | 71.2 | 86.2 | | 75.6 |
| 431 | MEFENAMIC ACID | 69.7 | 83.4 | | 76.6 |
| 432 | MINAPRINE HYDROCHLORIDE | 70.9 | 89.2 | | 75.4 |
| 433 | ZOMEPIRAC SODIUM | 75.8 | 91.1 | | 81.0 |
| 434 | ADENOSINE 5-MONOPHOSPHATE | 74.1 | 88.2 | | 78.8 |
| 435 | NEBIVOLOL HYDROCHLORIDE | 85.2 | 117.3 | | 79.8 |
| 436 | ALLANTOIN | 71.5 | 83.5 | | 79.3 |
| 437 | CHOLESTEROL | 71.9 | 84.3 | | 79.3 |
| 438 | FAMOTIDINE | 72.6 | 85.5 | | 73.8 |
| 439 | METHACYCLINE HYDROCHLORIDE | 74.7 | 87.7 | | 80.0 |
| 440 | MEMANTINE HYDROCHLORIDE | 74.7 | 81.9 | | 77.5 |
| 441 | MERBROMIN | 106.7 | 124.5 | | 113.8 |
| 442 | KETOTIFEN FUMARATE | 72.9 | 86.6 | | 78.9 |
| 443 | HYDRASTINE (1R, 9S) | 70.9 | 80.8 | | 76.8 |
| 444 | ALTHIAZIDE | 73.7 | 85.2 | | 76.8 |
| 445 | PIPERINE | 70.5 | 82.7 | | 80.9 |
| 446 | ETODOLAC | 74.7 | 87.3 | | 78.9 |
| 447 | PUROMYCIN DIHYDROCHLORIDE | 72.6 | 81.9 | | 79.4 |
| 448 | ACECLIDINE | 72.0 | 86.9 | | 78.1 |
| 449 | PHENACETIN | 75.5 | 88.3 | | 78.3 |
| 450 | BETAHISTINE HYDROCHLORIDE | 75.1 | 78.0 | | 79.9 |
| 451 | LIDOCAINE HYDROCHLORIDE | 74.7 | 88.1 | | 77.4 |
| 452 | ADENINE | 74.7 | 87.5 | | 78.6 |
| 453 | ETOPOSIDE | 73.3 | 82.2 | | 73.8 |
| 454 | FENOTEROL HYDROBROMIDE | 74.3 | 84.8 | | 79.1 |
| 455 | MEFEXAMIDE HYDROCHLORIDE | 72.6 | 84.0 | | 73.3 |
| 456 | TRAMIPROSATE | 74.3 | 83.7 | | 77.0 |
| 457 | CHLOROQUINE DIPHOSPHATE | 75.6 | 83.0 | | 78.9 |
| 458 | ETICLOPRIDE HYDROCHLORIDE | 80.6 | 94.3 | | 76.3 |
| 459 | PHENTOLAMINE HYDROCHLORIDE | 76.3 | 87.6 | | 75.1 |
| 460 | AMINACRINE | 82.2 | 99.2 | | 88.9 |
| 461 | DEHYDROCHOLIC ACID | 76.2 | 86.6 | | 75.7 |
| 462 | FENBUFEN | 71.6 | 84.5 | | 78.1 |
| 463 | PROBUCOL | 72.3 | 79.8 | | 75.3 |
| 464 | ATENOLOL | 75.7 | 84.4 | | 73.7 |
| 465 | PHENYLMERCURIC ACETATE | | | | |
| 466 | MYCOPHENOLIC ACID | 68.9 | 83.0 | | 79.8 |
| 467 | NALIDIXIC ACID | 76.0 | 88.0 | | 79.9 |
| 468 | BEKANAMYCIN SULFATE | 74.9 | 84.4 | | 78.3 |
| 469 | AZATADINE MALEATE | 76.8 | 87.1 | | 76.5 |
| 470 | FENOFIBRATE | 75.6 | 84.8 | | 79.6 |
| 471 | MEBENDAZOLE | 77.2 | 85.6 | | 79.7 |
| 472 | CAPSAICIN | 74.8 | 88.6 | | 79.7 |
| 473 | AZELAIC ACID | 62.7 | 84.7 | | 74.6 |
| 474 | INDOPROFEN | 70.4 | 82.7 | | 77.1 |
| 475 | BUTAMBEN | 58.9 | 86.9 | | 77.3 |
| 476 | BUDESONIDE | 73.2 | 86.5 | | 78.6 |
| 477 | FLUMEQUINE | 75.0 | 89.0 | | 78.0 |
| 478 | FENOPROFEN | 73.0 | 86.4 | | 89.2 |
| 479 | PROPOXYCAINE HYDROCHLORIDE | 75.3 | 85.1 | | 57.6 |
| 480 | CARBETAPENTANE CITRATE | 76.9 | 87.4 | | 81.1 |
| 481 | ROSIGLITAZONE MALEATE | 78.4 | 80.7 | | 81.4 |
| 482 | SULFAQUINOXALINE SODIUM | 72.3 | 81.7 | | 80.7 |
| 483 | SULFAMETHOXYPYRIDAZINE | 72.6 | 87.6 | | 81.5 |
| 484 | GALANTAMINE | 72.3 | 82.0 | | 81.9 |
| 485 | ISOTRETINON | 73.8 | 83.8 | | 81.4 |
| 486 | CEFOXITIN SODIUM | 71.2 | 84.7 | | 79.3 |
| 487 | DAPAGLIFLOZIN | 67.9 | 82.5 | | 86.8 |
| 488 | CARBIDOPA | 76.8 | 83.5 | | 90.4 |
| 489 | NICERGOLINE | 76.1 | 89.2 | | 79.4 |
| 490 | SULFAMONOMETHOXINE | 71.2 | 84.4 | | 82.5 |
| 491 | SACCHARIN | 69.8 | 88.0 | | 79.4 |
| 492 | RETINOL | 72.6 | 82.5 | | 83.4 |
| 493 | MESNA | 71.9 | 84.7 | | 80.4 |
| 494 | SURAMIN HEXASODIUM | 61.6 | 81.5 | | 86.9 |
| 495 | ALRESTATIN | 71.6 | 85.4 | | 84.2 |
| 496 | PIRACETAM | 73.4 | 86.9 | | 83.5 |
| 497 | PIMOZIDE | 80.2 | 99.9 | | 80.3 |
| 498 | SULCONAZOLE NITRATE | | | | |

| | | | | | |
|---|---|---|---|---|---|
| 499 | OCTODRINE | 73.6 | 84.4 | | 85.8 |
| 500 | LANATOSIDE C | 73.1 | 84.4 | | 84.9 |
| 501 | TRETINOIN | 74.6 | 83.0 | | 88.3 |
| 502 | CEFUROXIME SODIUM | 71.9 | 80.1 | | 84.5 |
| 503 | PROADIFEN HYDROCHLORIDE | 81.1 | 100.8 | | 81.9 |
| 504 | ETHOSUXIMIDE | 72.3 | 86.5 | | 80.8 |
| 505 | NICARDIPINE HYDROCHLORIDE | 78.6 | 84.8 | | 80.7 |
| 506 | RITODRINE HYDROCHLORIDE | 71.9 | 84.2 | | 79.5 |
| 507 | ERYTHROMYCIN ESTOLATE | 106.9 | 143.8 | | 83.1 |
| 508 | ENALAPRIL MALEATE | 71.9 | 77.1 | | 79.6 |
| 509 | BRETYLIUM TOSYLATE | 68.6 | 81.5 | | 80.2 |
| 510 | SUVOREXANT | 68.1 | 78.8 | | 85.2 |
| 511 | BRINZOLAMIDE | 72.6 | 83.0 | | 80.4 |
| 512 | PIPERIDOLATE HYDROCHLORIDE | 72.6 | 86.7 | | 82.3 |
| 513 | NEFOPAM | 74.5 | 83.3 | | 76.2 |
| 514 | SULPIRIDE (S [-]) | 72.3 | 82.5 | | 85.0 |
| 515 | ESTRADIOL BENZOATE | 70.4 | 86.2 | | 87.7 |
| 516 | KETOPROFEN | 68.2 | 82.3 | | 84.0 |
| 517 | FOSCARNET SODIUM | 70.2 | 84.8 | | 83.4 |
| 518 | CEFAMANDOLE SODIUM | 73.3 | 85.4 | | 83.4 |
| 519 | CISPLATIN | 54.7 | 78.1 | | 78.5 |
| 520 | ANISINDIONE | 71.9 | 83.2 | | 82.5 |
| 521 | PIRENZEPINE HYDROCHLORIDE | 76.6 | 81.7 | | 78.0 |
| 522 | RANITIDINE HYDROCHLORIDE | 75.0 | 83.5 | | 82.4 |
| 523 | ECONAZOLE NITRATE | | | | |
| 524 | LISINOPRIL | 60.5 | 81.9 | | 81.4 |
| 525 | OXYCLOZANIDE | 73.0 | 79.1 | | 86.1 |
| 526 | FOSFOMYCIN CALCIUM | 72.6 | 81.2 | | 81.5 |
| 527 | ZIDOVUDINE [AZT] | 74.1 | 83.0 | | 80.6 |
| 528 | CYCLOSPORINE | 83.4 | 78.0 | | 83.8 |
| 529 | PRAMOXINE HYDROCHLORIDE | 101.3 | 124.5 | | 94.5 |
| 530 | SPIPERONE | 73.0 | 84.1 | | 82.7 |
| 531 | FLUNISOLIDE | 69.9 | 75.5 | | 80.7 |
| 532 | BUMETANIDE | 71.6 | 84.7 | | 81.1 |
| 533 | PHTHALYLSULFATHIAZOLE | 74.4 | 81.5 | | 81.8 |
| 534 | CEFMETAZOLE SODIUM | 73.0 | 83.2 | | 81.1 |
| 535 | AZACITIDINE | 72.3 | 85.4 | | 82.5 |
| 536 | FLEROXACIN | 71.9 | 83.0 | | 79.3 |
| 537 | MEPHENESIN | 98.1 | 86.2 | | 83.2 |
| 538 | SULOCTIDIL | | | | |
| 539 | FLUMETHASONE | 76.7 | 81.7 | | 82.3 |
| 540 | CARBENOXOLONE SODIUM | 74.9 | 83.2 | | 80.0 |
| 541 | SUCCINYLSULFATHIAZOLE | 73.9 | 77.5 | | 81.6 |
| 542 | CEFAMANDOLE NAFATE | 74.6 | 77.6 | | 79.8 |
| 543 | CYCLOHEXIMIDE | | | | |
| 544 | ASCORBIC ACID | 71.9 | 82.5 | | 81.0 |
| 545 | SULFACHLORPYRIDAZINE | 74.3 | 81.8 | | 85.0 |
| 546 | RONIDAZOLE | 69.8 | 82.5 | | 83.1 |
| 547 | XYLAZINE | 68.5 | 80.4 | | 78.3 |
| 548 | CARPROFEN | 71.2 | 82.5 | | 77.8 |
| 549 | CEPHALEXIN | 74.4 | 83.7 | | 82.4 |
| 550 | CEFOPERAZONE | 71.5 | 83.4 | | 75.4 |
| 551 | AZASERINE | 71.3 | 74.7 | | 77.5 |
| 552 | MENADIONE | 76.5 | 87.5 | | 76.1 |
| 553 | SULFADIMETHOXINE | 84.9 | 84.4 | | 79.3 |
| 554 | SULFAMETER | 72.6 | 87.0 | | 81.9 |
| 555 | TOLAZAMIDE | 77.4 | 84.2 | | 77.6 |
| 556 | AVANAFIL | 68.4 | 86.2 | | 79.9 |
| 557 | ACARBOSE | 73.9 | 88.2 | | 81.8 |
| 558 | BEZAFIBRATE | 75.0 | 85.6 | | 70.9 |
| 559 | TINIDAZOLE | 76.6 | 84.2 | | 62.1 |
| 560 | SALICIN | 73.6 | 85.2 | | 80.7 |
| 561 | MONENSIN SODIUM | 89.3 | 88.2 | | 83.2 |
| 562 | DIAVERIDINE | 85.7 | 89.5 | | 84.5 |
| 563 | TENYLIDONE | 85.0 | 83.5 | | 81.4 |
| 564 | LOMEFLOXACIN HYDROCHLORIDE | 84.4 | 88.4 | | 79.4 |
| 565 | DIFLOXACIN HYDROCHLORIDE | 84.1 | 87.1 | | 80.7 |
| 566 | AMLEXANOX | 85.4 | 88.7 | | 80.3 |
| 567 | DOXIFLURIDINE | 86.1 | 91.2 | | 80.0 |
| 568 | CEFMENOXIME HYDROCHLORIDE | 85.8 | 90.1 | | 85.0 |
| 569 | ISOCONAZOLE NITRATE | 459.5 | | | 582.2 |
| 570 | LOMUSTINE | 85.0 | 85.2 | | 75.3 |

| | | | | | |
|---|---|---|---|---|---|
| 571 | QUINESTROL | 88.2 | 93.7 | | 81.2 |
| 572 | URACIL | 83.9 | 88.7 | | 76.9 |
| 573 | ZALCITABINE | 81.8 | 87.7 | | 80.6 |
| 574 | AMPROLIUM | 84.7 | 90.1 | | 80.8 |
| 575 | JOSAMYCIN | 79.3 | 89.0 | | 85.5 |
| 576 | CEFORANIDE | 83.7 | 89.6 | | 80.8 |
| 577 | LEVETIRACETAM | 84.4 | 89.6 | | 83.7 |
| 578 | LORNOXICAM | 83.3 | 89.7 | | 81.4 |
| 579 | RACTOPAMINE HYDROCHLORIDE | 81.6 | 87.8 | | 82.3 |
| 580 | VORICONAZOLE | 204.8 | 225.7 | | 140.1 |
| 581 | RITONAVIR | 82.5 | 87.8 | | 79.4 |
| 582 | MARBOFLOXACIN | 84.1 | 83.8 | | 82.0 |
| 583 | DIHYDROSTREPTOMYCIN SESQUISULFATI | 78.4 | 85.5 | | 80.1 |
| 584 | CEFOTETAN | 83.4 | 87.2 | | 81.2 |
| 585 | alpha-TOCHOPHEROL [4 mM] | 83.7 | 84.3 | | 82.3 |
| 586 | IOHEXOL | 83.0 | 85.5 | | 84.0 |
| 587 | RASAGILINE | 80.5 | 91.0 | | 74.4 |
| 588 | DICLAZURIL | 82.0 | 84.0 | | 78.5 |
| 589 | GANCICLOVIR HYDRATE | 83.7 | 88.0 | | 82.5 |
| 590 | NITRENDIPINE | 81.1 | 87.1 | | 80.2 |
| 591 | ANTAZOLINE | 81.1 | 85.1 | | 84.0 |
| 592 | CLINAFOXACIN HYDROCHLORIDE | 83.9 | 91.5 | | 82.0 |
| 593 | CAPECITABINE | 81.6 | 90.1 | | 75.6 |
| 594 | MEROPENEM | 83.9 | 89.6 | | 80.9 |
| 595 | ROXATIDINE ACETATE HYDROCHLORIDE | 82.3 | 84.5 | | 79.7 |
| 596 | IDOXURIDINE | 81.1 | 86.7 | | 77.2 |
| 597 | DEXIBUPROFEN | 81.8 | 86.2 | | 82.0 |
| 598 | EPRINOMECTIN | 84.0 | 92.6 | | 81.5 |
| 599 | OFLOXACIN | 81.6 | 85.2 | | 77.6 |
| 600 | ONDANSETRON HYDROCHLORIDE | 83.7 | 86.1 | | 82.2 |
| 601 | alpha-TOCHOPHERYL ACETATE [4mM] | 78.6 | 86.7 | | 81.8 |
| 602 | NEFAZODONE HYDROCHLORIDE | 84.8 | 96.9 | | 76.1 |
| 603 | SERATRODAST | 81.8 | 80.1 | | 81.6 |
| 604 | ABACAVIR SULFATE | 81.4 | 85.0 | | 80.4 |
| 605 | PIZOTYLINE MALATE | 82.3 | 92.6 | | 77.6 |
| 606 | FENCLONINE (+/-) | 81.4 | 83.8 | | 77.7 |
| 607 | NITARSONE | 79.0 | 85.2 | | 76.5 |
| 608 | CINROMIDE | 81.5 | 89.3 | | 79.8 |
| 609 | SISOMICIN SULFATE | 89.7 | 102.1 | | 69.8 |
| 610 | NIFURSOL | 81.2 | 83.9 | | 77.4 |
| 611 | SPARFLOXACIN | 83.0 | 88.9 | | 83.0 |
| 612 | LEVONORGESTREL | 81.4 | 86.9 | | 70.3 |
| 613 | AMOROLFINE HYDROCHLORIDE | 135.9 | 213.4 | | 168.5 |
| 614 | LINEZOLID | 82.5 | 85.2 | | 77.8 |
| 615 | ACENOCOUMAROL | 79.3 | 85.4 | | 81.3 |
| 616 | CINTRIAMIDE | 82.3 | 86.1 | | 77.3 |
| 617 | ANASTROZOLE | 82.3 | 87.4 | | 76.9 |
| 618 | PALIPERIDONE | 80.7 | 87.7 | | 78.0 |
| 619 | STAVUDINE | 79.1 | 84.6 | | 83.0 |
| 620 | LEVOSIMENDAN | 80.1 | 87.4 | | 80.5 |
| 621 | NEPAFENAC | 80.0 | 87.1 | | 75.9 |
| 622 | LUMIRACOXIB | 79.9 | 84.0 | | 79.1 |
| 623 | BALSALAZIDE DISODIUM | 79.4 | 93.7 | | 79.7 |
| 624 | BENZOIC ACID | 81.0 | 85.0 | | 77.5 |
| 625 | THIAMINE HYDROCHLORIDE | 82.5 | 86.9 | | 86.1 |
| 626 | PENCICLOVIR | 82.1 | 85.9 | | 77.6 |
| 627 | SULBACTAM | 79.3 | 87.1 | | 74.8 |
| 628 | LOFEXIDINE HYDROCHLORIDE | 80.0 | 84.0 |  | 75.4 |
| 629 | CANDESARTAN | 79.3 | 88.0 | | 77.3 |
| 630 | DIPERODON HYDROCHLORIDE | 78.3 | 88.9 | | 79.3 |
| 631 | VALDECOXIB | 79.3 | 84.7 | | 78.0 |
| 632 | BENZYL BENZOATE | 80.6 | 84.0 | | 81.6 |
| 633 | NALOXONE HYDROCHLORIDE | 80.1 | 87.1 | | 78.7 |
| 634 | MIZORIBINE | 79.4 | 87.2 | | 82.5 |
| 635 | TAZOBACTAM | 79.4 | 90.4 | | 77.5 |
| 636 | DOXOFYLLINE | 79.0 | 86.5 | | 78.2 |
| 637 | ADAPALENE | 81.8 | 87.2 | | 81.2 |
| 638 | OLSALAZINE SODIUM | 81.2 | 84.8 | | 77.3 |
| 639 | CEFEPIME HYDROCHLORIDE | 80.6 | 87.8 | | 80.9 |
| 640 | BENZOYL PEROXIDE | 82.5 | 88.3 | | 78.3 |
| 641 | BETAINE HYDROCHLORIDE | 82.5 | 88.7 | | 84.4 |
| 642 | QUINAPRIL HYDROCHLORIDE | 81.0 | 88.5 | | 83.7 |

| | | | | | |
|---|---|---|---|---|---|
| 643 | BROMHEXINE HYDROCHLORIDE | 88.9 | 96.0 | | 86.3 |
| 644 | CHLOROXINE | 419.3 | 257.5 | | |
| 645 | PERHEXILINE MALEATE | 414.3 | 422.7 | | 98.2 |
| 646 | METARAMINOL BITARTRATE | 84.0 | 86.4 | | 85.0 |
| 647 | PAROMOMYCIN SULFATE | 83.0 | 87.6 | | 85.9 |
| 648 | THIRAM | 161.2 | 343.0 | | 94.3 |
| 649 | BIOTIN | 80.6 | 84.7 | | 85.9 |
| 650 | METHYSERGIDE MALEATE | 80.4 | 86.7 | | 86.0 |
| 651 | TYROSINE HYDROCHLORIDE | 82.7 | 81.5 | | 86.3 |
| 652 | CHLORPROTHIXENE HYDROCHLORIDE | 109.6 | 130.4 | | 83.8 |
| 653 | METHAPYRILENE HYDROCHLORIDE | 79.5 | 84.3 | | 80.1 |
| 654 | METHAZOLAMIDE | 82.7 | 87.7 | | 82.7 |
| 655 | TRANILAST | 81.6 | 86.2 | | 80.8 |
| 656 | THIOTEPA | 83.2 | 83.9 | | 81.4 |
| 657 | AKLOMIDE | 81.5 | 90.6 | | 88.0 |
| 658 | AMIFOSTINE | 83.0 | 84.6 | | 83.1 |
| 659 | CEFTRIAXONE SODIUM TRIHYDRATE | 81.1 | 85.5 | | 89.1 |
| 660 | CINNARAZINE | 93.2 | 95.6 | | 88.7 |
| 661 | MEPRYLCAINE HYDROCHLORIDE | 84.0 | 87.0 | | 82.9 |
| 662 | METHYLBENZETHONIUM CHLORIDE | | | | 163.7 |
| 663 | PRILOCAINE HYDROCHLORIDE | 81.7 | 84.4 | | 82.9 |
| 664 | TETROQUINONE | 84.8 | 86.2 | | 83.2 |
| 665 | MONOBENZONE | 84.9 | 85.5 | | 85.4 |
| 666 | INAMRINONE | 79.6 | 85.2 | | 83.8 |
| 667 | VINPOCETINE | 85.2 | 84.0 | | 83.7 |
| 668 | CYCLOBENZAPRINE HYDROCHLORIDE | 84.6 | 93.8 | | 84.2 |
| 669 | HALCINONIDE | 80.8 | 85.7 | | 74.8 |
| 670 | METHYLPREDNISOLONE SODIUM SUCCIN | 82.5 | 84.8 | | 80.5 |
| 671 | HYDROCORTISONE BUTYRATE | 83.5 | 87.2 | | 80.7 |
| 672 | SULFANITRAN | 84.6 | 84.4 | | 82.1 |
| 673 | NICOTINYL ALCOHOL TARTRATE | 83.7 | 87.7 | | 88.4 |
| 674 | TIAPRIDE HYDROCHLORIDE | 82.0 | 87.8 | | 76.8 |
| 675 | TRIMIPRAMINE MALEATE | 85.5 | 90.3 | | 83.7 |
| 676 | DANTROLENE SODIUM | 82.7 | 91.5 | | 83.6 |
| 677 | HYCANTHONE | 88.5 | 97.0 | | 86.9 |
| 678 | AMSACRINE | 86.4 | 88.7 | | 89.1 |
| 679 | ROXITHROMYCIN | 83.4 | 83.1 | | 85.3 |
| 680 | OXIBENDAZOLE | 82.7 | 85.0 | | 85.7 |
| 681 | FLOXURIDINE | 84.1 | 82.5 | | 85.6 |
| 682 | GLUCONOLACTONE | 82.5 | 82.7 | | 85.5 |
| 683 | RITANSERIN | 85.5 | 88.2 | | 84.5 |
| 684 | BETAMETHASONE 17,21-DIPROPIONATE | 81.6 | 85.2 | | 85.9 |
| 685 | PYRIDOSTIGMINE BROMIDE | 83.2 | 89.1 | | 78.1 |
| 686 | MIDODRINE HYDROCHLORIDE | 83.4 | 84.4 | | 83.3 |
| 687 | MITOXANTRONE HYDROCHLORIDE | 87.9 | 98.0 | | 86.4 |
| 688 | PIPOBROMAN | 84.4 | 85.5 | | 84.5 |
| 689 | ALTRETAMINE | 86.1 | 83.6 | | 87.4 |
| 690 | AZLOCILLIN SODIUM | 85.2 | 88.8 | | 87.6 |
| 691 | TRAZODONE HYDROCHLORIDE | 84.9 | 86.6 | | 86.5 |
| 692 | IMIQUIMOD HYDROCHLORIDE | 85.7 | 88.8 | | 85.1 |
| 693 | ISOXICAM | 84.0 | 87.1 | | 85.7 |
| 694 | BERGENIN | 84.4 | 83.0 | | 87.9 |
| 695 | OXETHAZAINE | 79.9 | 84.4 | | 88.2 |
| 696 | NAFRONYL OXALATE | 86.8 | 94.3 | | 89.9 |
| 697 | AMINOHIPPURIC ACID | 82.2 | 92.7 | | 86.5 |
| 698 | BACAMPICILLIN HYDROCHLORIDE | 83.5 | 86.1 | | 86.0 |
| 699 | THONZYLAMINE HYDROCHLORIDE | 84.5 | 87.1 | | 89.2 |
| 700 | EDOXUDINE | 82.0 | 85.4 | | 82.0 |
| 701 | LABETALOL HYDROCHLORIDE | 85.6 | 81.5 | | 88.0 |
| 702 | NALTREXONE HYDROCHLORIDE | 83.2 | 86.9 | | 81.0 |
| 703 | DIPYRONE | 81.8 | 86.1 | | 81.9 |
| 704 | QUIPAZINE MALEATE | 89.6 | 89.0 | | 81.5 |
| 705 | MEFLOQUINE HYDROCHLORIDE | 119.1 | 147.0 | | 90.5 |
| 706 | BENDROFLUMETHIAZIDE | 80.3 | 85.1 | | 74.3 |
| 707 | THIAMPHENICOL | 81.4 | 88.1 | | 85.0 |
| 708 | ENOXACIN | 82.0 | 88.2 | | 87.7 |
| 709 | LASALOCID SODIUM | 87.1 | 89.1 | | 85.1 |
| 710 | CYCLOTHIAZIDE | 83.0 | 88.3 | | 83.4 |
| 711 | SULFANILATE ZINC | 80.7 | 82.0 | | 86.7 |
| 712 | FOLIC ACID | 83.5 | 86.5 | | 86.2 |
| 713 | ADIPHENINE HYDROCHLORIDE | 83.2 | 89.3 | | 74.4 |
| 714 | OLMESARTAN | 82.7 | 86.5 | | 89.6 |

| | | | | | |
|---|---|---|---|---|---|
| 715 | TENOXICAM | 82.2 | 88.2 | | 80.4 |
| 716 | ETHISTERONE | 83.0 | 81.4 | | 78.9 |
| 717 | LEVAMISOLE HYDROCHLORIDE | 83.0 | 87.6 | | 74.2 |
| 718 | NICLOSAMIDE | 113.5 | 120.2 | | 98.0 |
| 719 | URETHANE | 82.7 | 79.3 | | 59.0 |
| 720 | HOMOSALATE | 83.7 | 86.7 | | 84.8 |
| 721 | SPIRAMYCIN | 91.0 | 90.8 | | 81.3 |
| 722 | BENZALKONIUM CHLORIDE HYDRATE | | | | |
| 723 | MEPHENYTOIN | 91.6 | 85.0 | | 82.6 |
| 724 | AMCINONIDE | 89.5 | 84.7 | | 82.3 |
| 725 | LAMIVUDINE | 86.8 | 83.2 | | 86.2 |
| 726 | PHYTONADIONE [5mM] | 91.3 | 86.1 | | 80.8 |
| 727 | SODIUM GLUCONATE | 87.8 | 85.9 | | 81.4 |
| 728 | LEVOCARNITINE | 88.2 | 86.0 | | 83.2 |
| 729 | ESZOPICLONE | 88.6 | 88.3 | | 81.7 |
| 730 | PENTOXIFYLLINE | 89.9 | 86.9 | | 77.6 |
| 731 | CLOPIDOGREL SULFATE | 85.7 | 84.1 | | 83.0 |
| 732 | BUPIVACAINE HYDROCHLORIDE | 86.1 | 86.1 | | 80.1 |
| 733 | BETAZOLE HYDROCHLORIDE | 90.6 | 86.6 | | 80.8 |
| 734 | PREGABALIN | 86.2 | 83.7 | | 83.7 |
| 735 | QUINETHAZONE | 92.3 | 86.4 | | 80.7 |
| 736 | IOPANIC ACID | 83.6 | 86.5 | | 81.6 |
| 737 | CHOLINE CHLORIDE | 89.5 | 88.7 | | 91.0 |
| 738 | CIPROFLOXACIN | 88.9 | 87.8 | | 83.2 |
| 739 | LORATADINE | 93.7 | 83.2 | | 83.5 |
| 740 | DROSPIRENONE | 84.9 | 85.4 | | 77.1 |
| 741 | DIXANTHOGEN | 111.2 | 102.1 | | 86.4 |
| 742 | CARBARSONE | 85.7 | 85.9 | | 82.7 |
| 743 | TILETAMINE HYDROCHLORIDE | 84.2 | 86.1 | | 83.0 |
| 744 | KETOROLAC TROMETHAMINE | 85.5 | 87.0 | | 81.4 |
| 745 | CLOFIBRATE | 91.5 | 84.9 | | 76.3 |
| 746 | BACLOFEN HYDROCHLORIDE (+/-) | 89.1 | 82.3 | | 79.6 |
| 747 | SELAMECTIN | 89.9 | 84.2 | | 81.6 |
| 748 | ECAMSULE TRIETHANOLAMINE | 88.6 | 80.9 | | 80.4 |
| 749 | NITAZOXANIDE | 90.7 | 89.2 | | 91.3 |
| 750 | TROLEANDOMYCIN | 88.1 | 81.6 | | 83.1 |
| 751 | DOCUSATE SODIUM | 219.9 | 178.9 | | 76.1 |
| 752 | LANSOPRAZOLE | 97.5 | 84.6 | | 80.7 |
| 753 | RESORCINOL MONOACETATE | 88.0 | 88.5 | | 84.2 |
| 754 | NABUMETONE | 93.7 | 87.2 | | 82.3 |
| 755 | ATORVASTATIN CALCIUM | 92.6 | 84.8 | | 83.4 |
| 756 | ENALAPRILAT | 88.6 | 82.5 | | 79.1 |
| 757 | COLESEVALAM HYDROCHLORIDE (high m | 88.0 | 83.5 | | 80.8 |
| 758 | VALGANCICLOVIR HYDROCHLORIDE | 84.2 | 84.4 | | 82.5 |
| 759 | RISEDRONATE SODIUM | 85.7 | 84.0 | | 80.1 |
| 760 | LEFLUNOMIDE | 86.8 | 86.4 | | 83.4 |
| 761 | NIMODIPINE | 86.9 | 85.5 | | 81.8 |
| 762 | CELECOXIB | 88.0 | 83.4 | | 82.4 |
| 763 | BENZYL ALCOHOL | 87.3 | 84.5 | | 82.2 |
| 764 | GADOTERIDOL | 85.7 | 85.2 | | 80.9 |
| 765 | TERCONAZOLE | 92.6 | 91.5 | | 80.3 |
| 766 | TAURINE | 88.0 | 83.4 | | 82.7 |
| 767 | DABIGATRAN ETEXILATE MESYLATE | 87.8 | 84.7 | | 80.7 |
| 768 | MEXILETINE HYDROCHLORIDE | 91.5 | 84.2 | | 72.3 |
| 769 | ACYCLOVIR | 87.6 | 89.5 | | 81.9 |
| 770 | ALENDRONATE SODIUM TRIHYDRATE | 88.7 | 86.9 | | 95.2 |
| 771 | ALLYLESTRENOL | 88.9 | 81.2 | | 76.4 |
| 772 | SEVOFLURANE | 87.4 | 83.5 | | 80.1 |
| 773 | VORINOSTAT | 85.7 | 82.5 | | 81.8 |
| 774 | SUCRALFATE [5mM] | 89.1 | 85.3 | | 80.7 |
| 775 | ALBENDAZOLE | 89.4 | 86.4 | | 83.5 |
| 776 | MORANTEL CITRATE | 90.1 | 85.6 | | 81.7 |
| 777 | PENFLURIDOL | 291.8 | 195.1 | | |
| 778 | BLEOMYCIN (bleomycin B2 shown) | 378.7 | 329.1 | | 343.0 |
| 779 | CEFTAZIDIME | 86.8 | 85.6 | | 80.6 |
| 780 | PHYSOSTIGMINE SALICYLATE | 87.6 | 81.5 | | 81.4 |
| 781 | PEMIROLAST POTASSIUM | 88.8 | 84.4 | | 81.0 |
| 782 | THONZONIUM BROMIDE | | | | |
| 783 | PACLITAXEL | 89.8 | 82.7 | | 79.5 |
| 784 | OXYPHENCYCLIMINE HYDROCHLORIDE | 91.1 | 86.4 | | 79.4 |
| 785 | THALIDOMIDE | 86.4 | 85.1 | | 82.3 |
| 786 | ANETHOLE | 80.1 | 77.0 | | 80.1 |

| | | | | | |
|---|---|---|---|---|---|
| 787 | COLFORSIN | 86.4 | 83.0 | | 79.6 |
| 788 | PHENFORMIN HYDROCHLORIDE | 90.2 | 85.2 | | 83.5 |
| 789 | ACETOPHENAZINE MALEATE | 92.1 | 90.9 | | 81.8 |
| 790 | PIPERACETAZINE | 87.0 | 80.8 | | 82.5 |
| 791 | BUTACAINE SULFATE | 88.6 | 81.5 | | 83.4 |
| 792 | PERPHENAZINE | 288.9 | 296.2 | | 99.5 |
| 793 | IOXILAN | 84.5 | 81.6 | | 85.5 |
| 794 | TERFENADINE | 133.5 | 125.3 | | 81.5 |
| 795 | ISOSORBIDE MONONITRATE | 74.5 | 81.3 | | 78.0 |
| 796 | TRIFLUPROMAZINE HYDROCHLORIDE | | | | |
| 797 | TOPOTECAN HYDROCHLORIDE | 86.6 | 83.7 | | 86.0 |
| 798 | PIROCTONE OLAMINE | 87.1 | 81.7 | | 80.8 |
| 799 | CLOBETASOL PROPIONATE | 86.7 | 81.3 | | 79.8 |
| 800 | PROPAFENONE HYDROCHLORIDE | 116.5 | 118.5 | | 81.1 |
| 801 | ALOGLIPTIN BENZOATE | 92.6 | 85.1 | | 77.1 |
| 802 | TANNIC ACID | 88.7 | 92.8 | | 85.3 |
| 803 | CITALOPRAM HYDROBROMIDE | 89.2 | 82.7 | | 79.4 |
| 804 | CHLOROGUANIDE HYDROCHLORIDE | 88.9 | 84.5 | | 78.3 |
| 805 | CARVEDILOL | 106.1 | 97.5 | | 75.2 |
| 806 | DEXLANSOPRAZOLE | 87.6 | 78.4 | | 77.3 |
| 807 | FENOFIBRIC ACID | 85.9 | 80.8 | | 75.5 |
| 808 | LETROZOLE | 86.1 | 80.2 | | 76.3 |
| 809 | TACROLIMUS | 87.7 | 82.0 | | 80.2 |
| 810 | TEPOXALIN | 113.4 | 88.7 | | 77.7 |
| 811 | FLUOXETINE HYDROCHLORIDE | 119.0 | 101.7 | | 76.2 |
| 812 | TRIMETOZINE | 87.8 | 83.2 | | 75.5 |
| 813 | NATEGLINIDE | 88.2 | 84.3 | | 75.4 |
| 814 | ARMODAFINIL | 84.6 | 82.3 | | 77.3 |
| 815 | CARVEDILOL PHOSPHATE | 101.5 | 90.7 | | 76.1 |
| 816 | CLOSANTEL | 77.8 | 80.7 | | 77.8 |
| 817 | FLUCONAZOLE | 99.0 | 93.2 | | 75.4 |
| 818 | AZILSARTAN MEDOXOMIL | 92.1 | 81.8 | | 73.5 |
| 819 | BUPROPION | 92.1 | 81.2 | | 76.2 |
| 820 | ACRISORCIN | 113.5 | 103.3 | | 87.0 |
| 821 | IRBESARTAN | 87.5 | 78.9 | | 75.0 |
| 822 | ROCURONIUM BROMIDE | 91.1 | 80.3 | | 77.5 |
| 823 | VILAZODONE HYDROCHLORIDE | 86.6 | 79.4 | | 77.1 |
| 824 | ARGININE HYDROCHLORIDE | 90.0 | 81.8 | | 74.4 |
| 825 | RAMELTEON | 86.9 | 84.9 | | 75.8 |
| 826 | BUTOCONAZOLE | 244.7 | | | 169.1 |
| 827 | CEFUROXIME AXETIL | 86.7 | 84.0 | | 73.4 |
| 828 | CYSTEAMINE HYDROCHLORIDE | 86.7 | 81.3 | | 73.7 |
| 829 | LEVOFLOXACIN | 90.5 | 81.1 | | 74.7 |
| 830 | ORLISTAT | 78.5 | 77.4 | | 69.3 |
| 831 | TRILOSTANE | 83.7 | 83.5 | | 75.1 |
| 832 | OSELTAMIVIR PHOSPHATE | 88.1 | 81.0 | | 76.4 |
| 833 | BROMPHENIRAMINE MALEATE | 79.8 | 82.7 | | 77.2 |
| 834 | ACETRIAZOIC ACID | 89.0 | 80.8 | | 76.3 |
| 835 | FEXOFENADINE HYDROCHLORIDE | 91.5 | 81.0 | | 77.0 |
| 836 | METAXALONE | 88.1 | 81.2 | | 76.5 |
| 837 | CANDESARTAN CILEXTIL | 102.9 | 89.1 | | 74.8 |
| 838 | MOXIFLOXACIN HYDROCHLORIDE | 87.1 | 83.0 | | 77.3 |
| 839 | MYCOPHENOLATE MOFETIL | 87.7 | 79.7 | | 76.6 |
| 840 | LEVOCETIRIZINE DIHYDROCHLORIDE | 88.0 | 81.5 | | 71.6 |
| 841 | SIROLIMUS | 509.4 | 274.9 | | 213.4 |
| 842 | DIRITHROMYCIN | 91.9 | 91.4 | | 74.5 |
| 843 | TRIFLURIDINE | 92.6 | 86.4 | | 76.3 |
| 844 | CLARITHROMYCIN | 91.0 | 80.6 | | 76.5 |
| 845 | OXIGLUTATIONE | 86.1 | 82.7 | | 76.1 |
| 846 | ENROFLOXACIN | 115.4 | 134.9 | | 75.5 |
| 847 | MILNACIPRAN HYDROCHLORIDE | 95.8 | 94.5 | | 75.9 |
| 848 | GLIMEPIRIDE | 87.0 | 79.4 | | 75.9 |
| 849 | SUPLATAST TOSYLATE | 86.2 | 85.3 | | 75.8 |
| 850 | MEPIVACAINE HYDROCHLORIDE | 84.6 | 84.2 | | 77.1 |
| 851 | AMINOLEVULINIC ACID HYDROCHLORIDE | 87.4 | 80.1 | | 76.3 |
| 852 | DOBUTAMINE HYDROCHLORIDE | 87.7 | 85.7 | | 74.8 |
| 853 | LOSARTAN | 85.5 | 82.0 | | 76.9 |
| 854 | TRIBROMOETHANOL | 85.4 | 82.7 | | 74.7 |
| 855 | IBANDRONATE SODIUM | 86.6 | 83.0 | | 74.4 |
| 856 | OCTINOXATE | 87.1 | 78.8 | | 77.2 |
| 857 | ETHYLNOREPINEPHRINE HYDROCHLORID | 88.2 | 88.3 | | 76.4 |
| 858 | MELOXICAM | 88.0 | 82.3 | | 76.6 |

| | | | | | |
|---|---|---|---|---|---|
| 859 | DAPTOMYCIN (5 millimolar/DMSO) | 86.6 | 81.3 | | 74.7 |
| 860 | SIMVASTATIN | 85.4 | 83.5 | | 74.8 |
| 861 | LITHIUM CITRATE HYDRATE | 85.7 | 74.7 | | 77.0 |
| 862 | PIOGLITAZONE HYDROCHLORIDE | 86.8 | 85.7 | | 67.4 |
| 863 | AZITHROMYCIN | 84.2 | 76.5 | | 76.0 |
| 864 | MANGAFODIPIR TRISODIUM | 87.4 | 82.5 | | 76.1 |
| 865 | TENIPOSIDE | 87.5 | 82.7 | | 74.4 |
| 866 | NILUTAMIDE | 88.2 | 77.6 | | 77.1 |
| 867 | AVOBENZONE | 85.8 | 79.4 | | 74.3 |
| 868 | HYDROQUINONE | 85.4 | 84.0 | | 79.2 |
| 869 | MIGLITOL | 85.2 | 79.9 | | 77.1 |
| 870 | DONEPEZIL HYDROCHLORIDE | 92.1 | 89.4 | | 77.2 |
| 871 | MEPHENTERMINE SULFATE | 85.0 | 81.3 | | 76.1 |
| 872 | FLUCYTOSINE | 172.4 | 160.6 | | 122.8 |
| 873 | CANDICIDIN | 221.6 | 374.2 | | 174.9 |
| 874 | VENLAFAXINE HYDROCHLORIDE | 86.6 | 83.4 | | 77.2 |
| 875 | ATOVAQUONE | 83.8 | 87.4 | | 79.6 |
| 876 | OXCARBAZEPINE | 85.0 | 81.6 | | 76.7 |
| 877 | DESVENLAFAXINE SUCCINATE | 82.7 | 83.3 | | 79.3 |
| 878 | FELODIPINE | 87.4 | 80.7 | | 79.4 |
| 879 | CEFTIOFUR HYDROCHLORIDE | 89.8 | 82.2 | | 79.4 |
| 880 | LEVALBUTEROL HYDROCHLORIDE | 86.6 | 81.2 | | 76.4 |
| 881 | PROPOFOL | 93.19 | 93.2 | | 72.3 |
| 882 | ESTROPIPATE | 88.33 | 86.6 | | 83.7 |
| 883 | AMLODIPINE BESYLATE | 140.38 | 147.1 | | 78.1 |
| 884 | PERINDOPRIL ERBUMINE | 86.23 | 84.9 | | 67.0 |
| 885 | TELMISARTAN | 86.57 | 82.5 | | 81.5 |
| 886 | OXFENDAZOLE | 86.06 | 85.3 | | 77.8 |
| 887 | LINAGLIPTIN | 85.70 | 84.2 | | 71.2 |
| 888 | MOMETASONE FUROATE | 81.41 | 86.2 | | 81.3 |
| 889 | MESORIDAZINE BESYLATE | 82.97 | 91.2 | | 80.8 |
| 890 | AZTREONAM | 82.99 | 82.5 | | 80.8 |
| 891 | EZETIMIBE | 81.04 | 84.7 | | 77.8 |
| 892 | ROSUVASTATIN CALCIUM | 79.89 | 89.8 | | 79.4 |
| 893 | SERTRALINE HYDROCHLORIDE | 209.97 | 180.5 | | 82.2 |
| 894 | AMITRAZ | 83.00 | 89.0 | | 73.9 |
| 895 | PAROXETINE HYDROCHLORIDE | 108.19 | 105.8 | | 77.1 |
| 896 | VALACYCLOVIR HYDROCHLORIDE | 85.36 | 84.9 | | 82.5 |
| 897 | ARTEMETHER | 133.90 | 151.9 | | 106.5 |
| 898 | CLAVULANATE LITHIUM | 85.04 | 85.8 | | 90.5 |
| 899 | ALMOTRIPTAN | 83.81 | 91.2 | | 95.5 |
| 900 | RAMIPRIL | 84.59 | 86.2 | | 93.2 |
| 901 | ALFUZOSIN HYDROCHLORIDE | 83.51 | 80.7 | | 91.5 |
| 902 | ASPARTAME | 85.04 | 83.3 | | 89.6 |
| 903 | AZELASTINE HYDROCHLORIDE | 116.93 | 117.1 | | 92.5 |
| 904 | ZOLPIDEM | 84.62 | 89.2 | | 95.8 |
| 905 | LOXAPINE SUCCINATE | 83.27 | 86.8 | | 81.0 |
| 906 | ETIDRONATE DISODIUM | 79.58 | 80.9 | | 73.4 |
| 907 | OLMESARTAN MEDOXOMIL | 81.25 | 84.5 | | 80.9 |
| 908 | TEGASEROD MALEATE | 88.41 | 88.5 | | 78.0 |
| 909 | TRANDOLAPRIL | 82.49 | 84.2 | | 76.7 |
| 910 | BIFONAZOLE | 291.00 | 65.3 | | 164.1 |
| 911 | KETANSERIN | 83.00 | 87.8 | | 81.1 |
| 912 | CETIRIZINE HYDROCHLORIDE | 85.55 | 84.2 | | 83.3 |
| 913 | FLUBENDAZOLE | 83.95 | 83.3 | | 80.1 |
| 914 | FLUVASTATIN SODIUM | 86.07 | 91.9 | | 86.1 |
| 915 | DICLORALUREA | 80.75 | 80.9 | | 77.7 |
| 916 | ESCITALOPRAM OXALATE | 78.41 | 82.7 | | 75.7 |
| 917 | TELITHROMYCIN | 82.74 | 87.5 | | 83.6 |
| 918 | TYLOSIN TARTRATE | 79.79 | 91.2 | | 74.7 |
| 919 | RIBOFLAVIN | 83.14 | 88.7 | | 82.7 |
| 920 | SUMATRIPTAN | 79.76 | 81.6 | | 83.0 |
| 921 | THIOSTREPTON | 80.84 | 84.4 | | 74.1 |
| 922 | PANTOPRAZOLE | 79.43 | 85.1 | | 78.3 |
| 923 | CEFTIBUTEN | 79.76 | 84.7 | | 79.6 |
| 924 | DERACOXIB | 79.94 | 84.1 | | 79.6 |
| 925 | OXAPROZIN | 76.83 | 80.1 | | 80.3 |
| 926 | SARAFLOXACIN HYDROCHLORIDE | 84.02 | 84.2 | | 73.7 |
| 927 | FIROCOXIB | 79.42 | 87.4 | | 80.2 |
| 928 | VARDENAFIL HYDROCHLORIDE | 80.65 | 84.5 | | 82.2 |
| 929 | TILMICOSIN | 87.22 | 86.6 | | 82.5 |
| 930 | QUETIAPINE FUMARATE | 83.00 | 82.7 | | 78.9 |

| | | | | | |
|---|---|---|---|---|---|
| 931 | CEFDINIR | 84.12 | 86.2 | | 79.6 |
| 932 | CILOSTAZOL | 86.09 | 84.6 | | 85.0 |
| 933 | PROPRANOLOL HYDROCHLORIDE (+/-) | 86.98 | 87.7 | | 73.3 |
| 934 | CLOPIDOL | 81.25 | 84.9 | | 72.8 |
| 935 | CEFDITORIN PIVOXIL | 83.27 | 84.4 | | 78.4 |
| 936 | ACEDAPSONE | 80.55 | 79.6 | | 83.2 |
| 937 | DESLORATADINE HYDROCHLORIDE | 91.45 | 104.6 | | 79.9 |
| 938 | BEMOTRIZINOL | 81.58 | 82.0 | | 74.6 |
| 939 | VALSARTAN | 81.53 | 88.3 | | 82.2 |
| 940 | CANRENONE | 79.76 | 87.5 | | 77.2 |
| 941 | CROTAMITON | 80.55 | 83.6 | | 77.0 |
| 942 | RIFAXIMIN | 86.50 | 88.4 | | 83.0 |
| 943 | MODAFINIL | 81.32 | 82.5 | | 83.7 |
| 944 | ATOMOXETINE HYDROCHLORIDE | 82.27 | 80.7 | | 87.2 |
| 945 | CLIOQUINOL | 117.39 | 113.2 | | 80.3 |
| 946 | DRONEDARONE HYDROCHLORIDE | 306.77 | 91.6 | | 79.4 |
| 947 | ISOFLUPREDONE ACETATE | 80.84 | 85.9 | | 74.7 |
| 948 | APRAMYCIN SULFATE | 80.02 | 78.6 | | 85.9 |
| 949 | NONOXYNOL-9 | 85.76 | 90.7 | | 86.5 |
| 950 | CHLORMADINONE ACETATE | 78.47 | 95.2 | | 85.8 |
| 951 | CEFPROZIL | 79.10 | 84.2 | | 73.4 |
| 952 | LORGLUMIDE SODIUM | 80.84 | 83.8 | | 74.4 |
| 953 | DIMPYLATE | 80.13 | 87.7 | | 77.6 |
| 954 | FAMCICLOVIR | 82.74 | 85.1 | | 78.2 |
| 955 | TORSEMIDE | 81.75 | 85.3 | | 76.7 |
| 956 | HYDROXYCHLOROQUINE SULFATE | 79.89 | 82.2 | | 80.5 |
| 957 | CARBADOX | 83.24 | 86.8 | | 81.6 |
| 958 | OXICONAZOLE NITRATE | 311.80 | 96.8 | | 205.4 |
| 959 | RANOLAZINE DIHYDROCHLORIDE | 80.07 | 88.1 | | 82.1 |
| 960 | LEVOBUNOLOL HYDROCHLORIDE | 77.86 | 88.3 | | 82.1 |
| 961 | TRAMADOL HYDROCHLORIDE | 250.54 | 46.8 | | 133.9 |
| 962 | BENZOXIQUINE | 83.84 | 80.5 | | 78.5 |
| 963 | CARAZOLOL | 80.13 | 85.3 | | 77.5 |
| 964 | PARAMETHADIONE | 85.15 | 83.0 | | 76.9 |
| 965 | PREDNISOLONE HEMISUCCINATE | 88.09 | 84.6 | | 80.0 |
| 966 | UNDECYLENIC ACID | 83.51 | 87.8 | | 75.7 |
| 967 | OLANZAPINE | 87.10 | 76.8 | | 79.2 |
| 968 | RIVASTIGMINE TARTRATE | 84.62 | 83.8 | | 78.3 |
| 969 | NISOLDIPINE | 78.41 | 84.6 | | 76.7 |
| 970 | BENURESTAT | 83.00 | 81.1 | | 77.8 |
| 971 | ETHOXZOLAMIDE | 85.53 | 77.9 | | 75.2 |
| 972 | HYDROCORTISONE VALERATE | 85.28 | 79.3 | | 75.4 |
| 973 | PREDNISOLONE TEBUTATE | 88.08 | 84.3 | | 77.0 |
| 974 | DISOPYRAMIDE PHOSPHATE | 92.14 | 83.7 | | 78.2 |
| 975 | BENOXINATE HYDROCHLORIDE | 108.84 | 84.8 | | 77.2 |
| 976 | BICALUTAMIDE | 101.98 | 82.2 | | 72.2 |
| 977 | MONTELUKAST SODIUM | 84.83 | 82.7 | | 79.6 |
| 978 | DECOQUINATE [5mM] | 88.79 | 81.9 | | 80.1 |
| 979 | ETHOTOIN | 81.94 | 85.8 | | 74.1 |
| 980 | LOBENDAZOLE | 87.34 | 79.4 | | 76.8 |
| 981 | PYRIDOXINE HYDROCHLORIDE | 82.44 | 80.9 | | 81.0 |
| 982 | CEFONICID SODIUM | 80.75 | 85.3 | | 79.7 |
| 983 | BETAXALOL HYDROCHLORIDE | 84.69 | 79.1 | | 76.2 |
| 984 | CYCLOSERINE (L) | 90.87 | 83.8 | | 78.9 |
| 985 | TERBINAFINE HYDROCHLORIDE | 141.29 | 114.5 | | 63.8 |
| 986 | BISMUTH SUBSALICYLATE | 85.36 | 77.9 | | 84.1 |
| 987 | FOSINOPRIL SODIUM | 90.01 | 82.5 | | 79.4 |
| 988 | METHSUXIMIDE | 84.83 | 75.4 | | 79.5 |
| 989 | PHENSUCCIMIDE | 82.74 | 83.0 | | 79.4 |
| 990 | IFOSFAMIDE | 87.72 | 83.9 | | 74.0 |
| 991 | BIPERIDEN | 96.01 | 82.7 | | 74.9 |
| 992 | PENBUTOLOL SULFATE | 95.45 | 90.2 | | 73.6 |
| 993 | DESLORATIDINE | 96.56 | 90.1 | | 75.8 |
| 994 | BENZOYLPAS | 87.22 | 81.1 | | 66.4 |
| 995 | DEFLAZACORT | 87.30 | 81.1 | | 76.1 |
| 996 | METHYLENE BLUE | 87.97 | 85.4 | | 82.0 |
| 997 | RIMANTADINE HYDROCHLORIDE | 91.18 | 86.6 | | 79.6 |
| 998 | DOXORUBICIN | 90.51 | 86.1 | | 88.5 |
| 999 | NEOSTIGMINE METHYLSULFATE | 79.98 | 77.6 | | 77.5 |
| 1000 | AMINOPENTAMIDE SULFATE | 89.27 | 81.6 | | 71.4 |
| 1001 | MOXIDECTIN | 94.49 | 84.5 | | 77.6 |
| 1002 | BROMINDIONE | 84.29 | 78.6 | | 76.3 |

| | | | | | |
|---|---|---|---|---|---|
| 1003 | FOMEPIZOLE HYDROCHLORIDE | 89.01 | 80.3 | | 76.7 |
| 1004 | METHYLATROPINE NITRATE | 86.50 | 80.9 | | 80.0 |
| 1005 | SULFISOXAZOLE ACETYL | 82.74 | 80.3 | | 76.2 |
| 1006 | CLOZAPINE | 85.92 | 82.2 | | 74.5 |
| 1007 | CITICOLINE | 87.22 | 83.6 | | 74.5 |
| 1008 | GENTAMICIN SULFATE | 88.49 | 82.2 | | 77.0 |
| 1009 | FLUORESCEIN | 102.37 | 103.6 | | 88.1 |
| 1010 | BURAMATE | 83.01 | 85.3 | | 78.5 |
| 1011 | GLIPIZIDE | 82.01 | 82.9 | | 76.6 |
| 1012 | NAFTIFINE HYDROCHLORIDE | 89.39 | 80.3 | | 78.7 |
| 1013 | SULISOBENZONE | 84.02 | 84.1 | | 78.4 |
| 1014 | PRULIFLOXACIN | 84.31 | 84.7 | | 95.0 |
| 1015 | DILOXANIDE FUROATE | 90.12 | 86.1 | | 75.0 |
| 1016 | FIPRONIL | 84.02 | 86.7 | | 72.7 |
| 1017 | NIACINAMIDE | 82.74 | 78.6 | | 76.1 |
| 1018 | CAPOBENIC ACID | 81.53 | 82.2 | | 75.1 |
| 1019 | HALOTHANE | 82.53 | 81.9 | | 84.2 |
| 1020 | MINOXIDIL | 82.49 | 80.7 | | 75.3 |
| 1021 | AMMONIUM LACTATE | 82.24 | 83.0 | | 83.9 |
| 1022 | HYDRALAZINE HYDROCHLORIDE | 83.55 | 70.5 | | 78.0 |
| 1023 | NADOLOL | 89.57 | 79.4 | | 78.9 |
| 1024 | GUANETHIDINE MONOSULFATE | 86.07 | 83.8 | | 71.1 |
| 1025 | PHENYLETHYL ALCOHOL | 87.05 | 83.3 | | 85.0 |
| 1026 | DEXPANTHENOL | 83.90 | 76.7 | | 79.2 |
| 1027 | GUANFACINE HYDROCHLORIDE | 80.96 | 81.2 | | 66.6 |
| 1028 | NITHIAMIDE | 81.32 | 81.7 | | 81.4 |
| 1029 | TRICLOSAN | 130.86 | 134.2 | | 107.9 |
| 1030 | SOLIFENACIN SUCCINATE | 83.55 | 91.7 | | 76.5 |
| 1031 | DYDROGESTERONE | 80.02 | 80.3 | | 79.8 |
| 1032 | ERGOTAMINE TARTRATE | 82.01 | 81.5 | | 74.6 |
| 1033 | OXYBUTYNIN CHLORIDE | 78.92 | 84.6 | | 83.1 |
| 1034 | ETHOPABATE | 82.99 | 83.0 | | 72.7 |
| 1035 | D-LACTITOL MONOHYDRATE | 83.20 | 80.1 | | 76.8 |
| 1036 | PRALIDOXIME CHLORIDE | 79.56 | 85.2 | | 74.7 |
| 1037 | TRIMETHADIONE | 80.92 | 82.3 | | 79.4 |
| 1038 | ACEPROMAZINE MALEATE | 81.79 | 81.2 | | 76.0 |
| 1039 | FENOLDOPAM MESYLATE | 82.51 | 82.5 | | 90.5 |
| 1040 | ANAGRELIDE HYDROCHLORIDE | 79.43 | 83.8 | | 78.7 |
| 1041 | FLUOROURACIL | 95.9 | 88.2 | | 84.7 |
| 1042 | ARSANILIC ACID | 93.2 | 91.6 | | 85.6 |
| 1043 | ANTIMONY POTASSIUM TARTRATE TRIHY | 96.7 | 87.7 | | 84.2 |
| 1044 | FLUDARABINE PHOSPHATE | 76.9 | 90.1 | | 82.2 |
| 1045 | RETINYL PALMITATE | 87.2 | 90.1 | | 81.1 |
| 1046 | ASCORBYL PALMITATE | 85.1 | 93.2 | | 81.3 |
| 1047 | TOPIRAMATE | 94.1 | 89.3 | | 79.4 |
| 1048 | IRINOTECAN HYDROCHLORIDE | 94.1 | 89.4 | | 75.6 |
| 1049 | ETOMIDATE | 91.3 | 86.6 | | 75.4 |
| 1050 | PANTHENOL (dl) | 89.0 | 83.0 | | 75.9 |
| 1051 | QUINAPRILAT | 93.8 | 85.7 | | 76.9 |
| 1052 | MUPIROCIN | 97.4 | 84.5 | | 76.9 |
| 1053 | CHLOROPHYLLIDE Cu COMPLEX Na SALT | 107.1 | 96.6 | | 80.6 |
| 1054 | ERYTHROSINE SODIUM | 107.6 | 105.2 | | 99.2 |
| 1055 | GEMIFLOXACIN MESYLATE | 96.8 | 86.0 | | 83.0 |
| 1056 | SITAGLIPTIN PHOSPHATE | 100.1 | 89.0 | | 79.5 |
| 1057 | FLORFENICOL | 92.6 | 93.8 | | 82.6 |
| 1058 | ANIRACETAM | 90.9 | 94.3 | | 80.6 |
| 1059 | PROPARACAINE HYDROCHLORIDE | 93.8 | 87.6 | | 81.2 |
| 1060 | TEICOPLANIN [A(2-1) shown] | 84.9 | 86.8 | | 80.2 |
| 1061 | DESOXYMETASONE | 90.4 | 85.3 | | 78.9 |
| 1062 | GLYCOPYRROLATE | 92.6 | 88.5 | | 73.0 |
| 1063 | PRAVASTATIN SODIUM | 93.8 | 86.2 | | 77.1 |
| 1064 | ZINC UNDECYLENATE [4mM] | 91.5 | 85.0 | | 77.8 |
| 1065 | FLUVOXAMINE MALEATE | 87.4 | 91.1 | | 80.3 |
| 1066 | VINCRISTINE SULFATE | 95.7 | 88.2 | | 75.9 |
| 1067 | PIPAMPERONE | 93.2 | 87.1 | | 76.2 |
| 1068 | EPIRUBICIN HYDROCHLORIDE | 90.1 | 88.1 | | 95.1 |
| 1069 | BETAMETHASONE ACETATE | 90.3 | 88.4 | | 77.6 |
| 1070 | CARTEOLOL HYDROCHLORIDE | 93.8 | 81.7 | | 77.1 |
| 1071 | GABAPENTIN | 94.6 | 87.1 | | 78.9 |
| 1072 | PIRENPERONE | 93.2 | 87.4 | | 81.3 |
| 1073 | LAMOTRIGINE | 90.2 | 90.5 | | 78.5 |
| 1074 | TRIENTINE HYDROCHLORIDE | 99.2 | 87.3 | | 81.7 |

| | | | | | |
|---|---|---|---|---|---|
| 1075 | PEFLOXACINE MESYLATE | 99.2 | 82.9 | | 79.8 |
| 1076 | VECURONIUM BROMIDE | 98.6 | 85.9 | | 79.9 |
| 1077 | BETAMETHASONE SODIUM PHOSPHATE | 93.2 | 83.3 | | 83.2 |
| 1078 | OCTISALATE | 90.5 | 74.7 | | 83.0 |
| 1079 | ALISKIREN HEMIFUMARATE | 98.8 | 89.5 | | 77.5 |
| 1080 | ARIPIPRAZOLE | 128.0 | 97.6 | | 81.5 |
| 1081 | MIFEPRISTONE | 94.9 | 89.0 | | 79.4 |
| 1082 | TICLOPIDINE HYDROCHLORIDE | 97.5 | 89.0 | | 76.3 |
| 1083 | NETILMICIN SULFATE | 93.2 | 87.8 | | 75.3 |
| 1084 | ACAMPROSATE CALCIUM | 95.0 | 86.6 | | 75.9 |
| 1085 | NATAMYCIN | 97.6 | 84.2 | | 75.6 |
| 1086 | TRICHLORFON | 92.0 | 86.0 | | 77.0 |
| 1087 | METFORMIN HYDROCHLORIDE | 95.6 | 85.8 | | 81.5 |
| 1088 | METHYCLOTHIAZIDE | 97.9 | 82.2 | | 76.2 |
| 1089 | RALOXIFENE HYDROCHLORIDE | 95.5 | 84.9 | | 84.4 |
| 1090 | TICARCILLIN DISODIUM | 92.6 | 86.9 | | 83.4 |
| 1091 | OMEPRAZOLE | 97.5 | 91.4 | | 82.4 |
| 1092 | PREDNISOLONE SODIUM PHOSPHATE | 94.3 | 88.4 | | 79.1 |
| 1093 | DESONIDE | 100.3 | 85.2 | | 75.4 |
| 1094 | DIATRIZOIC ACID | 97.1 | 88.5 | | 77.3 |
| 1095 | FLUOROMETHOLONE | 96.9 | 86.6 | | 90.1 |
| 1096 | ATRACURIUM BESYLATE | 251.8 | 170.0 | | 89.3 |
| 1097 | CEFPODOXIME PROXETIL | 94.3 | 86.4 | | 82.6 |
| 1098 | TETRAMIZOLE HYDROCHLORIDE | 95.0 | 88.8 | | 83.5 |
| 1099 | FIUMAZENIL | 95.0 | 86.2 | | 78.1 |
| 1100 | PREGNENOLONE SUCCINATE | 94.3 | 79.1 | | 86.8 |
| 1101 | MELENGESTROL ACETATE | 94.3 | 79.3 | | 79.8 |
| 1102 | BEPHENIUM HYDROXYNAPTHOATE | 98.0 | 84.3 | | 65.3 |
| 1103 | BUTYLATED HYDROXYANISOLE | 91.2 | 74.5 | | 82.2 |
| 1104 | EXEMESTANE | 96.8 | 76.2 | | 73.5 |
| 1105 | TADALAFIL | 95.6 | 91.0 | | 82.2 |
| 1106 | TOLTRAZURIL | 90.1 | 80.4 | | 87.0 |
| 1107 | ALTRENOGEST | 93.2 | 79.1 | | 79.1 |
| 1108 | ACADESINE | 96.7 | 81.7 | | 79.9 |
| 1109 | ENTACAPONE | 96.3 | 85.0 | | 82.1 |
| 1110 | MANIDIPINE HYDROCHLORIDE | 89.7 | 79.6 | | 93.6 |
| 1111 | LOVASTATIN | 99.2 | 84.9 | | 84.0 |
| 1112 | PROSCILLARIDIN A | 91.5 | 80.1 | | 75.8 |
| 1113 | CLORGILINE HYDROCHLORIDE | 93.2 | 87.5 | | 85.4 |
| 1114 | TIBOLONE | 90.8 | 84.3 | | 78.7 |
| 1115 | FINASTERIDE | 92.7 | 84.7 | | 77.7 |
| 1116 | MEGLUMINE | 92.6 | 87.3 | | 77.1 |
| 1117 | MIRTAZAPINE | 99.8 | 82.5 | | 80.8 |
| 1118 | 2,4-DINITROPHENOL | 84.2 | 81.7 | | 76.6 |
| 1119 | PROPIOLACTONE | 92.1 | 89.8 | | 83.6 |
| 1120 | CRYOFLURANE | 63.7 | 80.4 | | 76.2 |
| 1121 | ZILEUTON | 98.7 | 90.4 | | 71.4 |
| 1122 | CLOFAZIMINE | 255.0 | 175.7 | | 84.3 |
| 1123 | NIZATIDINE | 97.8 | 88.4 | | 70.1 |
| 1124 | REPAGLINIDE | 93.2 | 83.5 | | 76.7 |
| 1125 | PENTAGASTRIN | 91.5 | 84.9 | | 83.8 |
| 1126 | FELBINAC | 93.8 | 80.1 | | 85.2 |
| 1127 | IMEXON | 91.5 | 79.9 | | 81.2 |
| 1128 | THIOPENTAL SODIUM | 102.7 | 89.3 | | 68.3 |
| 1129 | METHYLPHENIDATE HYDROCHLORIDE | 93.8 | 87.3 | | 81.4 |
| 1130 | BENZYDAMINE HYDROCHLORIDE | 110.6 | 104.0 | | 83.7 |
| 1131 | DENATONIUM BENZOATE | 93.2 | 82.5 | | 82.3 |
| 1132 | RISPERIDONE | 91.4 | 78.4 | | 80.7 |
| 1133 | PROTIRELIN | 95.3 | 82.5 | | 82.9 |
| 1134 | FLUROXENE | 94.8 | 84.2 | | 80.8 |
| 1135 | PHYSOSTIGMINE SULFATE | 94.3 | 77.0 | | 78.8 |
| 1136 | TEMAZEPAM | 95.0 | 86.0 | | 78.4 |
| 1137 | ZALEPLON | 89.9 | 91.0 | | 69.1 |
| 1138 | DOXAZOSIN MESYLATE | 94.5 | 86.2 | | 71.4 |
| 1139 | DECAMETHONIUM BROMIDE | 87.2 | 83.0 | | 67.8 |
| 1140 | SOTALOL HYDROCHLORIDE | 91.4 | 85.7 | | 68.6 |
| 1141 | ALLYLISOTHIOCYANATE | 91.4 | 85.5 | | 78.7 |
| 1142 | ETHANOLAMINE OLEATE | 99.5 | 90.0 | | 83.0 |
| 1143 | MEGLUTOL | 89.2 | 86.6 | | 81.0 |
| 1144 | TAPENTADOL HYDROCHLORIDE | 88.2 | 87.2 | | 80.4 |
| 1145 | RABEPRAZOLE SODIUM | 88.5 | 86.4 | | 67.5 |
| 1146 | ISOETHARINE MESYLATE | 93.8 | 83.5 | | 69.2 |

| | | | | | |
|---|---|---|---|---|---|
| 1147 | MESALAMINE | 93.7 | 86.0 | | 67.0 |
| 1148 | AMINOPTERIN | 92.6 | 81.5 | | 80.8 |
| 1149 | ACESULFAME POTASSIUM | 93.8 | 83.0 | | 66.2 |
| 1150 | PHENOTHIAZINE | 96.6 | 96.0 | | 92.1 |
| 1151 | SYMCLOSENE | 89.2 | 79.3 | | 79.4 |
| 1152 | TILORONE | 91.0 | 74.8 | | 81.9 |
| 1153 | PAMABROM | 93.2 | 85.5 | | 74.6 |
| 1154 | FAMPRIDINE | 93.2 | 88.1 | | 69.4 |
| 1155 | ETHAMIVAN | 92.0 | 84.9 | | 70.6 |
| 1156 | BENAZEPRIL HYDROCHLORIDE | 123.1 | 114.8 | | 80.2 |
| 1157 | PANTOTHENIC ACID(d) Na salt | 85.4 | 82.2 | | 67.8 |
| 1158 | HYMECHROME | 82.5 | 83.0 | | 83.0 |
| 1159 | OCTOCRYLENE | 84.6 | 78.6 | | 68.6 |
| 1160 | CLOMIPRAMINE HYDROCHLORIDE | 96.5 | 99.9 | | 83.4 |
| 1161 | COLISTIN SULFATE | 291.8 | 280.6 | | 80.5 |
| 1162 | ETHYNODIOL DIACETATE | 85.7 | 78.8 | | 82.1 |
| 1163 | BUTYL PARABEN | 87.7 | 87.8 | | 84.0 |
| 1164 | ENILCONAZOLE SULFATE | 223.4 | 194.6 | | 75.3 |
| 1165 | MODALINE SULFATE | 84.6 | 83.3 | | 80.0 |
| 1166 | PIMAGEDINE HYDROCHLORIDE | 87.3 | 86.4 | | 79.3 |
| 1167 | TROCLOSENE SODIUM | 85.9 | 79.8 | | 81.5 |
| 1168 | CHLORMEZANONE | 83.5 | 84.9 | | 80.0 |
| 1169 | ARSENIC TRIOXIDE DIETHANOLAMINE SA | 87.4 | 89.9 | | 68.2 |
| 1170 | ORNIDAZOLE | 87.7 | 86.9 | | 71.4 |
| 1171 | IMIPENEM | 84.9 | 83.5 | | 67.7 |
| 1172 | DARIFENACIN HYDROBROMIDE | 88.1 | 84.5 | | 79.8 |
| 1173 | ADRENALONE HYDROCHLORIDE | 90.0 | 89.5 | | 72.8 |
| 1174 | ISOVALERAMIDE | 87.8 | 81.0 | | 77.6 |
| 1175 | ISOBUTAMBEN | 89.4 | 89.3 | | 76.6 |
| 1176 | AMOXAPINE | 100.6 | 104.9 | | 81.1 |
| 1177 | BENZBROMARONE | 83.5 | 79.9 | | 74.3 |
| 1178 | OXANTEL PAMOATE | 87.6 | 84.8 | | 66.3 |
| 1179 | METYRAPONE | 87.8 | 83.5 | | 72.3 |
| 1180 | BISOPROLOL FUMARATE | 84.7 | 85.7 | | 70.5 |
| 1181 | DIMETHYL FUMARATE | 83.3 | 88.7 | | 71.8 |
| 1182 | D-(+)-MALTOSE | 83.0 | 83.5 | | 80.5 |
| 1183 | ELETRIPTAN HYDROBROMIDE | 85.5 | 84.9 | | 77.8 |
| 1184 | TACRINE HYDROCHLORIDE | 85.4 | 83.0 | | 82.7 |
| 1185 | BROMPERIDOL | 143.8 | 157.9 | | 89.6 |
| 1186 | PHENFORMIN HYDROCHLORIDE | 82.5 | 86.2 | | 73.1 |
| 1187 | MOLINDONE HYDROCHLORIDE | 85.5 | 82.7 | | 73.1 |
| 1188 | PENTETIC ACID | 87.1 | 83.5 | | 84.3 |
| 1189 | NIKETHAMIDE | 86.6 | 83.8 | | 71.4 |
| 1190 | CASANTHRANOL [cascaroside A shown] | 85.6 | 83.8 | | 82.1 |
| 1191 | DIETHYLTOLUAMIDE | 87.5 | 85.9 | | 82.5 |
| 1192 | ACECAINIDE HYDROCHLORIDE | 83.8 | 87.7 | | 85.9 |
| 1193 | CYPROHEPTADINE HYDROCHLORIDE | 90.4 | 88.9 | | 81.0 |
| 1194 | PROTRYPTYLINE HYDROCHLORIDE | 87.0 | 97.1 | | 82.3 |
| 1195 | NICORANDIL | 86.4 | 86.6 | | 71.8 |
| 1196 | SULFADOXINE | 85.0 | 85.4 | | 76.5 |
| 1197 | TEMOZOLOMIDE | 82.7 | 84.1 | | 75.5 |
| 1198 | EPRODISATE DISODIUM | 85.6 | 86.6 | | 85.6 |
| 1199 | THIAMYLAL SODIUM | 85.8 | 86.2 | | 74.7 |
| 1200 | TOREMIFENE CITRATE | | | | |
| 1201 | THEOBROMINE | 93.7 | 90.8 | | 68.0 |
| 1202 | DEBRISOQUIN SULFATE | 90.3 | 185.7 | | 71.3 |
| 1203 | CAPTAMINE | 90.2 | 90.8 | | 69.5 |
| 1204 | ALPRENOLOL HYDROCHLORIDE | 88.5 | 90.2 | | 72.3 |
| 1205 | ZOLMITRIPTAN | 90.7 | 93.2 | | 89.3 |
| 1206 | MITOTANE | 90.2 | 90.7 | | 69.7 |
| 1207 | ISRADIPINE | 79.4 | 93.7 | | 70.0 |
| 1208 | NOCODAZOLE | 89.1 | 94.0 | | 79.5 |
| 1209 | BUSPIRONE HYDROCHLORIDE | 97.4 | 101.9 | | 72.6 |
| 1210 | NICOTINE BITARTRATE | 87.8 | 87.2 | | 71.1 |
| 1211 | TERPENE HYDRATE | 88.4 | 85.6 | | 71.5 |
| 1212 | ETHYL PARABEN | 86.0 | 86.9 | | 77.2 |
| 1213 | ORBIFLOXACIN | 92.5 | 85.9 | | 75.8 |
| 1214 | TEMEFOS | 69.5 | 87.1 | | 70.4 |
| 1215 | LACITOL | 89.9 | 86.6 | | 85.9 |
| 1216 | LIOTHYRONINE (L- isomer) SODIUM | 81.6 | 87.0 | | 63.5 |
| 1217 | VORTIOXETINE HYDROBROMIDE | 455.3 | 417.3 | | 89.0 |
| 1218 | VERAPAMIL HYDROCHLORIDE | 96.7 | 93.2 | | 78.4 |

| | | | | | |
|---|---|---|---|---|---|
| 1219 | SODIUM MONOFLUOROPHOSPHATE | 89.2 | 89.2 | | 75.1 |
| 1220 | EDETATE DISODIUM | 93.2 | 88.5 | | 75.0 |
| 1221 | IVERMECTIN | 101.9 | 103.3 | | 71.9 |
| 1222 | PERMETHRIN | 87.1 | 89.0 | | 72.6 |
| 1223 | PODOFILOX | 90.5 | 89.4 | | 73.7 |
| 1224 | CREATININE | 86.6 | 87.4 | | 70.0 |
| 1225 | LEVODOPA | 93.7 | 91.4 | | 73.8 |
| 1226 | METOLAZONE | 89.7 | 81.2 | | 76.6 |
| 1227 | TROSPIUM CHLORIDE | 90.2 | 92.7 | | 69.9 |
| 1228 | FLUTICASONE PROPIONATE | 88.8 | 89.6 | | 69.2 |
| 1229 | TROMETHAMINE | 92.1 | 92.7 | | 75.1 |
| 1230 | EDITOL | 75.5 | 93.7 | | 76.2 |
| 1231 | OMEGA-3-ACID ESTERS (EPA shown) | 87.0 | 92.2 | | 70.9 |
| 1232 | GABOXADOL HYDROCHLORIDE | 89.3 | 92.2 | | 70.3 |
| 1233 | DIAZOXIDE | 91.9 | 90.3 | | 74.1 |
| 1234 | PINACIDIL | 68.7 | 91.3 | | 76.8 |
| 1235 | AMIKACIN HYDRATE | 88.0 | 88.5 | | 74.0 |
| 1236 | AMIODARONE HYDROCHLORIDE | 447.0 | 271.3 | | 110.6 |
| 1237 | OXTRIPHYLLINE | 85.0 | 89.2 | | 70.2 |
| 1238 | ERYTHRITOL | 88.1 | 91.7 | | 70.9 |
| 1239 | RIZATRIPTAN BENZOATE | 84.8 | 94.3 | | 67.8 |
| 1240 | DEHYDROACETIC ACID | 82.2 | 85.1 | | 69.2 |
| 1241 | ITRACONAZOLE HYDROCHLORIDE | 138.4 | 154.6 | | 118.8 |
| 1242 | MIANSERIN HYDROCHLORIDE | 98.4 | 94.5 | | 71.9 |
| 1243 | PENICILLAMINE | 69.4 | 89.4 | | 68.7 |
| 1244 | ADENOSINE TRIPHOSPHATE DISODIUM | 84.5 | 87.2 | | 73.7 |
| 1245 | ESTRAMUSTINE | 80.8 | 92.4 | | 73.6 |
| 1246 | MALATHION | 83.5 | 89.7 | | 74.3 |
| 1247 | HYDROXYAMPHETAMINE HYDROBROMID | 85.4 | 90.1 | | 74.9 |
| 1248 | CYCLAMIC ACID | 84.0 | 88.5 | | 72.3 |
| 1249 | DILTIAZEM HYDROCHLORIDE | 92.7 | 94.8 | | 77.0 |
| 1250 | PENTYLENETETRAZOL | 91.7 | 92.1 | | 74.0 |
| 1251 | LINDANE | 91.5 | 96.0 | | 71.2 |
| 1252 | EVANS BLUE | 99.5 | 130.6 | | 86.6 |
| 1253 | ARTESUNATE | 88.6 | 83.4 | | 75.4 |
| 1254 | SELEGILINE HYDROCHLORIDE | 86.2 | 88.7 | | 72.6 |
| 1255 | ACRIFLAVINIUM HYDROCHLORIDE | 95.9 | 95.7 | | 77.3 |
| 1256 | RIBOFLAVIN 5-PHOSPHATE SODIUM | 80.5 | 85.6 | | 70.4 |
| 1257 | EDROPHONIUM CHLORIDE | 87.4 | 85.8 | | 76.8 |
| 1258 | TRANEXAMIC ACID | 86.9 | 89.1 | | 72.9 |
| 1259 | ACETOHYDROXAMIC ACID | 84.2 | 89.6 | | 77.5 |
| 1260 | LOPERAMIDE HYDROCHLORIDE | 92.1 | 97.1 | | 70.7 |
| 1261 | ACTINOQUINOL SODIUM | 86.8 | 89.0 | | 70.8 |
| 1262 | VIOMYCIN SULFATE | 88.9 | 90.7 | | 69.3 |
| 1263 | BUTYLATED HYDROXYTOLUENE | 84.5 | 88.4 | | 70.1 |
| 1264 | PRALIDOXIME MESYLATE | 86.6 | 88.9 | | 69.8 |
| 1265 | OXOLINIC ACID | 89.1 | 85.1 | | 71.9 |
| 1266 | CLONAZEPAM | 88.6 | 91.9 | | 69.3 |
| 1267 | IOTHALAMIC ACID | 86.2 | 89.1 | | 69.9 |
| 1268 | NALBUPHINE HYDROCHLORIDE | 88.2 | 83.4 | | 69.1 |
| 1269 | DULOXETINE HYDROCHLORIDE | 101.5 | 107.1 | | 70.0 |
| 1270 | NOMIFENSINE MALEATE | 85.9 | 91.4 | | 77.7 |
| 1271 | HEXYLENE GLYCOL | 83.9 | 91.3 | | 68.1 |
| 1272 | PHTHALYLSULFACETAMIDE | 85.9 | 90.2 | | 70.8 |
| 1273 | DACTINOMYCIN | 89.6 | 89.0 | | 72.6 |
| 1274 | AMYLENE HYDRATE | 84.0 | 91.3 | | 72.3 |
| 1275 | BEPRIDIL HYDROCHLORIDE | 79.6 | 89.9 | | 72.0 |
| 1276 | DIFLORASONE DIACETATE | 82.3 | 88.5 | | 79.4 |
| 1277 | AUROTHIOGLUCOSE | 86.4 | 88.7 | | 73.9 |
| 1278 | CALCIUM GLUCEPTATE | 83.7 | 89.1 | | 70.4 |
| 1279 | ETHYL VANILLIN | 81.6 | 90.2 | | 71.9 |
| 1280 | SILDENAFIL CITRATE | 89.5 | 91.9 | | 70.9 |
| 1281 | SULFACARBAMIDE | 86.2 | 91.2 | | 86.5 |
| 1282 | RUTIN | 90.0 | 88.1 | | 81.5 |
| 1283 | ACEGLUTAMIDE | 86.4 | 88.0 | | 80.5 |
| 1284 | HEXESTROL | 85.4 | 85.5 | | 83.0 |
| 1285 | DIBEKACIN | 84.7 | 86.2 | | 81.8 |
| 1286 | CINCHONIDINE | 87.1 | 88.1 | | 80.7 |
| 1287 | SULFAPHENAZOLE | 88.0 | 84.4 | | 79.8 |
| 1288 | RIBOSTAMYCIN SULFATE | 82.0 | 88.4 | | 71.1 |
| 1289 | GALLIC ACID | 84.4 | 83.4 | | 82.0 |
| 1290 | PIPEMIDIC ACID | 85.2 | 84.9 | | 78.6 |

| | | | | | |
|---|---|---|---|---|---|
| 1291 | CLOFIBRIC ACID | 85.2 | 80.2 | | 83.6 |
| 1292 | HEXETIDINE | 328.1 | 299.1 | | 103.9 |
| 1293 | ACEMETACIN | 83.7 | 85.8 | | 85.7 |
| 1294 | CINCHONINE | 83.5 | 85.1 | | 85.5 |
| 1295 | SULFAGUANIDINE | 86.7 | 84.7 | | 80.5 |
| 1296 | ACETYL-L-LEUCINE | 83.7 | 86.8 | | 82.9 |
| 1297 | KHELLIN | 87.3 | 84.4 | | 86.1 |
| 1298 | HYDRASTININE HYDROCHLORIDE | 87.7 | 88.8 | | 81.5 |
| 1299 | ERYTHROMYCIN STEARATE | 75.4 | 89.7 | | 81.1 |
| 1300 | IPRONIAZID PHOSPHATE | 90.1 | 89.5 | | 75.1 |
| 1301 | BENFLUOREX HYDROCHLORIDE | 97.5 | 100.3 | | 82.9 |
| 1302 | INDINAVIR SULFATE | 87.4 | 85.5 | | 74.9 |
| 1303 | D-PHENYLALANINE | 87.4 | 85.6 | | 66.7 |
| 1304 | BUFEXAMAC | 87.4 | 90.2 | | 62.0 |
| 1305 | DEQUALINIUM CHLORIDE | 120.7 | 337.8 | | 134.1 |
| 1306 | QUININE ETHYL CARBONATE | 87.7 | 90.5 | | 83.6 |
| 1307 | ACETARSOL | 85.5 | 85.9 | | 77.8 |
| 1308 | MECYSTEINE HYDROCHLORIDE | 84.1 | 89.1 | | 73.1 |
| 1309 | gamma-AMINOBUTYRIC ACID HYDROCHL | 83.5 | 45.3 | | 77.3 |
| 1310 | GLAFENINE | 83.7 | 87.4 | | 73.6 |
| 1311 | CIPROFIBRATE | 81.9 | 96.0 | | 74.4 |
| 1312 | DIACERIN | 87.4 | 91.6 | | 84.4 |
| 1313 | HYDROXYTOLUIC ACID | 90.2 | 88.5 | | 84.4 |
| 1314 | COUMOPHOS | 80.4 | 82.5 | | 82.3 |
| 1315 | TOLONIUM CHLORIDE | 89.4 | 96.1 | | 88.1 |
| 1316 | OCTOPAMINE HYDROCHLORIDE | 85.9 | 87.7 | | 81.6 |
| 1317 | PRASUGREL | 84.0 | 86.6 | | 77.4 |
| 1318 | DROFENINE HYDROCHLORIDE | 95.9 | 99.1 | | 82.0 |
| 1319 | ACETANILIDE | 82.2 | 80.3 | | 81.2 |
| 1320 | OXIRACETAM | 85.4 | 81.5 | | 79.2 |
| 1321 | CHLORQUINALDOL | 86.9 | 87.5 | | 84.1 |
| 1322 | XYLOSE | 82.2 | 85.9 | | 75.7 |
| 1323 | AMINOPYRINE | 85.5 | 86.9 | | 81.3 |
| 1324 | PENTAMIDINE ISETHIONATE | 115.8 | 131.4 | | 70.7 |
| 1325 | LOBELINE HYDROCHLORIDE | 62.0 | 88.3 | | 82.3 |
| 1326 | ETHAVERINE HYDROCHLORIDE | 91.9 | 102.9 | | 85.6 |
| 1327 | TODRALAZINE HYDROCHLORIDE | 84.0 | 87.2 | | 70.5 |
| 1328 | NIFLUMIC ACID | 79.8 | 89.6 | | 85.1 |
| 1329 | PRASTERONE ACETATE | 109.7 | 105.8 | | 82.4 |
| 1330 | CHLORPYRIFOS | 87.2 | 90.8 | | 83.2 |
| 1331 | BROXYQUINOLINE | 192.6 | 136.8 | | 83.5 |
| 1332 | PREGNENOLONE | 75.1 | 107.3 | | 64.4 |
| 1333 | BERBERINE CHLORIDE | 80.7 | 98.0 | | 97.4 |
| 1334 | DROPROPIZINE | 84.0 | 89.8 | | 72.3 |
| 1335 | EBSELEN | | | | |
| 1336 | TENOFOVIR | 84.9 | 89.0 | | 85.0 |
| 1337 | BERGAPTEN | 89.7 | 97.4 | | 88.9 |
| 1338 | FENTHION | 78.0 | 90.6 | | 86.5 |
| 1339 | CARNITINE (dl) HYDROCHLORIDE | 67.5 | 88.3 | | 84.7 |
| 1340 | SULFANILAMIDE | 64.9 | 86.1 | | 78.9 |
| 1341 | BENFOTIAMINE | 81.3 | 85.2 | | 82.0 |
| 1342 | FENDILINE HYDROCHLORIDE | 94.9 | 101.1 | | 75.8 |
| 1343 | SODIUM TETRADECYL SULFATE | 293.6 | 288.9 | | 89.3 |
| 1344 | ATROPINE OXIDE | 85.2 | 86.2 | | 81.6 |
| 1345 | TRICHLORMETHINE HYDROCHLORIDE | 86.1 | 88.8 | | 83.4 |
| 1346 | PROPOXUR | 85.9 | 86.1 | | 82.4 |
| 1347 | DICHLOROPHEN | 87.9 | 100.4 | | 92.5 |
| 1348 | VINCAMINE | 85.7 | 87.7 | | 80.7 |
| 1349 | BRUCINE | 86.2 | 86.4 | | 80.8 |
| 1350 | PROTOPORPHYRIN IX | 100.0 | 97.2 | | 92.6 |
| 1351 | TOLPERISONE HYDROCHLORIDE | 86.8 | 86.2 | | 73.8 |
| 1352 | COUMARIN | 86.6 | 86.8 | | 72.0 |
| 1353 | MEVASTATIN | 85.2 | 87.4 | | 84.8 |
| 1354 | IPRIFLAVONE | 61.2 | 92.1 | | 85.9 |
| 1355 | FLOPROPIONE | 83.9 | 88.1 | | 84.0 |
| 1356 | AJMALINE | 82.7 | 88.2 | | 77.5 |
| 1357 | CEPHALOSPORIN C Zn | 83.9 | 88.2 | | 87.8 |
| 1358 | MEBHYDROLIN | 81.9 | 90.2 | | 84.8 |
| 1359 | TOLFENAMIC ACID | 81.9 | 90.1 | | 90.5 |
| 1360 | ELLAGIC ACID | 86.1 | 88.0 | | 85.9 |
| 1361 | IOVERSOL | 98.6 | 88.4 | | 85.5 |
| 1362 | PIDOTIMOD | 97.8 | 90.5 | | 82.2 |

| | | | | | |
|---|---|---|---|---|---|
| 1363 | SODIUM CYCLAMATE | 96.9 | 89.8 | | 85.9 |
| 1364 | BORNYL ACETATE | 92.2 | 88.2 | | 83.3 |
| 1365 | BUCLADESINE SODIUM | 98.9 | 86.8 | | 84.7 |
| 1366 | URAPIDIL HYDROCHLORIDE | 96.6 | 91.0 | | 84.6 |
| 1367 | MECLOFENOXATE HYDROCHLORIDE | 96.3 | 88.8 | | 82.1 |
| 1368 | ALOIN | 94.2 | 87.3 | | 83.9 |
| 1369 | RAMOPLANIN [A2 shown; 2mM] | 93.6 | 94.3 | | 84.0 |
| 1370 | PRANOPROFEN | 88.3 | 88.8 | | 84.4 |
| 1371 | CLOFOCTOL | 82.1 | 76.8 | | 87.0 |
| 1372 | CHLOROBUTANOL | 95.6 | 89.0 | | 74.4 |
| 1373 | CHROMOCARB | 94.2 | 82.2 | | 85.6 |
| 1374 | BROMOPRIDE | 94.7 | 85.0 | | 86.6 |
| 1375 | NIFENAZONE | 95.5 | 86.4 | | 84.7 |
| 1376 | SALINOMYCIN, SODIUM | 98.7 | 99.3 | | 86.6 |
| 1377 | PYRONARIDINE TETRAPHOSPHATE | 98.5 | 90.6 | | 83.8 |
| 1378 | BRONOPOL | 96.9 | 91.5 | | 83.7 |
| 1379 | PALMIDROL [5mM] | 98.1 | 94.4 | | 84.2 |
| 1380 | OXALIPLATIN | 94.4 | 84.4 | | 85.9 |
| 1381 | ACEXAMIC ACID | 92.8 | 85.9 | | 83.7 |
| 1382 | LEVOMENTHOL | 95.2 | 88.5 | | 69.7 |
| 1383 | NIMESULIDE | 94.2 | 89.3 | | 83.9 |
| 1384 | CYPERMETHRIN | 90.3 | 85.3 | | 83.1 |
| 1385 | DIMESNA | 89.8 | 92.3 | | 84.7 |
| 1386 | RACECADOTRIL | 89.1 | 87.3 | | 85.1 |
| 1387 | CYACETACIDE | 92.7 | 86.0 | | 77.8 |
| 1388 | ETHENZAMIDE | 89.9 | 86.4 | | 85.0 |
| 1389 | RETINYL ACETATE | 86.1 | 86.1 | | 85.2 |
| 1390 | CETRIMONIUM BROMIDE | | | | |
| 1391 | HEXAMETHONIUM BROMIDE | 85.6 | 81.9 | | 83.2 |
| 1392 | PRONETALOL HYDROCHLORIDE | 93.7 | 89.9 | | 86.7 |
| 1393 | LOMERIZINE HYDROCHLORIDE | 95.5 | 99.3 | | 87.4 |
| 1394 | DOCOSANOL | 88.7 | 88.2 | | 72.4 |
| 1395 | ACECLOFENAC | 82.7 | 88.9 | | 65.0 |
| 1396 | BENZYL ISOTHIOCYANATE | 186.4 | 216.4 | | 88.2 |
| 1397 | PIPENZOLATE BROMIDE | 79.8 | 87.8 | | 86.4 |
| 1398 | CITIOLONE | 84.7 | 70.9 | | 85.5 |
| 1399 | TIMONACIC | 88.7 | 91.5 | | 84.2 |
| 1400 | BRIVUDINE | 85.6 | 85.0 | | 83.5 |
| 1401 | MOGUISTEINE | 81.1 | 84.2 | | 83.8 |
| 1402 | TENATOPRAZOLE | 85.1 | 86.8 | | 81.9 |
| 1403 | FERROUS FUMARATE | 84.4 | 90.2 | | 87.1 |
| 1404 | AMINOTHIAZOLE | 84.8 | 90.6 | | 84.4 |
| 1405 | NEVIRAPINE | 89.1 | 82.7 | | 84.1 |
| 1406 | AMPYZINE SULFATE | 87.6 | 87.7 | | 89.6 |
| 1407 | THIODIGLYCOL | 70.0 | 83.8 | | 88.1 |
| 1408 | EDARAVONE | 86.7 | 81.9 | | 89.5 |
| 1409 | NADIFLOXACIN | 89.6 | 92.7 | | 85.8 |
| 1410 | TROPISETRON HYDROCHLORIDE | 87.3 | 93.2 | | 87.2 |
| 1411 | LOXOPROFEN | 83.8 | 85.9 | | 85.8 |
| 1412 | DIPYROCETYL | 81.7 | 69.3 | | 87.7 |
| 1413 | PRIDINOL METHANESULFONATE | 82.1 | 86.7 | | 85.5 |
| 1414 | HEPTAMINOL HYDROCHLORIDE | 88.6 | 97.9 | | 83.6 |
| 1415 | PASINIAZID | 86.1 | 90.7 | | 84.4 |
| 1416 | METITEPINE MESYLATE | | | | 151.2 |
| 1417 | NEFIRACETAM | 84.8 | 88.0 | | 87.1 |
| 1418 | DIMINAZENE ACETURATE | 89.4 | 87.4 | | 85.4 |
| 1419 | ACEDOBEN | 85.7 | 86.9 | | 89.9 |
| 1420 | DINITOLMIDE | 83.9 | 92.3 | | 86.8 |
| 1421 | AMBROXOL HYDROCHLORIDE | 85.6 | 82.0 | | 92.1 |
| 1422 | DIOSMIN | 81.7 | 87.4 | | 86.4 |
| 1423 | PEMPIDINE TARTRATE | 83.4 | 81.0 | | 89.1 |
| 1424 | PREDNICARBATE | 85.9 | 88.8 | | 86.7 |
| 1425 | OLTIPRAZ | 87.6 | 99.7 | | 82.7 |
| 1426 | FASUDIL HYDROCHLORIDE | 81.8 | 86.2 | | 86.6 |
| 1427 | ACTARIT | 75.0 | 90.7 | | 84.4 |
| 1428 | THREONINE (dl) | 84.3 | 86.7 | | 76.3 |
| 1429 | PYRITHYLDIONE | 83.5 | 85.2 | | 87.1 |
| 1430 | FIPEXIDE HYDROCHLORIDE | 86.2 | 101.3 | | 86.1 |
| 1431 | EXALAMIDE | 74.1 | 84.4 | | 90.7 |
| 1432 | ORNITHINE HYDROCHLORIDE | 88.1 | 79.9 | | 85.4 |
| 1433 | PAZUFLOXACIN MESYLATE | 86.4 | 86.1 | | 58.2 |
| 1434 | ITOPRIDE HYDROCHLORIDE | 83.8 | 94.8 |  | 89.0 |

| | | | | | |
|---|---|---|---|---|---|
| 1435 | EPALRESTAT | 84.7 | 94.0 | | 89.3 |
| 1436 | ARTEMISININ | 107.1 | 116.8 | | 76.8 |
| 1437 | MILTEFOSINE [5mM] | | 201.2 | | 97.0 |
| 1438 | DOCETAXEL | | 84.4 | | 87.6 |
| 1439 | AMEZINIUM METILSULFATE | | 88.0 | | 89.7 |
| 1440 | TIOXOLONE | | 92.1 | | 88.9 |
| 1441 | CHLORMIDAZOLE | 114.7 | 121.9 | | 83.9 |
| 1442 | GATIFLOXACIN | 86.4 | 89.7 | | 85.9 |
| 1443 | CYTISINE | 87.6 | 83.8 | | 84.8 |
| 1444 | PIPERONYL BUTOXIDE | 85.6 | 84.9 | | 84.7 |
| 1445 | FTAXILIDE | 82.3 | 83.3 | | 83.9 |
| 1446 | CHINIOFON | 92.7 | 90.9 | | 81.5 |
| 1447 | MOROXYDINE HYDROCHLORIDE | 85.6 | 85.7 | | 80.8 |
| 1448 | OXEDRINE | 87.7 | 91.0 | | 83.0 |
| 1449 | ENOXOLONE | 94.7 | 93.2 | | 84.1 |
| 1450 | TAMSULOSIN HYDROCHLORIDE | 86.5 | 78.5 | | 84.1 |
| 1451 | ESCIN | 84.4 | 83.5 | | 82.0 |
| 1452 | ZOXAZOLAMINE | 89.1 | 83.6 | | 86.6 |
| 1453 | beta-NAPHTHOL | 89.3 | 85.1 | | 83.7 |
| 1454 | CHLORALOSE | 83.0 | 88.1 | | 84.6 |
| 1455 | OROTIC ACID | 75.3 | 83.5 | | 84.4 |
| 1456 | SULBENTINE | 150.2 | 173.8 | | 107.3 |
| 1457 | BROXALDINE | 146.8 | 125.8 | | 141.2 |
| 1458 | DEXFOSFOSERINE HYDROCHLORIDE | 86.7 | 91.6 | | 81.7 |
| 1459 | PHENOTHRIN | 84.0 | 79.4 | | 80.9 |
| 1460 | BETAMIPRON | 88.1 | 82.5 | | 81.4 |
| 1461 | BITOSCANATE | 85.2 | 83.0 | | 83.9 |
| 1462 | DEFERIPRONE | 75.6 | 87.2 | | 85.6 |
| 1463 | TIOGUANINE | 84.9 | 81.5 | | 88.8 |
| 1464 | PRASTERONE | 103.6 | 95.8 | | 84.4 |
| 1465 | CHLORAZANIL HYDROCHLORIDE | 89.0 | 79.7 | | 77.5 |
| 1466 | CAPRYILIDENE | 87.2 | 84.5 | | 87.6 |
| 1467 | METHOPRENE (S) | 87.2 | 82.0 | | 83.0 |
| 1468 | PROFLAVINE HEMISULFATE | 107.8 | 112.2 | | 111.2 |
| 1469 | REBAMIPIDE | 89.5 | 79.6 | | 81.6 |
| 1470 | AFALANINE | 88.7 | 85.4 | | 84.9 |
| 1471 | MOXISYLYTE HYDROCHLORIDE | 88.0 | 82.7 | | 85.0 |
| 1472 | NAFTOPIDIL | 100.7 | 93.9 | | 80.6 |
| 1473 | HOMIDIUM BROMIDE | 100.8 | 98.2 | | 94.7 |
| 1474 | FENACLON | 89.5 | 85.7 | | 78.3 |
| 1475 | FORMESTANE | 89.8 | 65.6 | | 78.0 |
| 1476 | PERINDOPRILAT | 90.4 | 83.7 | | 83.4 |
| 1477 | CARSALAM | 85.6 | 88.9 | | 82.0 |
| 1478 | TYLOXAPOL | 100.0 | 83.3 | | 82.5 |
| 1479 | ISAXONINE | 89.5 | 87.1 | | 82.1 |
| 1480 | ETHACRIDINE LACTATE | 97.7 | 99.3 | | 97.3 |
| 1481 | RAMIFENAZONE | 94.5 | 85.9 | | 82.3 |
| 1482 | CLENBUTEROL HYDROCHLORIDE | 91.4 | 83.7 | | 79.9 |
| 1483 | ANCITABINE HYDROCHLORIDE | 92.3 | 82.5 | | 81.2 |
| 1484 | CYCLANDELATE | 91.0 | 85.9 | | 79.3 |
| 1485 | CARZENIDE | 91.2 | 80.3 | | 77.0 |
| 1486 | DIACETAMATE | 91.7 | 79.7 | | 86.1 |
| 1487 | AMPIROXICAM | 88.4 | 81.0 | | 82.5 |
| 1488 | BAMBUTEROL HYDROCHLORIDE | 91.5 | 86.8 | | 80.9 |
| 1489 | PIROMIDIC ACID | 96.6 | 84.4 | | 83.0 |
| 1490 | CLOPERASTINE HYDROCHLORIDE | 99.1 | 97.1 | | 84.0 |
| 1491 | PROTIONAMIDE | 92.1 | 88.1 | | 83.4 |
| 1492 | TIOCONAZOLE | 831.7 | | | |
| 1493 | CHLORAMINE-T | 94.5 | 84.5 | | 83.0 |
| 1494 | IRSOGLADINE MALEATE | 94.4 | 80.5 | | 68.9 |
| 1495 | ACIPIMOX | 87.8 | 83.0 | | 80.4 |
| 1496 | TROXERUTIN | 91.2 | 85.7 | | 84.4 |
| 1497 | EPROBEMIDE | 84.8 | 88.7 | | 81.6 |
| 1498 | CYPROTERONE | 85.0 | 84.6 | | 79.9 |
| 1499 | GLICLAZIDE | 88.0 | 87.5 | | 76.3 |
| 1500 | CARBARIL | 89.2 | 86.6 | | 81.9 |
| 1501 | CHLORINDIONE | 84.7 | 83.7 | | 73.8 |
| 1502 | EFLOXATE | 86.4 | 82.0 | | 80.8 |
| 1503 | CLIMBAZOLE | 148.8 | 251.5 | | 127.0 |
| 1504 | OXOLAMINE CITRATE | 87.3 | 86.2 | | 83.5 |
| 1505 | METERGOLINE | 103.3 | 121.5 | | 80.7 |
| 1506 | THIOCTIC ACID | 81.4 | 84.7 | | 82.5 |

| | | | | | |
|---|---|---|---|---|---|
| 1507 | NIMUSTINE HYDROCHLORIDE | 80.8 | 82.2 | | 83.0 |
| 1508 | VANITIOLIDE | 84.7 | 85.5 | | 85.2 |
| 1509 | CINCHOPHEN | 84.8 | 87.7 | | 85.7 |
| 1510 | EPIESTRIOL | 83.7 | 83.7 | | 82.4 |
| 1511 | RONNEL | 81.7 | 80.3 | | 83.2 |
| 1512 | OUABAIN OCTAHYDRATE | 81.2 | 92.4 | | 85.1 |
| 1513 | ETOSALAMIDE | 78.7 | 84.4 | | 79.6 |
| 1514 | TULOBUTEROL HYDROCHLORIDE | 83.3 | 84.6 | | 81.6 |
| 1515 | CARMUSTINE | 81.0 | 78.5 | | 79.8 |
| 1516 | ETEBENECID | 82.1 | 83.2 | | 77.0 |
| 1517 | TRIMEBUTINE MALEATE | 80.6 | 84.7 | | 76.6 |
| 1518 | SECNIDAZOLE | 79.9 | 83.4 | | 86.9 |
| 1519 | TRIMETAZIDINE DIHYDROCHLORIDE | 83.1 | 88.6 | | 82.7 |
| 1520 | ADIPIC ACID | 79.6 | 84.8 | | 82.2 |
| 1521 | IDRAMANTONE | 86.4 | 88.5 | | 80.0 |
| 1522 | NITROXOLINE | 334.1 | 231.4 | | 115.8 |
| 1523 | meta-CRESYL ACETATE | 86.2 | 86.7 | | 79.6 |
| 1524 | PIDOLIC ACID | 91.1 | 83.9 | | 79.1 |
| 1525 | BENZOCLIDINE | 87.4 | 89.1 | | 80.3 |
| 1526 | TIOPRONIN | 86.0 | 97.4 | | 80.0 |
| 1527 | MEPIROXOL | 85.5 | 88.1 | | 89.9 |
| 1528 | CEPHARANTHINE | 89.6 | 88.0 | | 80.7 |
| 1529 | CARGLUMIC ACID | 84.4 | 83.3 | | 78.8 |
| 1530 | CHLOROPYRAMINE HYDROCHLORIDE | 94.8 | 90.5 | | 79.8 |
| 1531 | CYROMAZINE | 81.4 | 83.9 | | 79.6 |
| 1532 | IDAZOXAN HYDROCHLORIDE | 88.8 | 85.9 | | 79.7 |
| 1533 | ADELMIDROL | 80.3 | 90.9 | | 77.4 |
| 1534 | EFAROXAN HYDROCHLORIDE | 90.2 | 87.0 | | 81.8 |
| 1535 | CRESOPIRINE | 82.2 | 85.1 | | 81.5 |
| 1536 | PROCODAZOLE | 87.8 | 87.4 | | 80.9 |
| 1537 | CARBIMAZOLE | 92.2 | 86.7 | | 80.5 |
| 1538 | FAMPROFAZONE | 86.2 | 85.8 | | 80.1 |
| 1539 | PROXYPHYLLINE | 85.2 | 85.4 | | 80.0 |
| 1540 | PICOLAMINE | 91.2 | 85.9 | | 79.7 |
| 1541 | BUCETIN | 87.4 | 84.4 | | 79.5 |
| 1542 | N-METHYL (-)EPHEDRINE [1R,2S] | 88.4 | 109.2 | | 78.1 |
| 1543 | PHENOLSULFONPHTHALEIN | 87.7 | 87.2 | | 82.6 |
| 1544 | CINOCTRAMIDE | 90.5 | 86.0 | | 86.2 |
| 1545 | SECURININE | 97.2 | 97.9 | | 75.7 |
| 1546 | OXELAIDIN CITRATE | 74.8 | 87.2 | | 83.0 |
| 1547 | PANTETHINE | 84.6 | 87.0 | | 81.8 |
| 1548 | OXINIACIC ACID | 91.7 | 83.7 | | 78.6 |
| 1549 | BARBITAL | 89.5 | 108.2 | | 81.4 |
| 1550 | INOSINE | 87.6 | 85.9 | | 79.3 |
| 1551 | GUANIDINE HYDROCHLORIDE | 80.9 | 85.3 | | 80.8 |
| 1552 | FURALTADONE | 90.2 | 86.5 | | 82.9 |
| 1553 | ERDOSTEINE | | 85.8 | | 77.5 |
| 1554 | DICHLORISONE ACETATE | 87.5 | 88.0 | | 83.5 |
| 1555 | SUCRALOSE | 87.7 | 87.4 | | 80.1 |
| 1556 | DOCONEXENT | 86.9 | 84.2 | | 81.0 |
| 1557 | CORTISONE | 86.9 | 83.4 | | 80.8 |
| 1558 | PYRETHRINS | 86.2 | 102.1 | | 76.5 |
| 1559 | GUAIACOL | 90.7 | 86.2 | | 79.9 |
| 1560 | MEXENEONE | 87.1 | 82.2 | | 82.2 |
| 1561 | LUFENURON | 79.8 | 85.5 | | 79.6 |
| 1562 | OXYPHENONIUM BROMIDE | 84.2 | 84.0 | | 81.9 |
| 1563 | NIALAMIDE | 83.0 | 85.6 | | 80.7 |
| 1564 | MILRINONE | 90.5 | 84.6 | | 78.8 |
| 1565 | d-LIMONENE | 85.9 | 120.2 | | 77.0 |
| 1566 | THIOPHANATE METHYL | 86.2 | 84.2 | | 79.2 |
| 1567 | SILIBININ | 85.6 | 85.9 | | 79.7 |
| 1568 | METICRANE | 86.8 | 85.6 | | 78.8 |
| 1569 | BENZONATATE | 88.0 | 87.1 | | 78.0 |
| 1570 | CAMYLOFINE DIHYDROCHLORIDE | 83.0 | 85.1 | | 79.7 |
| 1571 | CLEMIZOLE HYDROCHLORIDE | 81.1 | 85.2 | | 80.0 |
| 1572 | DIBUTYL PHTHALATE | 83.8 | 81.6 | | 80.2 |
| 1573 | HYDROQUINIDINE | 85.1 | 86.9 | | 81.7 |
| 1574 | METACETAMOL | 88.0 | 88.8 | | 79.1 |
| 1575 | ALLYLTHIOUREA | 86.2 | 84.4 | | 76.4 |
| 1576 | MEPARFYLON | 86.9 | 91.6 | | 77.6 |
| 1577 | LIPOAMIDE | 88.0 | 86.2 | | 82.5 |
| 1578 | TRICLABENDAZOLE | 95.7 | 102.4 | | 114.1 |

| | | | | | |
|---|---|---|---|---|---|
| 1579 | BUFLOMEDIL HYDROCHLORIDE | 84.4 | 84.4 | | 83.2 |
| 1580 | TRICHLOROETHYLENE | 81.0 | 82.7 | | 82.9 |
| 1581 | PIMETHIXENE MALEATE | 95.6 | 94.5 | | 79.5 |
| 1582 | LEVOCARNITINE PROPIONATE HYDROCHI | 84.8 | 87.2 | | 80.4 |
| 1583 | DIMETRIDAZOLE | 82.3 | 84.4 | | 79.6 |
| 1584 | PICONOL | 84.6 | 84.5 | | 78.4 |
| 1585 | IDEBENONE | 85.9 | 83.6 | | 76.9 |
| 1586 | NIFUROXAZIDE | 81.4 | 83.6 | | 80.5 |
| 1587 | CEFALONIUM | 81.7 | 83.3 | | 79.8 |
| 1588 | TIRATRICOL | 81.3 | 82.5 | | 82.2 |
| 1589 | ARTENIMOL | 136.0 | 109.6 | | 118.1 |
| 1590 | 2-THIOURACIL | 86.4 | 84.0 | | 78.2 |
| 1591 | NONIVAMIDE | 80.7 | 85.6 | | 76.0 |
| 1592 | MENBUTONE | 85.2 | 87.2 | | 81.7 |
| 1593 | PYRITINOL | 83.0 | 80.2 | | 83.3 |
| 1594 | BECLAMIDE | 82.1 | 82.7 | | 83.5 |
| 1595 | NANOFIN | 82.7 | 83.0 | | 82.0 |
| 1596 | ANISODAMINE HYDROBROMIDE | 77.3 | 85.9 | | 82.9 |
| 1597 | GENISTEIN | 80.9 | 84.7 | | 80.5 |
| 1598 | PAROXYPROPIONE | 82.5 | 87.2 | | 81.4 |
| 1599 | CARMOFUR | 97.0 | 97.2 | | 82.7 |
| 1600 | HYDROXYPROGESTERONE | 82.5 | 83.9 | | 82.6 |
| 1601 | SODIUM PHENYLBUTYRATE | 82.2 | 84.9 | | 77.0 |
| 1602 | THYMOPENTIN | 86.2 | 82.0 | | 81.9 |
| 1603 | TRIAMCINOLONE DIACETATE | 83.0 | 79.9 | | 85.0 |
| 1604 | ABAMECTIN (avermectin B1a shown) | 99.7 | 88.0 | | 90.7 |
| 1605 | NICOPHOLINE | 87.0 | 84.2 | | 80.7 |
| 1606 | OLOPATADINE HYDROCHLORIDE | 85.1 | 82.0 | | 79.1 |
| 1607 | LEUCINE (L) | 86.7 | 84.9 | | 78.8 |
| 1608 | VOGLIBOSE | 85.3 | 84.5 | | 72.9 |
| 1609 | CHLORINDANOL | 92.1 | 91.6 | | 75.8 |
| 1610 | DEXTROSE | 86.9 | 84.4 | | 81.7 |
| 1611 | ROFLUMILAST | 93.2 | 84.4 | | 86.8 |
| 1612 | CILNIDIPINE | 81.4 | 77.1 | | 79.5 |
| 1613 | CYSTEINE HYDROCHLORIDE | 87.7 | 81.1 | | 78.4 |
| 1614 | D-ERYTHOBIC ACID | 84.4 | 81.2 | | 76.7 |
| 1615 | CALCIUM CHLORIDE | 86.5 | 71.3 | | 80.2 |
| 1616 | GLUTAMINE (L) HYDROCHLORIDE | 85.9 | 83.2 | | 84.2 |
| 1617 | GLYCERIN | 86.6 | 87.4 | | 79.4 |
| 1618 | TROXIPIDE | 88.4 | 83.6 | | 79.7 |
| 1619 | ASPARTIC ACID (L) | 86.6 | 78.3 | | 79.0 |
| 1620 | ASPARAGINE (L) HYDRATE | 88.6 | 83.0 | | 78.5 |
| 1621 | LYSINE (L) HYDROCHLORIDE | 88.1 | 83.7 | | 77.2 |
| 1622 | THREONINE (L) | 87.7 | 84.9 | | 81.1 |
| 1623 | BUTOPYRONOXYL | 84.9 | 83.5 | | 79.6 |
| 1624 | METHIONINE (L) | 87.1 | 82.7 | | 75.0 |
| 1625 | FLUPIRTINE MALEATE | 83.0 | 85.4 | | 78.4 |
| 1626 | FUMARIC ACID | 84.4 | 82.7 | | 73.7 |
| 1627 | VALINE (L) | 82.0 | 81.5 | | 74.7 |
| 1628 | ISOLEUCINE (L) | 84.5 | 82.7 | | 73.7 |
| 1629 | PHENYLALANINE (L) HYDROCHLORIDE | 83.7 | 82.5 | | 75.5 |
| 1630 | ZONISAMIDE | 71.5 | 82.7 | | 73.3 |
| 1631 | LACTOSE MONOHYDRATE | 86.2 | 80.8 | | 75.5 |
| 1632 | SODIUM SUCCINATE | 87.1 | 82.7 | | 77.6 |
| 1633 | PRAXADINE HYDROCHLORIDE | 83.3 | 84.6 | | 74.3 |
| 1634 | CYSTINE | 85.3 | 84.2 | | 74.6 |
| 1635 | EBASTINE | 134.3 | 131.7 | | 91.8 |
| 1636 | CLONIXIN | 81.6 | 80.4 | | 78.8 |
| 1637 | CETILISAT | 85.8 | 81.6 | | 74.4 |
| 1638 | TOSUFLOXACIN TOLUENESULFONATE HY | 79.6 | 81.6 | | 74.1 |
| 1639 | METACRESOL | 88.5 | 86.9 | | 76.6 |
| 1640 | MALEIC ACID | 85.4 | 82.0 | | 73.6 |
| 1641 | SULFISOMIDINE | 87.6 | 86.1 | | 77.4 |
| 1642 | GESTODENE | 82.2 | 81.1 | | 75.7 |
| 1643 | BRIMONIDINE | 84.8 | 82.5 | | 77.7 |
| 1644 | ANAZOLENE SODIUM | 91.6 | 87.7 | | 79.0 |
| 1645 | GLYCINE | 87.4 | 83.9 | | 78.8 |
| 1646 | ALGINIC ACID [Mol Wt ~200,000; monom | 85.4 | 83.0 | | 76.8 |
| 1647 | TETRABENAZINE | 86.7 | 81.1 | | 74.3 |
| 1648 | TARTARIC ACID | 85.3 | 82.3 | | 72.3 |
| 1649 | ADEFOVIR DIPIVOXYL | 88.6 | 88.0 | | 72.9 |
| 1650 | DESOGESTREL | 83.1 | 79.7 | | 72.0 |

| | | | | | |
|---|---|---|---|---|---|
| 1651 | BENACTYZINE HYDROCHLORIDE | 86.6 | 83.5 | | 80.6 |
| 1652 | BENDAMUSTINE HYDROCHLORIDE | 87.8 | 84.2 | | 75.0 |
| 1653 | BENZYL NICOTINATE | 87.5 | 84.2 | | 74.1 |
| 1654 | GEMCITABINE HYDROCHLORIDE | 86.6 | 83.7 | | 75.9 |
| 1655 | D-(-)-FRUCTOSE | 88.6 | 80.7 | | 74.8 |
| 1656 | FULVESTRANT | 87.8 | 81.0 | | 73.7 |
| 1657 | CITRIC ACID | 88.4 | 84.5 | | 68.2 |
| 1658 | SAXAGLIPTIN | 84.8 | 82.7 | | 68.7 |
| 1659 | PEMETREXED | 86.4 | 84.2 | | 72.3 |
| 1660 | FROVATRIPTAN SUCCINATE | 86.7 | 84.2 | | 68.8 |
| 1661 | RILUZOLE | 110.0 | 100.7 | | 74.6 |
| 1662 | BUTENAFINE HYDROCHLORIDE | 157.5 | 114.1 | | 48.5 |
| 1663 | PROTAMINE SULFATE | 88.1 | 142.7 | | 72.0 |
| 1664 | FOMEPIZOLE | 86.5 | 79.4 | | 71.3 |
| 1665 | PERGOLIDE | 92.3 | 91.9 | | 74.2 |
| 1666 | PHYSOSTIGMINE | 84.9 | 82.3 | | 72.3 |
| 1667 | LINACLOTIDE (1 mg/ml) | 84.9 | 82.5 | | 70.0 |
| 1668 | SODIUM PHENYLACETATE | 86.0 | 80.9 | | 69.7 |
| 1669 | AMISULPRIDE | 85.7 | 82.7 | | 72.3 |
| 1670 | ANTIMYCIN A (A1 shown) | 95.9 | 88.0 | | 71.9 |
| 1671 | SERINE (L) | 84.3 | 81.6 | | 70.1 |
| 1672 | VALETHAMATE BROMIDE | 82.2 | 80.4 | | 71.0 |
| 1673 | TANDUTINIB | 87.6 | 84.4 | | 70.9 |
| 1674 | NILVADIPINE | 85.6 | 83.0 | | 71.6 |
| 1675 | EMPAGLIFLOZIN | 85.6 | 85.0 | | 71.6 |
| 1676 | ROLIPRAM | 85.1 | 83.9 | | 68.7 |
| 1677 | PROCARBAZINE HYDROCHLORIDE | 84.9 | 81.7 | | 72.0 |
| 1678 | ACETOHEXAMIDE | 85.0 | 82.5 | | 71.9 |
| 1679 | PROLINE (L) | 87.8 | 82.0 | | 71.6 |
| 1680 | PITAVASTATIN CALCIUM | 89.3 | 82.7 | | 72.6 |
| 1681 | ARTEMOTIL | 137.9 | 138.7 | | 97.9 |
| 1682 | ACENEURAMIC ACID | 85.3 | 79.9 | | 77.6 |
| 1683 | RUFLOXACIN HYDROCHLORIDE | 86.4 | 81.3 | | 83.0 |
| 1684 | ACEFYLLINE | 85.7 | 80.6 | | 73.3 |
| 1685 | ZANAMIVIR | 84.7 | 79.1 | | 74.3 |
| 1686 | PIRFENIDONE | 85.4 | 79.4 | | 76.4 |
| 1687 | CLINDAMYCIN PALMITATE HYDROCHLOR | 98.4 | 93.2 | | 80.7 |
| 1688 | METHYLPREDNISOLONE SODIUM SUCCIN | 86.2 | 81.8 | | 78.6 |
| 1689 | FLUROTHYL | 87.2 | 81.1 | | 78.0 |
| 1690 | CARBAZOCHROME | 86.6 | 85.6 | | 80.7 |
| 1691 | CAPECITABINE | 82.7 | 78.4 | | 74.7 |
| 1692 | GLIQUIDONE | 79.3 | 77.5 | | 75.8 |
| 1693 | DIDANOSINE | 88.6 | 81.0 | | 74.2 |
| 1694 | CLORSULON | 81.1 | 83.5 | | 74.2 |
| 1695 | RIVAROXABAN | 84.4 | 85.0 | | 76.4 |
| 1696 | BISOCTRIZOLE | | 81.8 | | 74.2 |
| 1697 | OZAGREL HYDROCHLORIDE | 87.1 | 82.1 | | 81.5 |
| 1698 | CEFTEZOLE | 83.7 | 81.6 | | 78.7 |
| 1699 | METHYLMETHANE SULFONATE | 83.5 | 80.4 | | 77.5 |
| 1700 | DICHLORVOS | 86.9 | 80.8 | | 77.3 |
| 1701 | ZALTOPROFEN | 82.5 | 80.8 | | 76.9 |
| 1702 | EQUILIN | 89.0 | 92.6 | | 78.9 |
| 1703 | TREHALOSE DIHYDRATE | 79.8 | 78.8 | | 73.8 |
| 1704 | 3-[3,4-DICHLOROPHENYL]1,1-DIMETHYLU | 72.7 | 69.2 | | 81.6 |
| 1705 | SPAGLUMIC ACID | 84.0 | 83.0 | | 76.3 |
| 1706 | LUMEFANTRINE | 101.2 | 94.9 | | 80.9 |
| 1707 | IVABRADINE HYDROCHLORIDE | 82.7 | 79.9 | | 76.8 |
| 1708 | PIRIBEDIL HYDROCHLORIDE | 82.0 | 80.5 | | 78.4 |
| 1709 | PRAZOSIN HYDROCHLORIDE | 76.4 | 83.2 | | 77.1 |
| 1710 | VESAMICOL HYDROCHLORIDE | 80.3 | 77.0 | | 78.7 |
| 1711 | SODIUM LACTATE | 83.8 | 79.4 | | 78.6 |
| 1712 | PIMOBENDAN | 88.1 | 84.4 | | 76.5 |
| 1713 | TIOTROPIUM BROMIDE | 84.2 | 78.5 | | 72.8 |
| 1714 | EFLORNITHINE HYDROCHLORIDE HYDRAT | 86.2 | 82.7 | | 71.8 |
| 1715 | EMEDASTINE DIFUMARATE | 89.7 | 82.3 | | 82.2 |
| 1716 | DROXYDOPA | 83.5 | 78.6 | | 80.9 |
| 1717 | PENTOBARBITAL | 80.4 | 80.3 | | 79.8 |
| 1718 | SODIUM THIOGLYCOLATE | 79.3 | 80.8 | | 84.2 |
| 1719 | DELAPRIL HYDROCHLORIDE | 82.0 | 80.8 | | 83.0 |
| 1720 | PYRIPROXYFEN | 79.6 | 78.3 | | 84.9 |
| 1721 | SORBITOL | 84.0 | 83.5 | | 76.3 |
| 1722 | Ro15-413 | 81.3 | 79.4 | | 76.4 |

| | | | | | |
|---|---|---|---|---|---|
| 1723 | DORZOLAMIDE | 79.1 | 82.0 | | 85.0 |
| 1724 | TERAZOSIN HYDROCHLORIDE | 80.6 | 84.2 | | 81.0 |
| 1725 | GLYBURIDE | 83.3 | 82.3 | | 72.3 |
| 1726 | CURCUMIN | 83.7 | 84.7 | | 87.1 |
| 1727 | IMATINIB | 81.1 | 82.3 | | 71.3 |
| 1728 | SULFLURAMID | 77.9 | 78.7 | | 71.3 |
| 1729 | CEFSULODIN SODIUM | 84.4 | 81.8 | | 85.7 |
| 1730 | GLYCYRRHIZIN | 82.3 | 81.8 | | 75.8 |
| 1731 | MOLSIDOMINE | 79.3 | 80.4 | | 74.8 |
| 1732 | IPROHEPTINE HYDROCHLORIDE | 79.9 | 80.4 | | 73.3 |
| 1733 | MOEXIPRIL HYDROCHLORIDE | 81.5 | 82.7 | | 75.8 |
| 1734 | CORTEXOLONE ACETATE | 81.2 | 82.7 | | 74.8 |
| 1735 | CETALKONIUM CHLORIDE | | | | |
| 1736 | DOXAPRAM HYDROCHLORIDE | 85.4 | 79.9 | | 83.0 |
| 1737 | CIANIDANOL [+-catechin] | 77.0 | 83.2 | | 76.5 |
| 1738 | DEXRAZOXANE | 77.3 | 83.0 | | 78.8 |
| 1739 | METHYL PARABEN | 76.6 | 83.2 | | 74.7 |
| 1740 | CEFCAPENE PIVOXIL HYDROCHLORIDE | 77.2 | 81.5 | | 75.6 |
| 1741 | LYNESTRENOL | 82.2 | 92.0 | | 86.1 |
| 1742 | ISOBUTYLMETHYLXANTHINE | 77.6 | 81.5 | | 75.3 |
| 1743 | CAPREOMYCIN SULFATE [4mM] | 78.1 | 84.0 | | 74.3 |
| 1744 | ROPINIROLE HYDROCHLORIDE | 79.4 | 87.6 | | 75.8 |
| 1745 | TEBUCONAZOLE | 183.5 | 163.8 | | 140.8 |
| 1746 | TIGECYCLINE | 79.9 | 83.5 | | 76.3 |
| 1747 | ETAZOLATE HYDROCHLORIDE | 79.8 | 81.2 | | 75.4 |
| 1748 | EPINASTINE HYDROCHLORIDE | 81.1 | 83.0 | | 82.1 |
| 1749 | RAFOXANIDE | 79.0 | 78.1 | | 79.9 |
| 1750 | 7-HYDROXYETHYLTHEOPHYLLINE | 78.1 | 81.3 | | 75.1 |
| 1751 | CHLORPHENIRAMINE MALEATE | 75.9 | 83.8 | | 78.3 |
| 1752 | AZAPERONE | 92.7 | 102.5 | | 89.5 |
| 1753 | ASTEMIZOLE | 78.2 | 85.9 | | 77.3 |
| 1754 | DETOMIDINE HYDROCHLORIDE | 79.4 | 80.3 | | 81.7 |
| 1755 | SYNEPHRINE TARTRATE | 80.3 | 82.5 | | 79.1 |
| 1756 | FENIPENTOL | 76.4 | 83.3 | | 78.0 |
| 1757 | EPLERENONE | 78.1 | 84.9 | | 76.0 |
| 1758 | GRANISETRON HYDROCHLORIDE | 76.2 | 79.6 | | 73.3 |
| 1759 | FLUNIXIN MEGLUMINE | 78.7 | 81.3 | | 74.3 |
| 1760 | EDOXABAN TOSYLATE HYDRATE | 80.2 | 87.0 | | 73.3 |
