## Supplement table 2 for "Fluphenazine, an antipsychotic compound, ameliorates Alzheimer’s disease by clearing amyloid beta accumulation in *C. elegans*"

| Sr.No. | Compound | Doubling time inflection (min) with 2.5ng/ml Rap | Doubling time inflection (min) compound only | fpr1 doubling time inflection (min) with 2.5ng/ml Rap | fpr1 doubling time inflection (min) Compound only |
|---|---|---|---|---|---|
| 1 | BITHIONOL | 280.6 | 81.7 | 130.7 | 105.3 |
| 2 | ALVERINE CITRATE | 101.8 | 76.5 | 88.0 | 99.9 |
| 3 | AMPHOTERICIN B | 193.6 | 103.2 | 96.7 | 98.0 |
| 4 | CHLOROXYLENOL | 116.4 | 78.6 | 99.6 | 101.3 |
| 5 | CHLORPROMAZINE HYDROCHLORIDE | 175.7 | 89.5 | 104.3 | 102.5 |
| 6 | AMITRIPTYLINE HYDROCHLORIDE | 111.7 | 77.4 | 91.2 | 85.4 |
| 7 | CLOTRIMAZOLE | 265.7 | 80.3 | 325.7 | 346.5 |
| 8 | DICYCLOMINE HYDROCHLORIDE | 101.2 | 77.9 | 100.4 | 103.9 |
| 9 | DYCLONINE HYDROCHLORIDE | 127.6 | 74.3 | 101.2 | 103.3 |
| 10 | CICLOPIROX OLAMINE | 115.5 | 81.4 | 98.9 | 102.0 |
| 11 | CLEMASTINE FUMARATE | 102.9 | 80.2 | 99.6 | 99.3 |
| 12 | DIBENZOTHIOPHENE | 122.4 | 80.6 | 102.5 | 104.5 |
| 13 | DISULFIRAM | 125.7 | 89.8 | 105.3 | 101.8 |
| 14 | ESTRONE | 109.4 | 80.4 | 99.5 | 109.4 |
| 15 | FLUOCINOLONE ACETONIDE | 189.3 | 99.4 | 98.9 | 111.3 |
| 16 | DORAMECTIN | 101.8 | 81 | 99.7 | 98.1 |
| 17 | ETHOPROPAZINE HYDROCHLORIDE | 138.7 | 84.8 | 103.6 | 101.4 |
| 18 | MECLIZINE HYDROCHLORIDE | 104.3 | 85.9 | 98.0 | 101.0 |
| 19 | HALOPERIDOL | 169.6 | 92.6 | 103.2 | 103.3 |
| 20 | PRIMAQUINE PHOSPHATE | 100.4 | 81.5 | 77.2 | 107.7 |
| 21 | POLYMYXIN B SULFATE | 273.7 | 80 | 93.9 | 98.7 |
| 22 | PARGYLINE HYDROCHLORIDE | 278.7 | 82 | 100.0 | 99.9 |
| 23 | PROCHLORPERAZINE EDISYLATE | 329.1 | 92 | 107.8 | 105.4 |
| 24 | NORTRIPTYLINE HYDROCHLORIDE | 114.9 | 80.5 | 98.6 | 96.4 |
| 25 | TRICHLORMETHIAZIDE | 299.1 | 112.3 | 100.3 | 98.8 |
| 26 | THIORIDAZINE HYDROCHLORIDE | 110 | 67.2 | 99.2 | 99.7 |
| 27 | FLUPHENAZINE HYDROCHLORIDE | 141.8 | 76.2 | 99.2 | 104.3 |
| 28 | ERYTHROMYCIN ESTOLATE | 106.9 | 83.1 | 93.9 | 91.3 |
| 29 | PERHEXILINE MALEATE | 414.3 | 98.2 | 98.1 | 97.8 |
| 30 | MEFLOQUINE HYDROCHLORIDE | 119.1 | 90.5 | 95.2 | 95.0 |
| 31 | DIXANTHOGEN | 111.2 | 86.4 | 104.9 | 107.3 |
| 32 | DOCUSATE SODIUM | 219.9 | 76.1 | 93.2 | 91.3 |
| 33 | LANSOPRAZOLE | 97.5 | 80.7 | 101.7 | 100.4 |
| 34 | PERPHENAZINE | 288.9 | 99.5 | 97.4 | 97.8 |
| 35 | TERFENADINE | 133.5 | 81.5 | 99.2 | 95.1 |
| 36 | PROPAFENONE HYDROCHLORIDE | 116.5 | 81.1 | 98.8 | 97.6 |
| 37 | CARVEDILOL | 106.1 | 75.2 | 96.6 | 101.3 |
| 38 | TEPOXALIN | 113.4 | 77.7 | 102.6 | 97.1 |
| 39 | FLUOXETINE HYDROCHLORIDE | 119 | 76.2 | 90.6 | 106.2 |
| 40 | CARVEDILOL PHOSPHATE | 101.5 | 76.1 | 98.1 | 101.8 |
| 41 | ACRISORCIN | 113.5 | 87 | 109.6 | 108.7 |
| 42 | CANDESARTAN CILEXTIL | 102.9 | 74.8 | 96.8 | 97.6 |
| 43 | ENROFLOXACIN | 115.4 | 75.5 | 97.4 | 99.3 |
| 44 | PROPOFOL | 93.19 | 72.3 | 103.5 | 105.3 |
| 45 | AMLODIPINE BESYLATE | 140.4 | 78.1 | 99.7 | 97.4 |
| 46 | SERTRALINE HYDROCHLORIDE | 210 | 82.2 | 108.6 | 104.3 |

| | | | | | |
|---|---|---|---|---|---|
| 47 | PAROXETINE HYDROCHLORIDE | 108.2 | 77.1 | 96.6 | 97.6 |
| 48 | AZELASTINE HYDROCHLORIDE | 116.93 | 92.5 | 98.2 | 100.4 |
| 49 | CLIOQUINOL | 117.4 | 80.3 | 118.1 | 114.1 |
| 50 | DRONEDARONE HYDROCHLORIDE | 306.8 | 79.4 | 100.4 | 100.4 |
| 51 | BENOXINATE HYDROCHLORIDE | 108.8 | 77.2 | 99.7 | 107.6 |
| 52 | BICALUTAMIDE | 102 | 72.2 | 96.0 | 98.9 |
| 53 | TERBINAFINE HYDROCHLORIDE | 141.3 | 63.8 | 94.8 | 91.2 |
| 54 | FLUORESCEIN | 102.4 | 88.1 | 120.6 | 122.0 |
| 55 | CHLOROPHYLLIDE Cu COMPLEX Na SALT | 107.1 | 80.6 | 107.9 | 108.2 |
| 56 | ARIPIPRAZOLE | 128 | 81.5 | 98.8 | 98.2 |
| 57 | DESONIDE | 100.3 | 75.4 | 98.4 | 99.5 |
| 58 | ATRACURIUM BESYLATE | 251.8 | 89.3 | 112.8 | 112.5 |
| 59 | CLOFAZIMINE | 255 | 84.3 | 120.0 | 112.7 |
| 60 | THIOPENTAL SODIUM | 102.7 | 68.3 | 90.6 | 95.6 |
| 61 | BENZYDAMINE HYDROCHLORIDE | 110.6 | 83.7 | 101.4 | 103.6 |
| 62 | BENAZEPRIL HYDROCHLORIDE | 123.1 | 80.2 | 104.4 | 102.0 |
| 63 | COLISTIN SULFATE | 291.8 | 80.5 | 71.7 | 99.5 |
| 64 | ENILCONAZOLE SULFATE | 223.4 | 75.3 | 118.2 | 146.1 |
| 65 | AMOXAPINE | 100.6 | 81.1 | 75.9 | 98.7 |
| 66 | BROMPERIDOL | 143.8 | 89.6 | 80.9 | 99.9 |
| 67 | VORTIOXETINE HYDROBROMIDE | 455.3 | 89 | | |
| 68 | IVERMECTIN | 101.9 | 71.9 | | |
| 69 | AMIODARONE HYDROCHLORIDE | 447 | 110.6 | | |
| 70 | DULOXETINE HYDROCHLORIDE | 101.5 | 70 | | |
| 71 | HEXETIDINE | 328.1 | 103.9 | | |
| 72 | PENTAMIDINE ISETHIONATE | 115.8 | 70.7 | | |
| 73 | PRASTERONE ACETATE | 109.7 | 82.4 | | |
| 74 | BROXYQUINOLINE | 192.6 | 83.5 | | |
| 75 | SODIUM TETRADECYL SULFATE | 293.6 | 89.3 | | |
| 76 | BENZYL ISOTHIOCYANATE | 186.4 | 88.2 | | |
| 77 | ARTEMISININ | 107.1 | 76.8 | | |
| 78 | CHLORMIDAZOLE | 114.7 | 83.9 | | |
| 79 | SULBENTINE | 150.2 | 107.3 | | |
| 80 | PRASTERONE | 103.6 | 84.4 | | |
| 81 | NAFTOPIDIL | 100.7 | 80.6 | | |
| 82 | METERGOLINE | 103.3 | 80.7 | | |
| 83 | NITROXOLINE | 334.1 | 115.8 | | |
| 84 | EBASTINE | 134.3 | 91.8 | | |
| 85 | RILUZOLE | 110 | 74.6 | | |
| 86 | BUTENAFINE HYDROCHLORIDE | 157.5 | 48.5 | | |
| 87 | ARTEMOTIL | 137.9 | 97.9 | | |
